# Structural mechanism of regioselectivity in an unusual bacterial acyl-CoA dehydrogenase

**DOI:** 10.1101/736256

**Authors:** Jacquelyn M. Blake-Hedges, Jose Henrique Pereira, Pablo Cruz-Morales, Mitchell G. Thompson, Jesus F. Barajas, Jeffrey Chen, Rohith N. Krishna, Leanne Jade G. Chan, Danika Nimlos, Catalina Alonso-Martinez, Edward E.K. Baidoo, Yan Chen, Jennifer W. Gin, Leonard Katz, Christopher J. Petzold, Paul D. Adams, Jay D. Keasling

**Affiliations:** Department of Chemistry, University of California, Berkeley, Berkeley, CA 94720; Joint BioEnergy Institute, Emeryville, CA, 94608; Biological Systems and Engineering Division, Lawrence Berkeley National Laboratory, Berkeley, CA 94720; Molecular Biophysics and Integrated Bioimaging, Lawrence Berkeley National Laboratory, Berkeley, CA 94720; Department of Plant and Microbial Biology, University of California, Berkeley, Berkeley, CA 94720; Department of Energy Agile BioFoundry, Emeryville, CA 94608, USA; QB3 Institute, University of California, Berkeley, Emeryville, CA, 94608; Department of Chemical & Biomolecular Engineering, Department of Bioengineering, University of California, Berkeley, Berkeley, CA 94720; Novo Nordisk Foundation Center for Biosustainability, Technical University Denmark, DK2970-Horsholm, Denmark; Center for Synthetic Biochemistry, Shenzhen Institutes for Advanced Technologies, Shenzhen, China

## Abstract

Terminal alkenes are easily derivatized, making them desirable functional group targets for polyketide synthase (PKS) engineering. However, they are rarely encountered in natural PKS systems. One mechanism for terminal alkene formation in PKSs is through the activity of an acyl-CoA dehydrogenase (ACAD). Herein, we use biochemical and structural analysis to understand the mechanism of terminal alkene formation catalyzed by an γ,δ-ACAD from the biosynthesis of the polyketide natural product FK506, TcsD. While TcsD is homologous to canonical α,β-ACADs, it acts regioselectively at the γ,δ-position and only on α,β-unsaturated substrates. Furthermore, this regioselectivity is controlled by a combination of bulky residues in the active site and a lateral shift in the positioning of the FAD cofactor within the enzyme. Substrate modeling suggests that TcsD utilizes a novel set of hydrogen bond donors for substrate activation and positioning, preventing dehydrogenation at the α,β position of substrates. From the structural and biochemical characterization of TcsD, key residues that contribute to regioselectivity and are unique to the protein family were determined and used to identify other putative γ,δ-ACADs that belong to diverse natural product biosynthetic gene clusters. These predictions are supported by the demonstration that a phylogenetically distant homolog of TcsD also regioselectively oxidizes α,β-unsaturated substrates. This work exemplifies a powerful approach to understand unique enzymatic reactions and will facilitate future enzyme discovery, inform enzyme engineering, and aid natural product characterization efforts.

## Introduction

Natural products often have complex chemical structures which can confer potent biological activity. Evolution selects for the diversification of these secondary metabolites, making natural product biosynthetic pathways rich resources for the discovery of both lead compounds for drug discovery and also enzymes with unique functions.^1–3^ Next-generation sequencing has led to the identification of numerous biosynthetic gene clusters (BGCs), but the pool of “orphan” BGCs (i.e. BGCs with no cognate natural product identified) remains largely untapped. Predicting natural product structures from primary DNA sequence is challenging, as the *in silico* functional annotation of enzymes within BGCs is limited. Amino acid sequence homology can suggest a general function, but without in depth structural and biochemical characterization of one or more members of an enzyme family, precise predictions of the final natural product structure are difficult. The identification of specificity-conferring motifs in polyketide synthase (PKS) acyltransferase (AT) and ketoreductase (KR) domains, for example, has allowed for more precise predictions of final polyketide natural product structure, including the identity of the alkyl substituents incorporated by (AT) and final stereochemical outcome (KR) of a given PKS module.^4^ While much effort has been dedicated to elucidating signature motifs within PKS domains, many PKS-associated enzymes that generate less common functional groups (such as non-canonical starter and extender units) are not as well annotated or understood. Better characterization of the enzymes implicated in the biosynthesis of unique and reactive moieties would facilitate the engineering of novel polyketides with applications in medicine, in agriculture, or as commodity chemicals.^1,5,6^

One particular moiety of interest for PKS engineering is the alkene, as alkenes could easily be chemically derivatized to introduce a multitude of other desirable functionalities into a natural product.^7–19^ Alkenes are often generated within a polyketide via the action of reductive domains, but they are sequestered within the polyketide backbone^20^ and are therefore less sterically accessible for chemical modification than terminal alkenes.^8,10-16,21-29^ Terminal alkenes, in addition to their innate reactivity, can also confer improved biological activity to polyketides that display biological activity, as is exemplified by the improved drug tolerability and efficacy of a synthetic epothilone analog sagopilone (anti-cancer activity).^30,31^ There are few known examples of polyketides that contain terminal alkenes, including FK506, haliangicin, curacin A, and tautomycetin (Figure 1A).^32-33,34^ In curacin A, a terminal alkene is generated through a unique off-loading mechanism involving a sulfotransferase (ST) and thioesterase domain ^25^. The tautomycetin terminal alkene is known to be formed after the chain has been released from the PKS but the process has yet to be fully characterized.^36,37^

**Figure 1.**
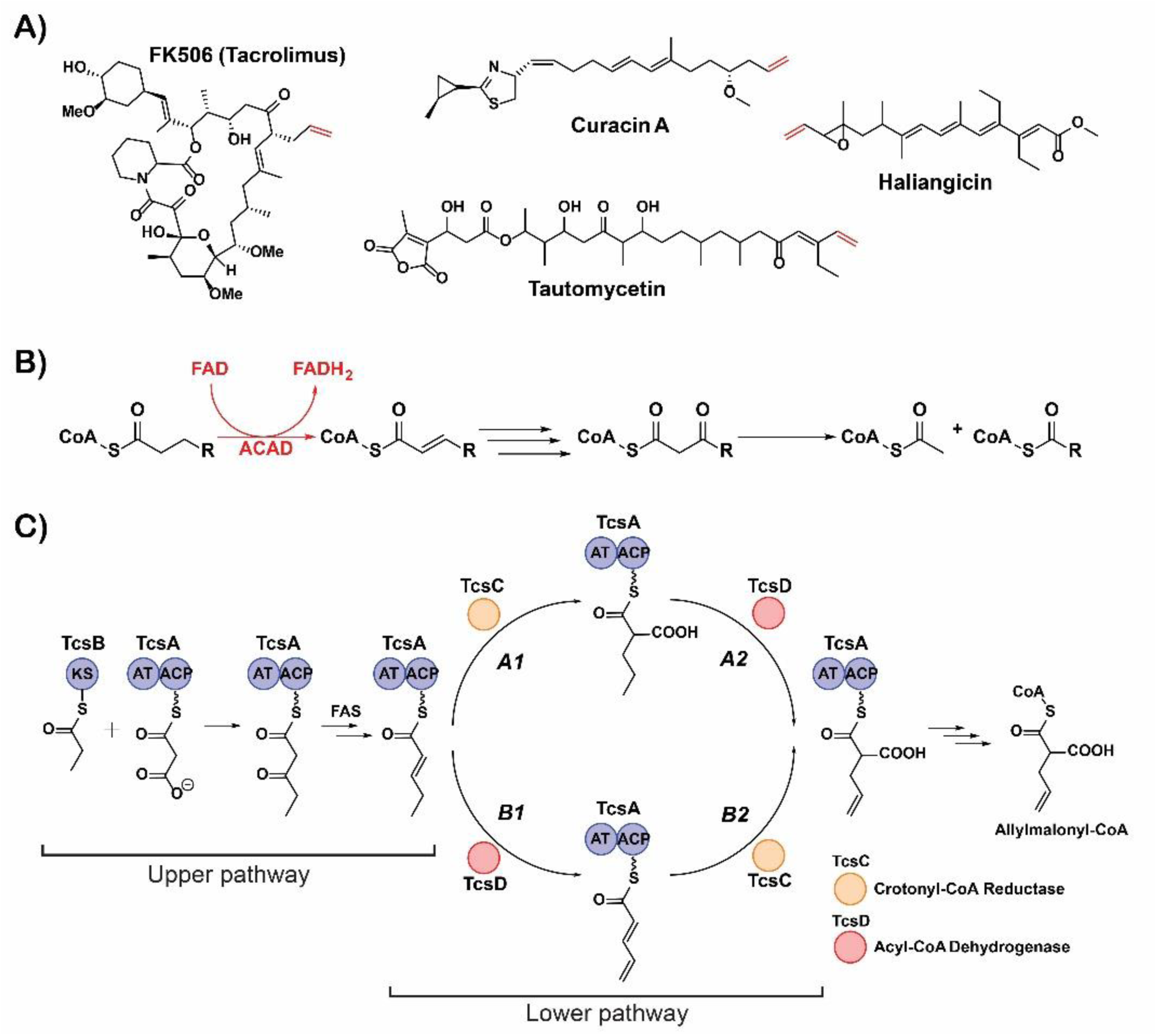
Terminal alkene-containing natural products and alkene formation by acyl-CoA dehydrogenases. **A)** Examples of terminal alkene-containing polyketide natural products. Terminal alkenes are highlighted in red. **B)** Process of fatty acid β-oxidation. The canonical activity of acyl-CoA dehydrogenases (ACADs) is the dehydrogenation of saturated fatty acyl-CoAs to form α,β-unsaturated products with concomitant reduction of FAD, highlighted in red. **C)** Proposed steps of allylmalonyl-CoA biosynthesis (adapted from reference 39). KS = ketosynthase domain, AT = acyltransferase domain, ACP = acyl carrier protein, FAS = fatty acid synthase.

A different mechanism for terminal alkene formation occurs via the action of an acyl-CoA dehydrogenase (ACAD) that oxidizes the γ,δ-position of a fatty acyl-CoA or acyl carrier protein (ACP), as observed in the haliangicin and FK506 pathways.^38,39^ The identification of other terminal alkene-forming ACADs is difficult, though, because of the enzymes’ homology to canonical ACADs. Thus, identifying important sequence motifs is critical to more accurate annotation. ACADs are oxidoreductase flavoenzymes well known for catalyzing the first oxidative step of fatty acid β-oxidation: the dehydrogenation of saturated fatty acyl-Coenzyme A (CoA) thioesters to form the corresponding α,β-unsaturated product (Figure 1B).^40^ The ACADs from the haliangicin and FK506 biosynthetic pathways, however, oxidize the γ,δ-position of a substrate. In order to facilitate better annotation of γ,δ-ACADs and, in turn, the identification of polyketide natural products that potentially contain terminal alkenes, a better functional characterization of γ,δ-ACADs is necessary. Herein, we report on our studies of the basis for the shift in regiochemistry of one of these unusual ACADs, TcsD, which forms the terminal alkene of the allylmalonyl-CoA extender unit in the biosynthesis of the polyketide FK506.^39^

The allylmalonyl-CoA biosynthetic pathway was initially proposed as a hybrid PKS-fatty acid synthase (FAS) pathway in which a free-standing ketosynthase, TcsB, first condenses a propionate group with a malonyl-CoA-derived extender unit selected by the AT domain of TcsA to form 3-oxo-pentanoyl-TcsA (Figure 1C, upper pathway).^39^ After reduction by the producing organism’s FAS, it was proposed that the final two enzymes TcsC (a crotonyl-CoA reductase) and TcsD (an ACAD) work interchangeably to convert 2-pentenoyl-TcsA to allylmalonyl-TcsA, with TcsD forming the unique terminal alkene moiety (lower pathway).

The sequence homology that TcsD shares with other ACADs suggests that it utilizes the same chemical mechanism to form the γ,δ-alkene of the allyl functional group. ACADs employ a catalytic base, typically a glutamate residue, to deprotonate the acidic α-proton of a fatty acyl-CoA substrate.^40-43^ The concomitant transfer of a hydride from the β-carbon of the substrate to FAD results in the formation of an α,β-unsaturated product and the reduced flavin, FADH_2_.^44^ The reaction is mediated by pK_a_ perturbations of both the catalytic glutamate residue and the substrate α-protons that occur within the active site of the enzyme. The pK_a_ of the glutamate is raised from ∼6 to ∼9 due to desolvation of the active site upon the binding of a hydrophobic substrate,^45-47^ while the substrate protons are activated via hydrogen bonds of the substrate thioester carbonyl group with the amide backbone of the glutamate and the 2’-hydroxyl group of FAD.^48-51^ While this mechanism of substrate activation is plausible for the γ-protons of a substrate such as 2-pentenoyl-TcsA (which contains an α,β-alkene that can propagate electronic effects from the thioester to the γ-carbon), the activation of propylmalonyl-TcsA and its conversion to allylmalonyl-TcsA (Figure 1C, pathway *A2*) is highly unlikely due to the aliphatic nature of the substrate.

Here we interrogate the activity of TcsD on both potential substrates and show that pathway *B1* (Figure 1C) is the only route of allylmalonyl-ACP formation. Additionally, we show that TcsD oxidizes only α,β-unsaturated substrates and is regioselective for the γ,δ position of these substrates. Further, we present a high resolution TcsD crystal structure and propose a structural mechanism by which it exhibits precise regiocontrol over this transformation. Combined structural and biochemical analyses of this enzyme revealed signature residues that contribute to its unique regioselectivity and facilitate the identification of homologs that display the same biochemical activity. A better understanding of this unique enzyme’s activity will not only inform the characterization of other homologs and their associated BGCs, but the insights gained herein can also aid future enzyme engineering efforts.

## Results and Discussion

### Biochemical activity of TcsD

In order to understand the mechanisms underlying TcsD activity, we initially sought to determine the native substrate(s) of the enzyme by biochemically interrogating both pathway *B*1 and *A*2 (Figure 1C). The substrates 2-pentenoyl-ACP (pathway *B1*) and propylmalonyl-ACP (pathway *A2*) were incubated with TcsD in the presence of the external electron acceptor ferrocenium hexafluorophosphate to facilitate enzyme turnover.^52^ Assays were analyzed using targeted LC-MS/MS to monitor for the characteristic phosphopantetheine ejection transition (Figure 2A).^53^ As expected, TcsD converted nearly all of the 2-pentenoyl substrate to the corresponding 2,4-pentadienoyl-TcsA product, with no activity detected in boiled TcsD controls (Figure 2B). However, TcsD was unable to convert propylmalonyl-TcsA to allylmalonyl-TcsA under identical assay conditions (Figure S1). We therefore concluded that the biosynthesis of allylmalonyl-CoA can only proceed through route *B*, in which TcsD first oxidizes 2-pentenoyl-TcsA to 2,4-pentadienoyl-TcsA (*B*1) and subsequently TcsC performs a reductive carboxylation to form allylmalonyl-TcsA (*B*2).

**Figure 2.**
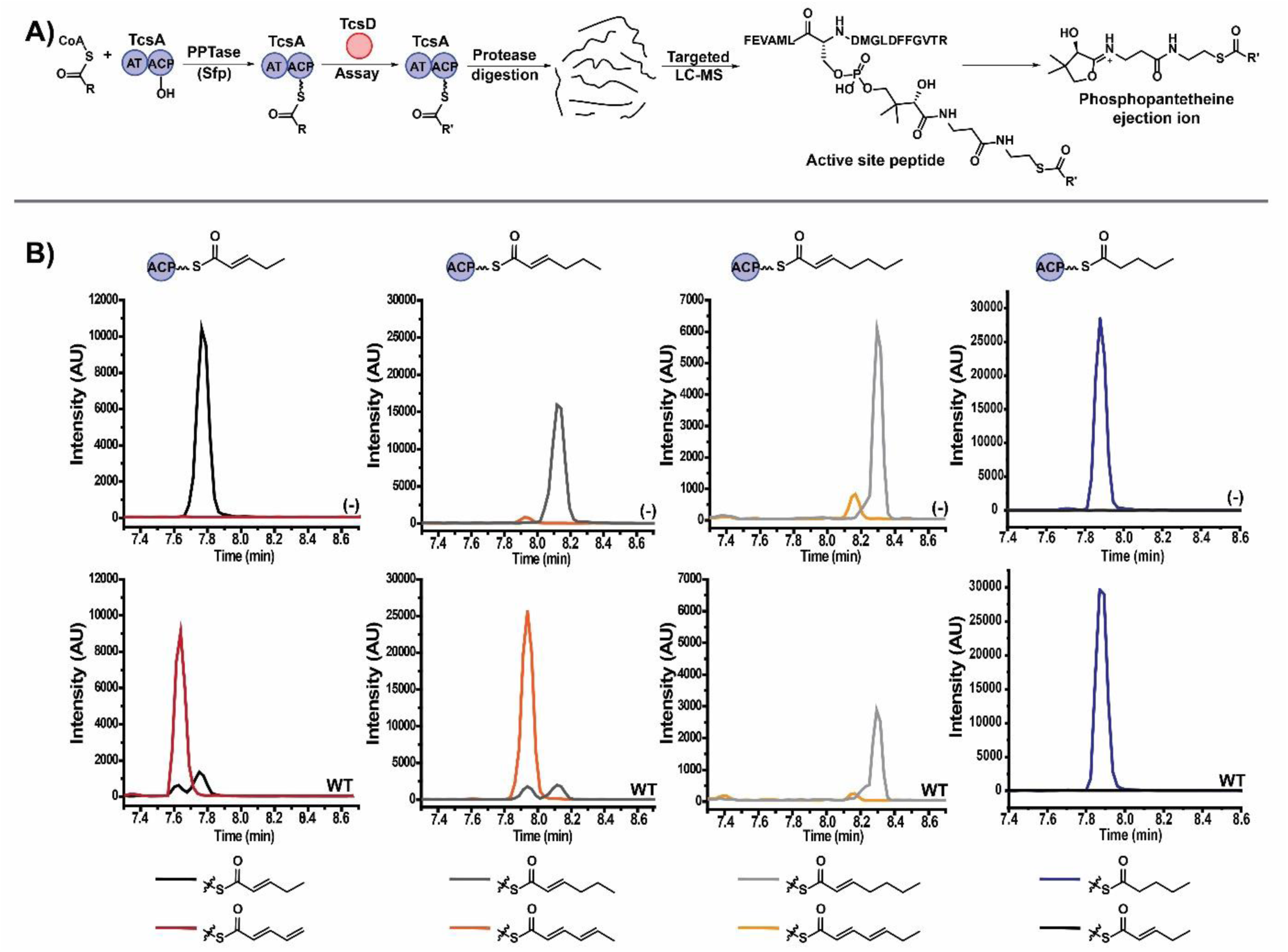
Biochemical activity of TcsD on ACP-bound substrates. **A)** Experimental design for phosphopantetheine ejection assays of TcsD activity on TcsA-bound substrates **B)** LC-MS/MS chromatograms of TcsD activity assays on various substrates. Substrates from left to right: 2-pentenoyl-TcsA, 2-hexenoyl-TcsA, 2-heptenoyl-TcsA, pentanoyl-TcsA. Top row: negative controls with boiled TcsD (-). Bottom row: assays with intact TcsD (WT).

Due to the noncanonical regiochemistry of the TcsD-mediated dehydrogenation of 2-pentenoyl-TcsA, we further investigated whether TcsD is regioselective for the γ,δ-position or if it simply oxidizes any appropriately activated substrate. It is known that some ACADs can abstract the acidic γ-proton of α,β-unsaturated substrates after dehydrogenation.^41,42,54^ Accordingly, it is plausible that TcsD is a promiscuous enzyme that dehydrogenates any substrate present on TcsA-ACP and that any specificity it exhibits *in vivo* arises from the sequestration of substrates on a protein (TcsA) instead of Coenzyme A. We therefore probed TcsD activity on a panel of α,β-unsaturated and fully saturated substrates. The enzyme was active on another α,β-unsaturated substrate, 2-hexenoyl-TcsA, but inactive on the seven carbon 2-heptenoyl-TcsA (Figure 2B). On the substrates butyryl-TcsA and pentanoyl-TcsA, where possible α,β-unsaturation could be expected, no oxidation activity was observed which indicated that TcsD acts regioselectively at the γ,δ-position (Figure 2B, S2 & S3).

### Crystal structure of TcsD

The strict substrate specificity of TcsD at the γ,δ-position suggested that there are structural elements within the enzyme’s active site that control regioselectivity. To test this hypothesis, we obtained a crystal structure of TcsD, which was solved to 1.75Å resolution. The TcsD structure displays many similarities to those of other ACADs, such as the conserved ACAD fold consisting of a tetrameric quaternary structure, which is further organized into two sets of homodimers (Figure S4). Each subunit contains a single FAD cofactor and is composed of 3 subdomains consisting of a set of N- and C-terminal alpha helix domains that surround a middle beta sheet domain. The FAD cofactor adopts an extended conformation and is situated in a pocket formed by the C-terminal alpha helix domain, the middle beta sheet domain, and the C-terminal domain of the adjacent subunit (Figure S4).

The active site of TcsD also shares a similar overall organization with α,β-ACADs. Like the structures of the human, rat, pig, and bacterial ACADs, the catalytic base of TcsD, Glu364, sits poised at the top of the active site pocket immediately above the fatty acyl binding chamber.^40,43,55-57^ The back side of the pocket that bounds the fatty acyl binding region is shaped by the residue immediately upstream of the catalytic glutamate, Ile363, and three residues from helix 5: Phe79, Leu83, and Leu86. The FAD cofactor sits at the bottom of the active site, positioned below Glu364(Figure 3A).

**Figure 3.**
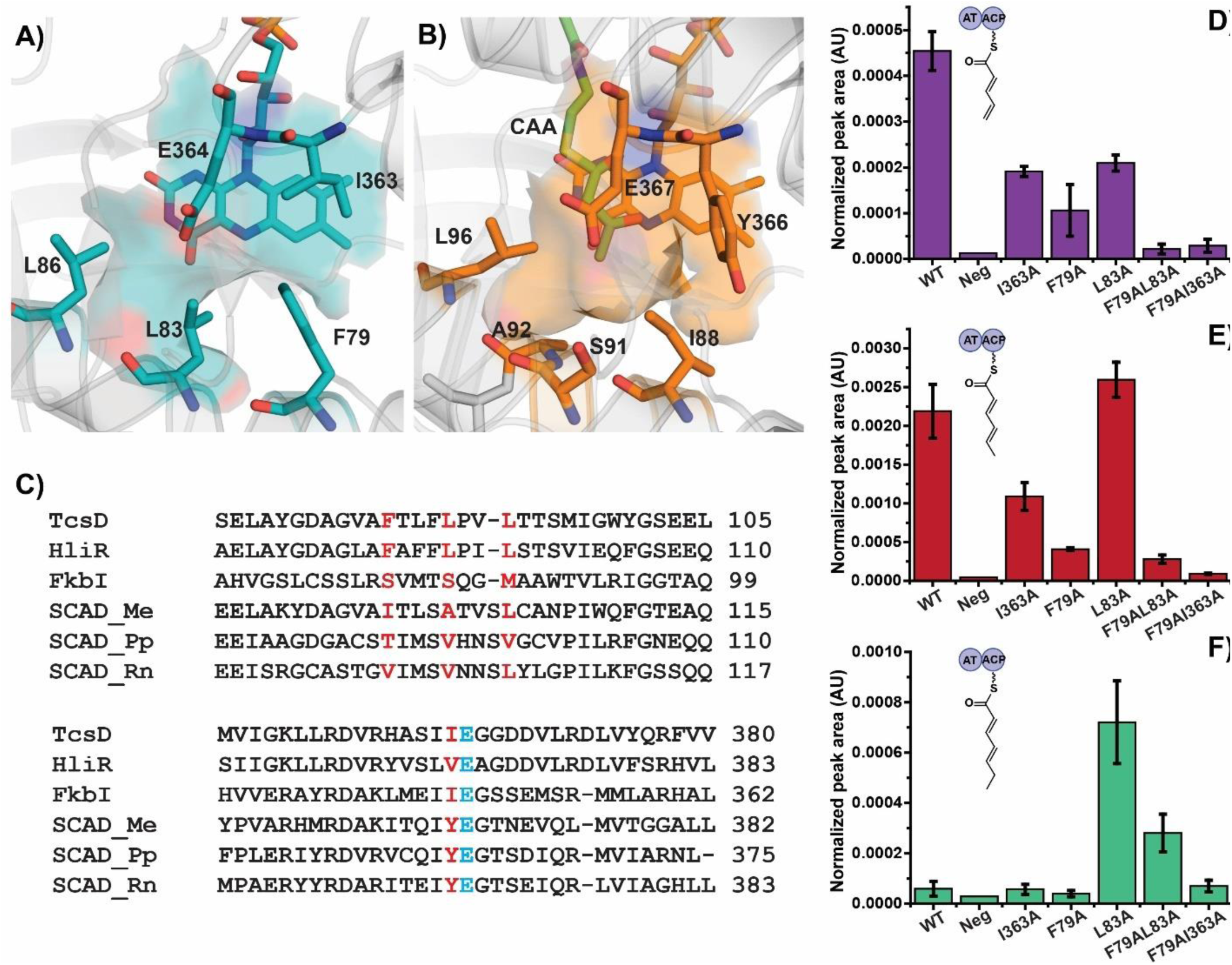
Unique active site features of TcsD and activity of active site mutants. **A)** Shape of the fatty acyl binding region of TcsD. **B)** Shape of the fatty acyl binding region of the *Megasphaera elsdenii* butyryl-CoA dehydrogenase (pdb entry 1buc), which was co-crystallized with acetoacetyl-CoA (CAA)^57^ **C)** Sequence alignment of TcsD with HliR,^38^ FkbI,^59^ *M. elsdenii* butyryl-CoA dehydrogenase (SCAD_Me, 1buc),^57^ *Pseudomonas putida* KT2440 short chain acyl-CoA dehydrogenase (SCAD_Pp),^58^ and rat short chain acyl-CoA dehydrogenase (SCAD_Rn, 1jqi).^56^ The residues in red correspond to residues that form the acyl-binding region of the enzyme active site (Phe79, Leu83, Leu86, and Ile363 in TcsD). The active site glutamate is highlighted in blue. **D), E), F)** TcsD mutant activity on 2-pentenoyl-TcsA* (**D**), 2-hexenoyl-TcsA* (**E**), and 2-heptenoyl-TcsA* (**F**) calculated as the LC-MS/MS peak area of the product normalized to a control peptide from TcsA (Normalized peak area, see Methods in SI). Data displayed are the average of three replicates. AU = arbitrary units *Note: TcsD activity on 5 carbon substrates cannot be compared to TcsD activity on 6 or 7 carbon substrates.

The isoalloxazine ring of FAD is anchored via conserved hydrogen bonds with residues Thr116, Gly118, Ser119, and Thr151, while the adenosine pyrophosphate portion of the molecule extends into a cavity formed between the loop that follows β-sheet 1 and helices 10 and 11 of the adjacent subunit; it is positioned through polar interactions with Ser125 and Glu337 of the same subunit and Met338, Gly340, and Gly341 of the adjacent subunit (Figure S5). Similar to other ACAD structures, an aromatic residue, Phe149, is positioned on the si face of the isoalloxazine ring. The FAD is positioned adjacent to the substrate binding cavity which is bound on the opposite side by several conserved residues from helix 7 and the loop following sheet 1, including Gly118, Ser119, Glu120, Ser125, Leu127, Leu224, and Leu228 which form the phosphopantetheine binding region.

Although similar in overall structure, TcsD possesses many features that distinguish it from canonical ACADs. The residues surrounding the TcsD active site entrance have diverged from the corresponding residues that are conserved in α,β-ACADs and contribute to the positioning of the adenosine portion of the coenzyme A substrate. TcsD does not possess the conserved Asp and Thr residues that form hydrogen bonds with the adenine base of Coenzyme A (Figure S6), nor the small helix between β-sheets 5 and 6 that often hydrogen bonds with the coenzyme A phosphate groups (Figure S7). The loop between β-sheets 5 and 6 in TcsD has instead been shortened. Additionally, a short helix, helix 9, is present in the TcsD structure in a position that would collide with the adenine base of Coenzyme A as it is positioned in α,β-ACAD structures. While helix 9 in the TcsD structure seems to interfere with Coenzyme A binding, it is also present in the TcsD homolog from the haliangicin pathway, HliR, suggesting that this helix might contribute to the regiochemical shift observed in both enzymes (Figure S8). TcsD has unique structural features that form the fatty acyl binding region of the active site pocket. In particular, Ile363 and the residues of helix 5 that line the back side of the active site pocket protrude further into the active site than other ACADs, forming a much shallower binding pocket (Figure 3A & B). More specifically, Phe79 and Leu83 protrude directly into the active site pointing toward the fatty acyl binding position. In addition, these two residues are bulkier than the residues forming the active site of canonical ACAD homologs, contributing to the reduced active site pocket depth seen in the TcsD structure. The difference in active site shape and depth is further illustrated by comparing TcsD with the structures and sequences of α,β-ACADs containing co-crystallized substrates. Alignment and superposition of the *Megasphaera elsdenii* butyryl-CoA dehydrogenase (BCAD) in complex with acetoacetyl-CoA^57^ with the TcsD active site shows that Phe79, Leu83, Leu86, Ile363 or a combination of these residues would sterically clash with substrates longer than acetoacetyl-CoA if they were to bind within the TcsD active site in a similar manner. The *M. elsdenii* BCAD and other short chain α,β-ACADs contain less bulky residues at the positions corresponding to Phe79, Leu83, and Ile363, but Leu86 is generally conserved in TcsD, HliR, and α,β-ACADs (Figure 3B & S8).^34,56,58-60^ In addition, the bulky residues Phe79 and Leu83 distinguish TcsD from FkbI, the hydroxymalonate semialdehyde dehydrogenase from the methoxymalonyl-ACP biosynthetic pathway of FK506 which has serines at both of these positions.^59^Given the unique structural characteristics of TcsD, we hypothesized that the positioning of the amino acids Phe79, Leu83, and Ile363 could control the regioselectivity of TcsD by preventing either proton abstraction by Glu364 or hydride transfer to FAD through steric repulsion. More specifically, these large residues would prohibit a five carbon substrate from entering the active site far enough so that Glu364 can access the protons bound to the substrate α-carbon or for the β-carbon to transfer a hydride to N5 of FAD. Rather, the substrate would be pushed towards the active site entrance, positioning Glu364 above the γ-carbon and the δ-carbon within an appropriate distance of N5 of FAD. As the substrates pentanoyl-ACP and 2-pentenoyl-ACP are similarly chemically activated (the pK_a_s of the α- and γ-protons are similar), the regioselectivity of TcsD would be controlled purely by steric interactions, not an electronic preference for one substrate.

### Biochemical activity of TcsD mutants

In order to verify whether the regioselectivity of TcsD is sterically controlled, a mutant of TcsD with a larger active site pocket that can accommodate a longer substrate is required. Specifically, TcsD mutants that act upon a substrate with two additional carbons should also be able to bind pentanoyl-ACP in a manner that properly positions the α- and β-carbons within an accessible distance of Glu364 and FAD, respectively, as the α-carbon is two carbons removed from the γ-carbon. The residues that surround the active site of TcsD were therefore selectively mutated to alanines in order to accommodate longer substrates. Individual alanine mutants of Phe79, Leu83, and Ile363 in addition to double mutants of neighboring residues (e.g. L83A/L86A, F79A/L83A, and F79A/I363A) were generated, and the dehydrogenation activity of mutants was then probed on a panel of α,β-unsaturated substrates. The active site mutants were generally less active on 2-pentenoyl-ACP than the wild type enzyme (Figure 3D). However, the I363A and L83A mutants displayed nearly equal or equivalent activity to the wild type on 2-hexenoyl-ACP (Figure 3E). Proteins containing the L83A mutation were also active on the 2-heptenoyl-ACP substrate (Figure 3F). This indicates that Leu83 controls the chain length of the substrate.

After demonstrating that the active site of the L83A mutant had been enlarged to accommodate two additional carbons, we next probed its activity on 4- and 5-carbon fully saturated substrates to test if the enzyme was now capable of dehydrogenating the α,β-position in the absence of the wild type steric interactions. We found that the mutant enzyme was still inactive on butyryl- and pentanoyl-ACP (data not shown), suggesting that the observed regioselectivity is not controlled exclusively by steric interactions with the fatty acyl tail of a substrate.

### Substrate modeling into TcsD active site

Given the retention of selectivity of the TcsD L83A mutant, we hypothesized that the regioselectivity of dehydrogenation could instead be controlled by intermolecular forces affecting the positioning of the thioester end of the substrate, which is determined by two hydrogen bonds (H-bonds) within the active site. We hypothesized that the canonical ACAD H-bonds should be conserved in TcsD as they are crucial to the chemical mechanism of the enzyme. To test this, we attempted to co-crystallize TcsD with 2-pentenoyl-CoA and 2-hexenoyl-CoA in order to show the substrate positioning within the active site; however, we were unable to observe density corresponding to either substrate in the TcsD active site pocket.

Instead, we computationally modeled the native TcsD substrate into the active site to understand which structural features might affect substrate binding. The positioning of the thioesters in the structures of several ACADs (which were co-crystallized with substrate analogs) was used to guide the placement of a substrate into the TcsD active site. More specifically, the structures of several homologs and TcsD were superimposed by aligning the peptide backbones of the three residue loop that contains the catalytic glutamate (e.g. Ile363, E364, and G365). This three residue loop was chosen as an anchor point because it is highly conserved and contains one of the H-bond donors that contributes to thioester positioning, so the relative positioning of structural elements within the ACAD active sites can be used to infer relative substrate positioning. Not only was this type of alignment useful for appropriately positioning modeled substrates, but it also allowed for a more precise determination of slight structural differences that were not as obvious when analyzing global structural alignments. In particular, it highlighted stark differences in the positioning of the FAD cofactor in TcsD relative to several homologs (Figure 4A and 4B).

**Figure 4.**
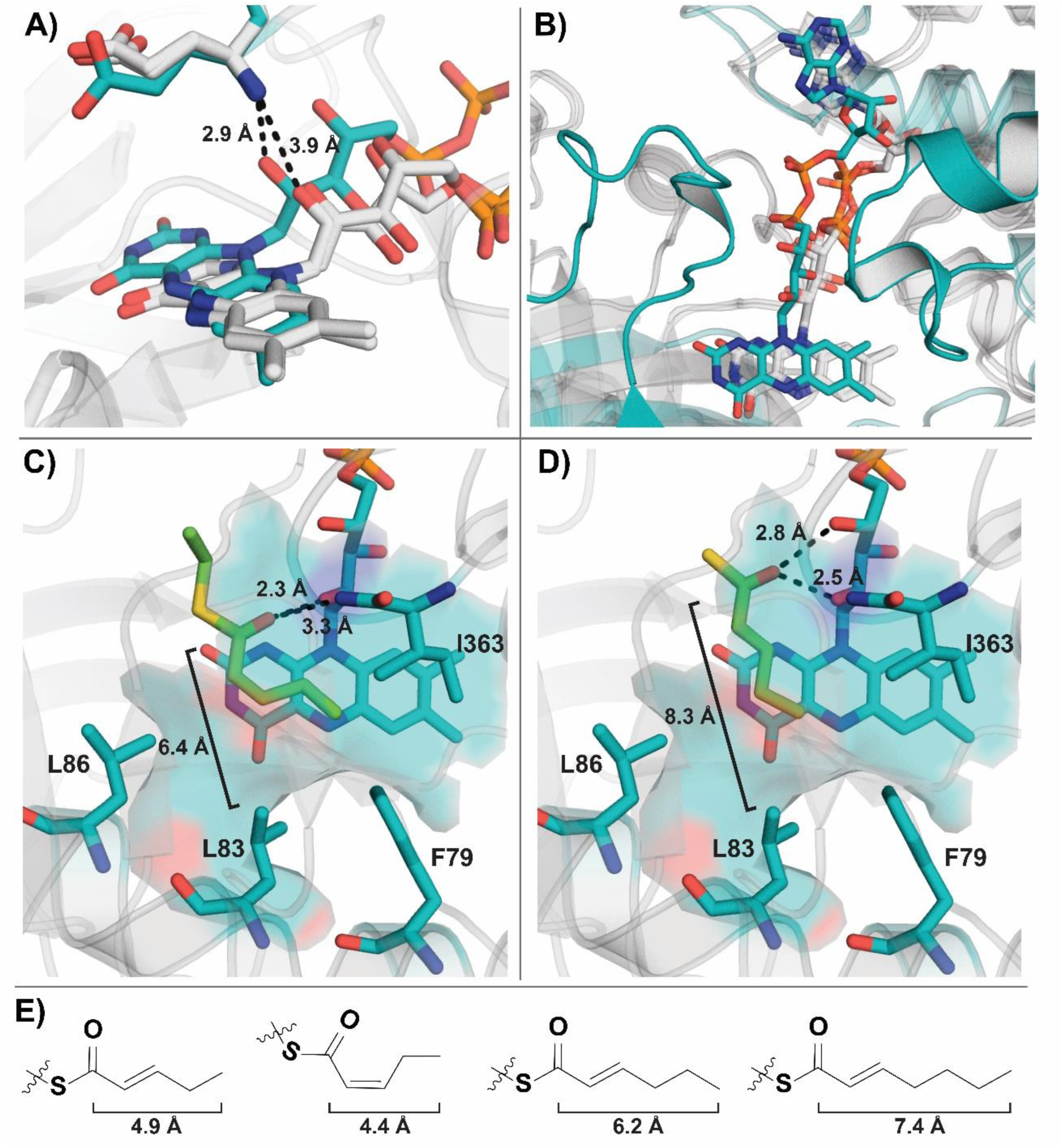
FAD shift and substrate modeling in TcsD active site. **A)** Shift in positioning of the 2’-OH group of FAD in TcsD (teal) relative to the active site glutamate and compared to the pig (*Sus scrofa*) medium chain ACAD in the apo (pdb code 3mdd)^43^ and substrate-bound (pdb code 1udy)^55^ forms (both white). Dotted lines represent distances between 2’-OH groups of FAD and amide nitrogens of active site glutamates. **B)** Lateral shift of FAD in TcsD (teal) with respect to the active site glutamate and compared to the pig (*Sus scrofa*) medium chain ACAD in the apo (pdb code 3mdd) and substrate-bound (pdb code 1udy) forms (both white) **C)** TcsD active site* containing a modeled 2-pentenoyl thioester group in the *cis* conformation. Dotted lines = distances between the substrate thioester carbonyl group and the 2’-OH of FAD and amide nitrogen of Glu364 **D)** TcsD active site* containing a modeled 2-pentenoyl thioester group in the *trans* conformation. Dotted lines = distances between the substrate thioester carbonyl group and the 2’-OH and 4’-OH of FAD **E)** Distance from carbonyl carbon to the tail carbon of various α,β-unsaturated fatty acyl substrates. *Note: Glu364 was omitted from C) and D) in order to visualize the hydrogen bonding interactions of the substrate with FAD.

In relation to the active site glutamate (Glu364), the FAD molecule bound within the TcsD active site is shifted laterally, moving it in the direction of the pantetheine-binding region of the enzyme (Figure 4A and B). This movement of the ribityl side chain of FAD is accompanied by a shift in α-helices 10 and 11 and the loop that follows β-sheet 1, which interact with the FAD through various H-bonds. The repositioning of α-helices 10 and 11 results in a change in the conformation of the FAD ribose ring due to H-bonding interactions of the 2’ and 3’-OH groups with the amide backbone of Met338 and of Glu337 and Gly339 of the adjacent subunit, respectively. The conformational change pushes the FAD phosphate groups towards the β-sheet domain of the enzyme. As a result of these structural changes, the positioning of the 2’-hydroxyl group of the FAD ribityl side chain is shifted closer to the nitrogen of the Glu364 amide bond (Figure 4A). We hypothesized that this would result in a concomitant shift in the positioning of the substrate thioester moiety, making TcsD act at the γ,δ-position instead ofat the canonical α,β-position of substrates.

In order to better understand how the change in FAD positioning affects substrate binding, we modeled a substrate analog consisting of a 2-pentenoyl thioester into the active site. Initially, we positioned the thioester carbonyl oxygen within hydrogen bonding distance of both the 2’-OH of the FAD and the nitrogen of the amide bond of Glu364 (Figure 4C). Anchoring the carbonyl group in this location would force the substrate alkene into a *cis* conformation, as this is the only conformation that places the δ-carbon in close enough proximity to N5 of FAD. However, while this conformation enforces the proper positioning of the substrate with respect to FAD, it places the δ-carbon too close to residues Phe79 and Ile363 which would result in unfavorable steric repulsion between the substrate and the amino acid side chains lining the active site. Moreover, with this substrate conformation there is no additional space within the active site to accommodate a sixth carbon on the substrate which conflicts with our biochemical data.

After eliminating the possibility of substrate binding in the *cis* conformation, a *trans* substrate was next modeled into the TcsD active site. When the substrate alkene bond adopts a *trans* conformation, its positioning is more consistent with the biochemical activity of the enzyme as it places the fatty acyl tail turned toward Leu83 and the δ-carbon in proximity to the N5 of FAD (Figure 4D). However, with the substrate in this extended position, H-bonding of the substrate with the canonical ACAD H-bond donors cannot occur without a steric clash between Leu83 and the fatty acyl tail of the substrate. We therefore modeled the carbonyl carbon of the substrate so that it H-bonds with the 2’ and 4’-OH groups of FAD, not with the canonical amide nitrogen of Glu364. Given the size dimensions of the active site and the biochemical data presented herein, this appears to be the most probable positioning of the substrate within the active site, suggesting that TcsD utilizes a novel H-bond donor pair to achieve its unique regioselectivity.

### Genome mining reveals previously unidentified γ,δ-ACADs

After identifying structural features that contribute to the unique regioselectivity of TcsD, we applied this mechanistic knowledge to find unidentified or misannotated γ,δ-ACADs among sequenced bacteria. In particular, we used the presence of bulky residues at positions corresponding to Phe79, Leu83, and Ile363 and helix 9 as requirements for the identification of γ,δ-ACADs. With these constraints, approximately 100 likely γ,δ-ACADs were identified using a combination of Hidden Markov Model (HMM), local sequence alignment (protein BLAST), and CORASON-BGC searches (Table S9 and S10). ^61-63^

Notably, both TcsD and HliR, the only γ,δ-ACADs with associated natural products, were re-identified in our search. Of the newly-identified enzymes one feature in particular was strongly conserved: the presence of helix 9, which we had identified as a unique feature in the TcsD structure (Figure 5A and B). The exact purpose of this loop remains unclear and will be the subject of further investigation. Additionally, we found that the presence of bulky residues at positions in the substrate acyl binding region were highly conserved across the family, with 98% of homologs displaying a Phe residue (2% Leu) and 96% displaying a Leu or Ile residue (4% Val) at the positions corresponding to Phe79 and Leu83 in TcsD (Figure 5C and S9). The residues immediately preceding the catalytic glutamate (i.e. residue Ile363 in TcsD) were all bulky aliphatic residues as well (87% Val, 11% Ile, 2% Phe). However, bulky residues at this position are not unique to the γ,δ-ACAD family as they are also observed in hydroxymalonate semialdehyde dehydrogenases such as FkbI.

**Figure 5.**
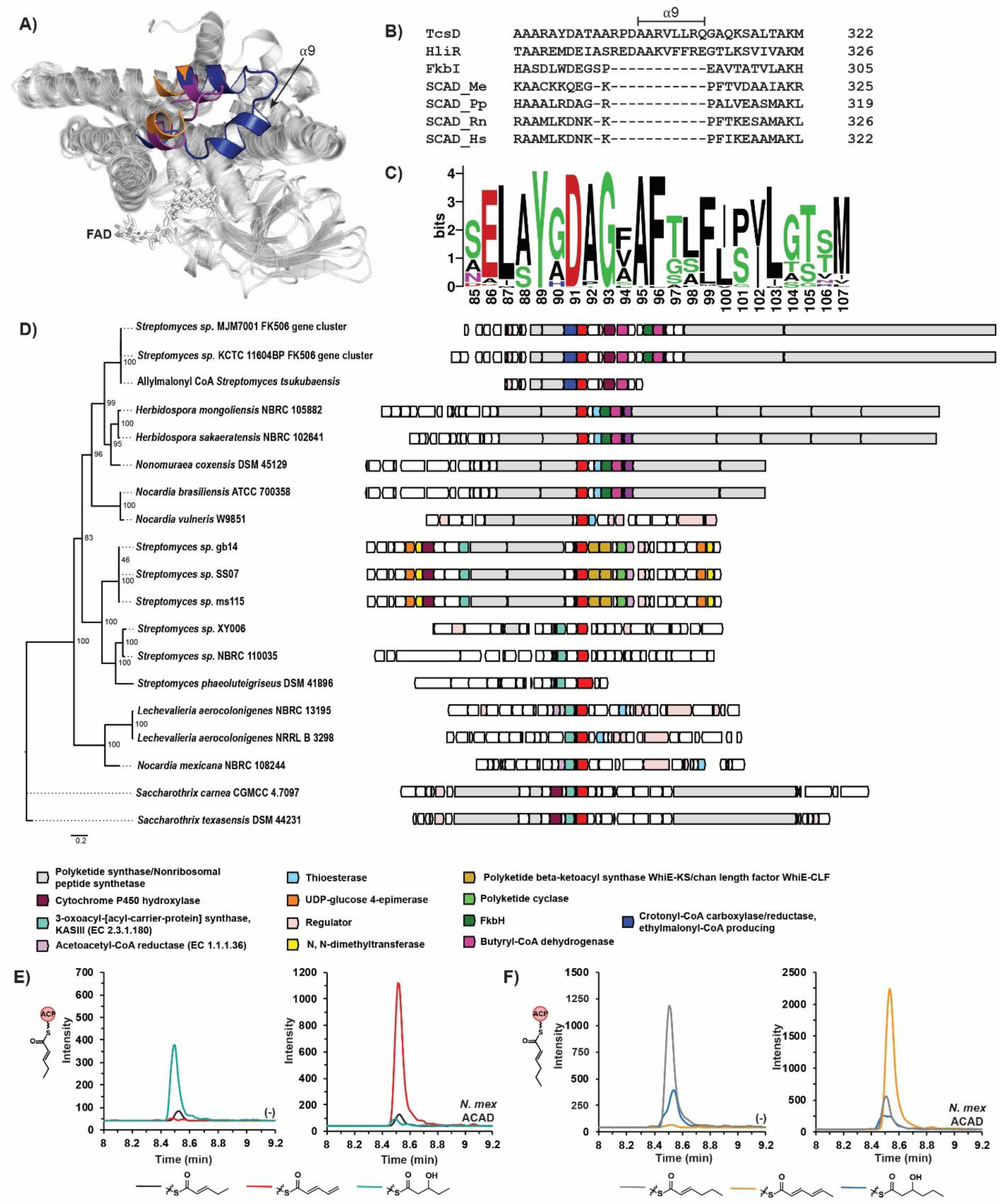
Genome mining and characterization of γ,δ-ACADs. **A)** Global overlay of TcsD with *M. elsdenii* butyryl-CoA dehydrogenase (BCAD_Me, 1buc)^57^ and *Streptomyces hygroscopicus* hydroxymalonate semialdehyde dehydrogenase, FkbI (1r2j)^59^ highlighting the insertion of helix 9 into γ,δ-ACAD structures. The colored regions are helix 9 of TcsD (blue, labeled α9) and the corresponding regions in BCAD_Me (purple) and FkbI (orange). FAD is depicted as sticks. **B)** Sequence alignment of TcsD with HliR,^38^ FkbI, *M. elsdenii* butyryl-CoA dehydrogenase (SCAD_Me, 1buc), *Pseudomonas putida* KT2440 short chain acyl-CoA dehydrogenase (SCAD_Pp),^59^ and rat short chain acyl-CoA dehydrogenase (SCAD_Rn, 1jqi).^56^ The sequence alignment highlights the presence of helix 9 in γ,δ-ACADs which is not encountered in other ACAD families. **C)** Amino acid position weight matrix showing the relative abundance of amino acids at each position within γ,δ-ACADs. Positions 96 and 100 correspond to the positions of Phe79 and Leu83 of TcsD, respectively. Logo was generated from a sequence alignment of all γ,δ-ACADs identified in this work. **D)** Genomic contexts and synteny of a representative set of γ,δ-ACADs identified in this work.TcsD homologs (putative γ,δ-ACADs) are shown in red and aligned. Other notable genes are annotated by color. **E)** Targeted LC-MS/MS chromatograms showing the activity of the *Nocardia mexicana* NBRC 108244 putative γ,δ-ACAD on 2-pentenoyl-ACP **F)** Targeted LC-MS/MS chromatograms showing the activity of the *Nocardia mexicana* NBRC 108244 putative γ,δ-ACAD on 2-hexenoyl-ACP. (-) is used to mark negative controls with boiled *N. mexicana* ACAD, while assays are marked as *N. mex* ACAD. Chromatograms representing the 2,3-enoyl-, 2,4-dienoyl-, and 3-hydroxy transitions are color-coded below each set of chromatograms. The 3-hydroxyacyl groups are normal degradation products of 2,3-enoyl-thioesters in aqueous environments where hydration of the alkene can occur.

Most of the TcsD homologs were associated with identifiable BGCs, although some were not located near any obvious secondary metabolite genes. The majority of the BGC-associated genes were located within type I PKS (T1PKS) clusters, but several were also identified within type II PKS (T2PKS) and nonribosomal peptide synthetase (NRPS) clusters (Figure 5D and S10). None of the newly-identified BGCs containing the putative γ,δ-ACADs have been experimentally characterized, but many show significant homology to known clusters (Table S10). Of the clusters that include putative γ,δ-ACADs, several contained proteins with homology to BGCs with known products, including the arsenopolyketides (T1PKS), oligomycin (T1PKS), E-837 (T1PKS), chlorothricin (T1PKS), butyrolactols (T1PKS), polyoxypeptin (T1PKS-NRPS), hedamycin (T2PKS), and erythrochelin (NRPS). Many of the homologous BGCs identified by antiSMASH have fatty acyl tails that could be potential substrates for γ,δ-ACADs in the uncharacterized clusters we identified.

Within the identified BGCs, analysis of the genomic contexts of the putative γ,δ-ACADs showed several patterns of syntenic genes (Figure 5D and S10). The syntenic genomic regions can be grouped based on the type of gene cluster (e.g., probable butyrolactol or arsenopolyketide BGC) within which the putative γ,δ-ACADs occur. The groups of genes that are found near γ,δ-ACADs can also be used to postulate the enzymes’ native substrates and even certain aspects of the molecular structure of the final natural product. The γ,δ-ACADs found in the erythrochelin-like gene clusters in *Saccharothrix sp.*, for example, are consistently clustered with a free-standing ketosynthase (KS) and acyl carrier protein (ACP) pair (Figure 5D). It is plausible that the KS domain forms a 5 carbon ACP-bound substrate upon which the γ,δ-ACAD can act after it is reduced to a 2-pentenoyl form. The genomic context of this and many of the other putative γ,δ-ACADs can be used to infer function and will inform future biosynthetic studies on the pathways and enzymes.

The presence of conserved residues and helix 9 in the putative γ,δ-ACADs is highly suggestive that they would exhibit the same activity as TcsD. To interrogate this hypothesis, we biochemically characterized a TcsD homolog. We chose the putative γ,δ-ACAD from *Nocardia mexicana* NBRC 108244 because it is located on a distant branch of the γ,δ-ACAD phylogenetic tree (Figure 5D and S10) and has low sequence identity relative to TcsD (50.7%). Though sharing only 51% sequence identity, the *N. mexicana* homolog (Nmex-ACAD) possesses the conserved Phe, Leu, and Ile residues that line the TcsD active site (Figure S11). It is located within a gene cluster that is predicted to encode a ladderane/butyrolactone-like natural product BGC (Figure S12). There are several related biosynthetic genes located immediately next to the Nmex-ACAD, including an acyl carrier protein, a ketosynthase, and a putative 3-oxoacyl-ACP reductase. Based on its genomic context we hypothesized that, like TcsD, the Nmex-ACAD may act on ACP-bound substrates. We therefore expressed and purified both the Nmex-ACAD and its neighboring acyl carrier protein (Nmex-ACP) and assayed Nmex-ACAD activity on the same panel of substrates. The Nmex-ACAD showed the same activity profile as TcsD, converting 2-pentenoyl-Nmex-ACP (Figure 5E) and 2-hexenoyl-Nmex-ACP (Figure 5F) to the corresponding dienoyl-ACP products, but it showed no activity on butyryl-, pentanoyl-, or 2-heptenoyl-ACP (Figure S13). While this data does not confirm that all the putative enzymes we have identified are γ,δ-ACADs, it strongly supports the prediction that these enzymes have the same activity as they are more closely phylogenetically related to TcsD than the Nmex-ACAD and share the conserved bulky residues that form the enzyme fatty acyl binding pocket.

### Conclusion

Terminal alkenes in polyketide natural products can be formed in several ways, including through the action of an acyl-CoA dehydrogenase. In this work we have described the biochemical and structural characterization of TcsD, the terminal alkene-forming γ,δ-ACAD from the biosynthesis of the allylmalonyl-CoA extender unit implicated in the biosynthesis of the polyketide FK506. We showed that TcsD acts on 2-pentenoyl-TcsA but not on propylmalonyl-TcsA, suggesting that the bottom half of the allylmalonyl-CoA pathway proceeds only through pathways *B1* and *B2*. TcsD only acts on 5- and 6-carbon α,β-unsaturated substrates and is regioselective for the γ,δ-position.

A crystal structure of TcsD revealed the unique features of the active site of the enzyme. Residues Phe79, Leu83, and Ile363 form a bulky wall in the substrate binding region of the enzyme, preventing the entrance of long fatty acyl substrates. Leu83 controls the chain length of the substrate. A TcsD L83A mutant acts on 2-heptenoyl-TcsA, but even with a larger active site pocket the mutant remains regioselective for the γ,δ-position of substrates. Closer analysis of the protein structure revealed that the enzyme regioselectivity is likely due to a shift in the positioning of the FAD cofactor. We show through substrate modeling that, because of the FAD shift and the dimensions of the TcsD active site, TcsD most likely employs a novel hydrogen bond donor pair (the 2’-OH and 4’-OH groups of FAD) to position and activate substrates. While Leu83 does not exclusively control regioselectivity, it contributes to regioselectivity by reducing the size of the active site.

The structural and biochemical conclusions from biochemically and structurally characterizing TcsD allowed us to determine key residues that define γ,δ-ACADs. Through HMM and local alignment searches, approximately 100 putative γ,δ-ACADs were identified in sequenced bacterial genomes. Nearly all of the homologs contained a Phe-Leu/Ile pair at the positions corresponding to Phe79-Leu83 in TcsD, respectively. The identification of other homologs also highlighted the conservation of helix 9, which appears to be a feature that is unique to the γ,δ-ACAD family. Most of the homologous enzymes were encountered in identifiable secondary metabolite BGCs, but some were notably located near no other canonical specialized metabolic enzymes. The synteny of genes located near the γ,δ-ACADs and the type of BGC to which they belong correlates strongly with their phylogenetic clustering and can be used to group the enzymes into several subfamilies. Finally, we showed that one of the TcsD homolog from *Nocardia mexicana*, which is phylogenetically one of the most distant enzymes from TcsD, also performs a regioselective dehydrogenation of the γ,δ-carbon of 2-pentenoyl- or 2-hexenoyl-ACP substrates, suggesting that this activity is conserved across the entire family.

This work exhibits how selective pressure causes enzymes to diverge not only at the amino acid sequence level, but also how it can result in significant shifts in protein structure to generate enzymes with divergent functions. Furthermore, it exemplifies how an understanding of the mechanisms employed by unique enzymes can be used as a means to refine the definitions of enzyme families and identify uncharacterized homologs. It will inform future efforts to characterize the identified homologs and the BGCs they reside within and can be used as a guide for the future discovery of natural products that contain terminal alkene handles.

## Acknowledgements

This work was performed as part of the DOE Joint BioEnergy Institute (https://www.jbei.org) supported by the U. S. Department of Energy, Office of Science, Office of Biological and Environmental Research, supported by the U.S. Department of Energy, Energy Efficiency and Renewable Energy, Bioenergy Technologies Office. The views and opinions of the authors expressed herein do not necessarily state or reflect those of the United States Government or any agency thereof. Neither the United States Government nor any agency thereof, nor any of their employees, makes any warranty, expressed or implied, or assumes any legal liability or responsibility for the accuracy, completeness, or usefulness of any information, apparatus, product, or process disclosed, or represents that its use would not infringe privately owned rights. The United States Government retains and the publisher, by accepting the article for publication, acknowledges that the United States Government retains a nonexclusive, paid-up, irrevocable, worldwide license to publish or reproduce the published form of this manuscript, or allow others to do so, for United States Government purposes. The Department of Energy will provide public access to these results of federally sponsored research in accordance with the DOE Public Access Plan (http://energy.gov/downloads/doe-public-access-plan). In addition, this work was supported by the Agile BioFoundry (https://agilebiofoundry.org), through contract DE-AC02-05CH11231 between Lawrence Berkeley National Laboratory and the U.S. Department of Energy. The Advanced Light Source is a Department of Energy Office of Science User Facility under Contract No. DE-AC02-05CH11231. The ALS-ENABLE beamlines are supported in part by the National Institutes of Health, National Institute of General Medical Sciences, grant P30 GM124169-01. J.M.B. was supported by the National Science Foundation Graduate Research Fellowship Program under Grant No. DGE 1106400.

## Notes

J.D.K. has financial interests in Amyris, Lygos, Demetrix, Napigen, Maple Bio, Ansa Biotechnologies, Berkeley Brewing Sciences, and Apertor Labs.

## Associated Content

### Supporting Information

Materials, methods, detailed experimental procedures, supplementary discussion, bioinformatic analysis, supplementary figures, crystallographic information.

### Structural data deposition

The atomic coordinates and structural factors of TcsD have been deposited in the Protein Data Bank, PDB ID code XXXX.

### LC-MS/MS data deposition

All targeted LC-MS/MS data from TcsD assays has been uploaded to Panorama Public^64^ and is publicly available at the following link: XXXX

