## Supplementary Info for "Structural mechanism of regioselectivity in an unusual bacterial acyl-CoA dehydrogenase"

Figure S10……………………………………………………………………………………………...44
Figure S11……………………………………………………………………………………………...46
Figure S12……………………………………………………………………………………………...47
Figure S13……………………………………………………………………………………………...48
Figure S14……………………………………………………………………………………………...49
Figure S15……………………………………………………………………………………………...50
Figure S16……………………………………………………………………………………………...51
Figure S17……………………………………………………………………………………………...52
Figure S18……………………………………………………………………………………………...53
Figure S19……………………………………………………………………………………………...54
Figure S20……………………………………………………………………………………………...55

**Data Availability**……………………………………………………………………………………56

**Supplementary References**..…………………………………………………………………………57

**Materials, Reagents, and Strains**

All strains used and generated in this work are listed in Table S2. Routine *E. coli* cultures were grown at 37°C in Luria-Bertani (LB) Miller medium (BD Biosciences, USA) and were supplemented with kanamycin (50 mg/L, Sigma Aldrich, USA), or carbenicillin (100mg/L, Sigma Aldrich, USA). *Pseudomonas putida* KT2440 was grown at 30°C in Luria-Bertani (LB) Miller medium (BD Biosciences, USA), and *Streptomyces tsukubaensis* NRRL 18488 was grown at 30°C in BD Bacto tryptic soy broth (TSB) medium (Fisher Scientific). All chemicals and reagents were purchased from Sigma Aldrich unless otherwise noted.

**DNA Manipulation**

All plasmids used and generated in this work are listed in Tables S1. The genomic DNA samples from *Pseudomonas putida* KT2440 were purified using a DNeasy Blood and Tissue Kit (Qiagen, USA). Plasmids and PCRs were routinely isolated using the Qiaprep Spin Miniprep kit (Qiagen, USA) and DNA Clean & Concentration Kit (Zymo Research, USA). All primers were purchased from Integrated DNA Technologies (IDT, Coralville, IA). Constructs were generated using restriction enzyme cloning (TcsD, TcsA wildtype coding sequences), Gibson assembly (TcsD mutants, PP2216), or Golden Gate assembly (*N. mex* ACAD and ACP) as previously described.[1,2]

**Construct Cloning: TcsD and TcsA**

Genomic DNA of *Streptomyces tsukubaensis* NRRL 18488 was prepared by resuspending cells grown in TSB medium in LC-grade DMSO. The coding sequences of TcsD and TcsA were amplified from this genomic DNA sample using Q5 Hotstart high-fidelity polymerase (New England Biolabs, USA) using primers which incorporated flanking NdeI and XhoI restriction sites. The genes were then ligated into a pET28a backbone which had been linearized with the FastDigest enzymes NdeI and XhoI (Thermo Scientific, USA), appending a C-terminal 6x histidine tag onto each. Targeted mutations were generated in TcsA and TcsD using Gibson cloning. Mutations were embedded in the overlap regions of the Gibson primers listed in Table S3**.**

**Construct Cloning: PP2216**

PP2216 was amplified from *Pseudomonas putida* KT2440 genomic DNA and cloned into a pBbE7a backbone with a C-terminal 6x histidine tag using Gibson assembly.

**Construct Cloning: *Nocardia mexicana* ACP and ACAD**

The coding sequences for the *Nocardia mexicana* NBRC 108244 putative γ,δ-acyl-CoA dehydrogenase (GenBank accession WP_068019852.1) and the neighboring acyl carrier protein (GenBank accession WP_068019854.1) were reverse translated and codon-optimized for *E. coli* using BOOST.[3] Genes were designed with flanking Golden Gate sites and purchased from Genscript (Piscataway, NJ, USA). The genes were then inserted into a pBbE7a backbone with a C-terminal 6x histidine tag using Golden Gate assembly. The sequences of the synthetic gene fragments are listed in Table S4.

**Protein Expression**

All proteins were expressed in *E. coli* BL21 (DE3) cells using the following protocol. An overnight culture of Luria Broth containing the appropriate antibiotic was used to inoculate 0.8L of Terrific Broth (EMD Millipore) in a 2L flask. Cells were grown at 37ºC to an OD of 0.6-0.8 then induced with 250 µM IPTG (Teknova, USA). Cells were grown for approximately 18 hours at 18ºC then pelleted and stored at -20°C until purification.

**Protein Purification for biochemical assays and substrate biosynthesis**

Frozen cell pellets were resuspended in lysis buffer that had been prechilled to 4°C. For TcsD, TcsD mutants, and the *Nocardia mexicana* ACAD, lysis buffer consisted of 50 mM Tris pH 9.0, 300 mM NaCl, 10 mM imidazole, 8% glycerol. For all other proteins (TcsAS98A, Sfp, PP2216, MatB T207G/M306I, *Nocardia mexicana* ACP), a lysis buffer composed of 50 mM sodium phosphate pH 7.2, 300 mM NaCl, 10 mM imidazole, 8% glycerol was used. After resuspension in lysis buffer, cells were sonicated on ice to lyse. Lysates were then centrifuged at 4°C at 40,000g for 30 minutes to pellet insolubles. The supernatants were then applied to Ni-NTA resin in an Econo-Pak column (Bio-Rad, USA) that had been pre-equilibrated with 10 column volumes of lysis buffer. The resin was washed with several column volumes of lysis buffer and subsequently with several column volumes of lysis buffer containing 50 mM imidazole. The proteins were then eluted using a stepwise gradient of lysis buffer containing 100 mM, 200 mM and 400 mM imidazole. Fractions were analyzed via SDS-PAGE, and fractions containing >90% pure protein were pooled and concentrated using Amicon Ultra 100 kDa (TcsD, TcsDmutants, and *Nocardia mexicana* ACAD), 3kDa (*Nocardia mexicana* ACP)10 kDa (all other proteins) Molecular Weight Cutoff (MWCO) centrifuge filters (Millipore). Proteins were transferred to storage buffer (lysis buffer containing no imidazole, 100 mM NaCl, and 8-10% glycerol) by buffer exchanging in the centrifuge filters and frozen in liquid nitrogen. Proteins were stored at -80ºC until assays.

**Protein purification for crystallography**

A frozen cell pellet of *E. coli* BL21 (DE3) expressing TcsD was purified using Ni-NTA affinity chromatography as described above. After concentration to a minimal volume, the protein was further purified using size exclusion chromatography. It was loaded onto a HiPrep 26/60 Sephacryl S-300 High Resolution column that had been pre-equilibrated with size exclusion buffer (50 mM Tris pH 9, 500 mM NaCl, 10% glycerol) at 4°C. The protein was eluted and concentrated again then dialyzed overnight at 4°C into crystallization buffer (50 mM Tris pH 9, 100 mM NaCl).

**Protein Crystallization and Structure Determination**

An initial crystallization screen was set up using a Phoenix robot (Art Robbins Instruments, Sunnyvale, CA) using the sparse matrix screening method.[4] The purified TcsD protein sample was concentrated to 9 mg/mL and crystallized at 25°C using the sitting drop method in 0.4 µL drops containing a 1:1 ratio of protein to crystallization solution: 0.1 M Sodium Citrate tribasic dihydrate, pH 5.0, and 10% PEG 6,000. Crystals were transferred to crystallization solution containing 20% glycerol prior to flash freezing in liquid nitrogen. X-ray diffraction data was collected at the Berkeley Center for Structural Biology on beamline 5.0.2 of the Advanced Light Source at the Lawrence Berkeley National Laboratory. The TcsD structure was determined by the molecular-replacement method with the program *PHASER*[5] using medium-chain acyl-CoA dehydrogenase from *Thermus thermophilus* (PDB ID: 1UKW) as the search model, which showed 32% sequence identity. Structure refinement was performed by *phenix.refine* program.[6] Manual rebuilding using COOT and the addition of water molecules allowed for construction of the final model.[7] The final models of the TcsD structure showed an R-work of 14.6% and R-free of 17.9% (Table S5). Root-mean-square deviations from ideal geometries for bond lengths, angles, and dihedrals were calculated with Phenix.[8] The overall stereochemical quality of the final models for tcsD was assessed using the MolProbity program.[9] The structural analysis was performed in COOT and PyMOL.[10]

**Synthesis of Pentanoyl-Coenzyme A.**

Pentanoyl-Coenzyme A was prepared using the anhydride method.[11] Approximately 0.02 mmol of Coenzyme A was added to a solution of saturated sodium bicarbonate in water. The solution was chilled to 0°C, then 1 mmol of valeric anhydride (Sigma Aldrich) was added. The reaction was allowed to proceed at 0°C with constant mixing. After approximately 6 hours, HCl was added to bring the pH to ~2. The solution was extracted twice with an equal volume of ethyl acetate, then the remaining aqueous solution was frozen until further purification. Pentanoyl-CoA was purified using an Agilent 1260 series preparatory HPLC system equipped with a 900µL sample loop and a UV detector coupled to a fraction collector. The product was isolated over an Agilent Prep-C18 column (21.2 x 150 mm, 5 µm pore size) using a mobile phase composed of 10 mM ammonium formate, pH 4.5 (solvent A) and methanol (solvent B) using the following method:

| Time (min) | %A | %B | Flow rate (mL/min) |
|---|---|---|---|
| 0 | 95 | 5 | 10 |
| 1 | 95 | 5 | 10 |
| 8 | 5 | 95 | 10 |
| 10 | 5 | 95 | 10 |
| 11 | 95 | 5 | 10 |
| 16 | 95 | 5 | 10 |

The fraction collector was programmed to collect all peaks that absorbed at 280 nm. Fractions containing pentanoyl-CoA were identified by direct infusion into an Applied Biosystems 4000 QTRAP mass spectrometer with the following mass spectrometer parameters: Q1 MS mode, scan range 100-1000 m/z, declustering potential: 70, entrance potential: 10, curtain gas: 10, IonSpray Voltage: 4800, Temperature: 300, Ion Source Gases: 40. The fractions containing the correct m/z for pentanoyl-Coenzyme A ([M+H]) were then combined and lyophilized, leaving behind pentanoyl-CoA as a white powder. The product was resuspended in LC grade water and quantified using the absorbance at 280 nm compared to a standard curve of hexanoyl-CoA in water (Sigma-Aldrich) measured on a Nanodrop ND-1000 Spectrophotometer. The identity of the Coenzyme A ester was additionally verified by targeted LC-MS/MS after loading the purified substrate onto TcsA S98A (methods described below).

**Trans-2-pentenoyl-, trans-2-hexenoyl-, and trans-2-heptenoyl-Coenzyme A.**

The 2,3-unsaturated CoA substrates were generated *in vitro* from pentanoyl-CoA (synthesized in this work), hexanoyl-CoA (Sigma Aldrich), or heptanoyl-CoA (CoALA Biosciences, USA) using the short chain acyl-CoA dehydrogenase PP2216.[12] Reactions contained the following components: 100mM sodium phosphate (pH 7.2), 500 µM Acyl-Coenzyme A, 1 mM ferrocenium hexafluorophosphate (FeHFP, Sigma Aldrich), 225 µM PP2216 in a final volume of 1 mL. FeHFP was always prepared fresh by dissolving solid FeHFP in 10mM HCl and sonicating in a water bath to dissolve solid particles.[13] Reactions were incubated at 25°C for 2 hours (2-

pentenoyl-CoA) or 6 hours (2-hexenoyl and 2-heptenoyl-CoA) then filtered through 3kDa MWCO Amicon microcentrifuge filters (Sigma Aldrich) to remove the remaining PP2216 protein. Samples were then aliquoted and lyophilized. After lyophilization, samples were stored at -20°C and resuspended in HPLC water (Honeywell) at a concentration of 4mM (assuming maximum theoretical yield) immediately prior to use.

**Allylmalonyl- and propylmalonyl-Coenzyme A.**

Allylmalonyl-CoA and propylmalonyl-CoA were biosynthesized from the corresponding malonic acids using an engineered mutant of the *Streptomyces coelicolor* malonyl-Coenzyme A transacylase MatB, MatB T207G/M306I.[14] Reactions contained the following components: 100 mM sodium phosphate (pH 7.2), 2 mM $MgCl_2$, 8 mM ATP, 4 mM Coenzyme A, 4 mM diacid, 100 µM MatB T207G/M306I in a final volume of 0.5 mL. Prior to addition to assays, allylmalonic acid (Sigma Aldrich) and propylmalonic acid (VWR, BeanTown Chemical) were dissolved in 50 mM sodium phosphate buffer, pH 7.2. Reactions were incubated for 3 hours the filtered through 3kDa MWCO Amicon microcentrifuge filters (Sigma Aldrich) to remove the remaining MatB protein. Mixtures were aliquoted and frozen at -20°C until use in TcsD assays.

**Enzyme Assays: TcsD and mutants**

Assays of TcsD activity on TcsAS98A-bound substrates were carried out as follows: First, purified TcsAS98A was loaded with acyl-Coenzyme A substrates via the action of the promiscuous phosphopantetheinyl transferase Sfp.[15] Loading reactions (150 µL) contained 60 µM TcsAS98A, 10 µM Sfp, 250 µM acyl-Coenzyme A, 20 mM $MgCl_2$, and 50 mM sodium phosphate, pH 7.2. Reactions were initiated by the addition of Sfp and allowed to proceed at 37°C for 30 minutes. The loaded protein was then immediately used for TcsD assays, each of which contained: 45 µL of loading reaction, 0.5 mM FeHFP, and 2 µM TcsD mutant (or boiled TcsD mutant). FeHFP was prepared as described above. Reactions were allowed to proceed for 16 hours then quenched with and equal volume of HPLC methanol (Sigma Aldrich) and either stored at -20°C or immediately processed for LC-MS analysis.

**Enzyme Assays: *Nocardia mexicana* ACAD**

The Nocardia mexicana ACP was loaded with butyrl-, pentanoyl-, 2-pentenoyl-, 2-hexenoyl, or 2-heptenoyl-CoA using Sfp.[15] Loading reactions (120 µL) consisted of 50 µM *N. mexicana* ACP, 10 µM Sfp, 250 µM acyl-Coenzyme A, 20 mM $MgCl_2$, and 40 mM Tris buffer, pH 8. Reactions were initiated by the addition of Sfp and allowed to proceed at 37°C for 1 hour. The loading reactions were then used for *N. mexicana* ACAD assays which consisted of: 40 µL of loading reaction, 0.5 mM FeHFP, and 2 µM *N. mexicana* ACAD. Reactions were allowed to proceed at 37°C for 16 hours then quenched with HPLC methanol and immediately processed for LC-MS analysis.

**Sample preparation for LC-MS/MS analysis of TcsD and Nmex-ACAD assays**

Samples were prepared for proteomics analysis by chloroform/methanol extraction as previously reported.[16] Briefly, 150 µL of LC-grade methanol and 50 µL of LC-grade chloroform were sequentially added to the methanol-quenched samples, and samples were vortexed after each addition. Then, 150 µL LC-grade water was added, samples were vortexed to mix, and they were subsequently centrifuged at 21,000g for 1 minute to promote phase separation. After removal and disposal of the top (aqueous) layer, another 150 µL of LC methanol was added, samples were vortexed, then they were centrifuged again at 21,000g for 2 minutes. After removal of the supernatant, samples were allowed to dry for 5 minutes in a fume hood. Then, they were resuspended in freshly-prepared 100 mM ammonium bicarbonate buffer (*N. mexicana* ACAD assays) or 100 mM ammonium bicarbonate containing 20% LC-grade DMSO (TcsD assays). Protein concentrations were analyzed via the DC Assay (Bio-Rad). After quantification, samples were digested with an appropriate protease. Samples digested with only trypsin (Sigma-Aldrich) or only chymotrypsin (Promega) were digested at a ratio of 1:50 w/w protease:protein sample at 37°C for at least 6 hours. For Asp-N/trypsin dual digestions (TcsD and TcsD mutant assays), proteins were digested with 1:25 w/w Asp-N (New England Biolabs) in the presence of 0.75 mM $ZnSO_4$ for 2-4 hours at 37°C then immediately digested again with 1:10 w/w trypsin for 1 hour at 37 $^{o}$C. The *N. mexicana* ACAD assays were digested with Glu-C (Promega) at a ratio of 1:50 protease:protein sample at 37°C for 6 hours. After digestion, the samples were either frozen for future analysis or directly analyzed via LC-MS/MS.[17]

**Shotgun LC-MS/MS Analysis of TcsA-bound substrates**

Samples prepared for shotgun proteomic analysis were analyzed using Agilent 6550 iFunnel Q-TOF mass spectrometer (Agilent Technologies, Santa Clara, CA) coupled to an Agilent 1290 UHPLC system. Twenty (20) µg of peptides were separated on a Sigma–Aldrich Ascentis Peptides ES-C18 column (2.1 mm × 100 mm, 2.7 µm particle size, operated at 60°C) at a 0.400 mL/min flow rate and eluted with the following gradient: initial condition was 95% solvent A (0.1% formic acid) and 5% solvent B (99.9% acetonitrile, 0.1% formic acid). Solvent B was increased to 35% over 5.5 min, and then increased to 80% over 1min, and held for 3.5 min at a flow rate of 0.6 mL/min, followed by a ramp back down to 5% B over 0.5 min where it was held for 2 min to re-equilibrate the column to original conditions. Peptides were introduced to the mass spectrometer from the LC by using a Jet Stream source (Agilent Technologies) operating in positive-ion mode (3,500 V). Source parameters employed gas temp (250°C), drying gas (14 L/min), nebulizer (35 psig), sheath gas temp (250°C), sheath gas flow (11 L/min), VCap (3,500 V), fragmentor (180 V), OCT 1 RF Vpp (750 V). The data were acquired with Agilent MassHunter Workstation Software, LC/MS Data Acquisition B.06.01 operating in Auto MS/MS mode whereby the 20 most intense ions (charge states, 2–5) within 300–1,400 m/z mass range above a threshold of 1,500 counts were selected for MS/MS analysis. MS/MS spectra (100–1,700 m/z) were collected with the quadrupole set to “Medium” resolution and were acquired until 45,000 total counts were collected or for a maximum accumulation time of 333 ms. Former parent ions were excluded for 0.1 min following MS/MS acquisition. The acquired data were exported as mgf files and searched against a customized protein database with Mascot search engine version 2.3.02 (Matrix Science).

**Targeted LC-MS/MS Analysis**

Samples were analyzed using an Agilent 1290 Infinity II liquid chromatography system coupled to an Agilent 6460 QQQ mass spectrometer (Agilent Technologies, Santa Clara, CA). Peptide samples (10 µg) were separated on an Ascentis Express Peptide ES-C18 column (2.7 µm particle size, 160 Å pore size, 50 x 2.1mm) fitted with a guard column (5 mm x 2.1 mm, Sigma Aldrich). The column was heated to 60°C. The mobile phase consisted of 0.1% formic acid in $H_2O$ (A) and 0.1% formic acid in acetonitrile (B). Peptides were eluted via the following gradient method:

| Time (min) | %A | %B | Flow rate (mL/min) |
|---|---|---|---|
| 0 | 95 | 5 | 0.4 |
| 2.2 | 95 | 5 | 0.4 |
| 7.7 | 65 | 35 | 0.4 |
| 8 | 20 | 80 | 0.4 |
| 10 | 20 | 80 | 0.4 |
| 10.5 | 95 | 5 | 0.4 |
| 12 | 95 | 5 | 0.4 |
| 15 | 95 | 5 | 0.4 |

Peptides were ionized using an Agilent Jet Stream ESI source operating in positive-ion mode with the following source parameters: Gas Temperature = 250°C, Gas Flow = 13 L/min, Nebulizer Pressure = 35 psi, Sheath Gas Temperature = 250°C, Sheath Gas Flow = 11 L/min, and Capillary Voltage = 3,500 V. Dwell times were set to 10ms. Data were acquired using Agilent MassHunter Data Acquisition (Version B.08.02, Build 8.2.8260.0). The mass spectrometry method was built using the Skyline targeted mass spectrometry environment (version 4.2.0 19072).[18] Collision energies were predicted by Skyline with the exception of phosphopantetheine-bearing peptides for which the collision energy was set to 30eV. The specific transitions monitored and the collision energies for each are listed in Tables S7 and S8.

**Targeted LC-MS/MS Data Availability**

Raw data were imported into Skyline and are available on the LC-MS data sharing platform Panorama Public at the following link: XXXX[19] Sample names for TcsD assays are labeled as "protein-substrate-replicate." For example, replicate one of the assay of wild type TcsD on pentanoyl-ACP is labeled "WT-val-01." Substrate abbreviations are as follows: val = pentanoyl-ACP, but = butyryl-ACP, 2pent = 2-pentenoyl-ACP, 2hex = 2-hexenoyl-ACP, 2hept = 2-heptenoyl-ACP, allylmal = allylmalonyl-ACP, propmal = propylmalonyl-ACP. The data for TcsD wild type assays on 2-pentenoyl-ACP, 2-hexenoyl-ACP, and 2-heptenoyl-ACP is located in the file named "WT assays_set 1_pub data.skyd." The data for TcsD wild type assays on propylmalonyl-ACP is located in the file named "Malonate assays_pub data.skyd." The data for TcsD wild type assays on butyryl-

ACP and pentanoyl-ACP and the data for all mutant TcsD proteins on all substrates is located in the file named "Mutants assays_WT-but-val_pub data.skyd."

Sample names for *N. mexicana* ACAD assays are labeled as "substrate_replicate," and substrate abbreviations are the same as for TcsD assays. The data for butyryl-, pentanoyl-, and 2-pentenoyl-ACP are located in the file named "NmexACP_C4_C5_pub data.skyd," and data for 2-hexenoyl- and 2-heptenoyl-ACP are located in the file named "NmexACP_C6_C7_pub data.skyd."

**Targeted LC-MS/MS Data Analysis: TcsD assays**

After importing data, chromatograms were manually curated. Correct peaks were identified by comparison of their retention times to control peaks (i.e. the 2-pentenoyl-ACP active site peptide should elute before the pentanoyl-ACP active site peptide). For phosphopantetheine ejection peaks, only the transition derived from the peptide "DLGVDSLAMTELQAHALQR" was used for analysis and quantification. Peaks that were not correctly automatically selected were manually selected in Skyline and integrated using the built in Skyline integration function. For negative controls, an area corresponding to the same retention time as the peak of interest was integrated. After integrating, all phosphopantetheine ejection ion peak areas were normalized to an internal control peptide from the within the TcsA protein (a peptide that does not contain catalytic residues) to account for differences in the amount of protein injected. The reported values in Figure 3 were averages of three normalized replicate peak areas. Error bars represent standard deviations of each set of replicates.

**Targeted LC-MS/MS Data Analysis: *N. mexicana* ACAD assays**

After importing data into Skyline, chromatograms were manually curated (including manual integration where necessary) as described above for TcsD assays. Plots in Figures 5E and 5F and Figure S13 correspond to the LC-MS/MS chromatograms of the phosphopantetheine ejection transition from the parent peptide "MDSLNLMDFLVYE" in the +2 charge state.

**Preparation of denatured TcsD supernatants for untargeted LC-MS analysis**

TcsD was expressed and purified via nickel affinity chromatography as described above. The highest purity nickel affinity elution fractions were pooled, and protein was concentrated to approximately 17 mg/mL or 375 μM (determined by absorbance using a Nanodrop and the molar absorbance of TcsD). After concentration, 150 μL of protein was denatured by either boiling for 10 minutes or the addition of 300 μL of LC grade acetonitrile. The denatured protein was pelleted by centrifuging at maximum speed in a benchtop centrifuge for

2 minutes. Supernatants were removed and lyophilized overnight. After lyophilization, supernatants were resuspended in 75 µL of LC grade water for an effective metabolite concentration of 750 µM. Samples were diluted to an effective metabolite concentration of 50 µM in 50:50 (v/v) LC grade water:methanol prior to untargeted LC-MS analysis. Standards of flavin adenine dinucleotide (FAD), Coenzyme A, pantetheine, pantothenate, and butyryl-CoA were prepared in 50:50 (v/v) LC grade methanol:water to a concentration of 20 µM.

**High resolution untargeted LC-MS analysis of denatured TcsD supernatants**

Supernatants of denatured TcsD samples were analyzed using an Agilent 6545 LC-QTOF system as previously described.[20] Analytes were separated using a SeQuant ZIC hydrophilic interaction chromatography (HILIC) column (150 mm length, 4.6 mm internal diameter, 5 µm particle size) coupled to a SeQuant ZIC-pHILIC guard column (20 mm length, 2.1 mm internal diameter, 5 µm particle size). The mobile phase consisted of 10 mM ammonium carbonate and 118.4 mM ammonium hydroxide in acetonitrile-water (60.2:39.8 v/v). Analytes were separated isocratically using the following pump parameters:

| **Time (min)** | **Mobile phase (%)** | **Flow rate (mL/min)** |
|---|---|---|
| 0 | 100 | 0.45 |
| 6 | 100 | 0.45 |
| 6.5 | 100 | 0.605 |
| 12.5 | 100 | 0.605 |

The mass spectrometer source was operated with the following parameters: Gas Temperature = 300°C, Drying Gas = 10 L/min, Nebulizer Pressure = 20 psi, Sheath Gas Temperature = 350°C, Sheath Gas Flow = 12 L/min, and Capillary Voltage = 3500 V, Nozzle Voltage = 2000 V. The TOF mass spectrometer was programmed to operated in scan mode with the following parameters: Fragmentor Voltage = 100 V, Skimmer Voltage = 50 V, Oct 1 RF Vpp = 400 V, Mass Range = 70 m/z – 1100 m/z, Acquisition Rate = 0.86 spectra/s, Acquisition Time = 1162.8 ms/spectrum, and Transients/spectrum = 9532 (negative ion mode) or 9485 (positive ion mode). All samples were analyzed in both negative ion mode and subsequently in positive ion mode. Data analysis was performed in Agilent MassHunter Qualitative Analysis (v B.05.00).

**Substrate modeling into TcsD active site**

Substrates for modeling were drawn using ChemDraw, and ligand restraints were generated using eLBOW in the Phenix software suite. Substrates were manually placed into the TcsD active site using Coot. Real Space Refinements of the substrates were performed using the unknown density in the substrate-binding region of the enzyme active site. The refinements resulted in the generation of both *cis* and *trans* isomers of the substrate with planar fatty acyl tails. After substrate generation, the TcsD structure was aligned with the structures of ACADs which had been co-crystallized with thioester substrates. Alignments were performed using the least squares fit LSQ superpose function of Coot. Proteins were aligned using the 3-residue loop containing the catalytic glutamate and the residues immediately upstream and downstream of the glutamate (Table S6). These alignments were used for analysis of the relative positioning of FAD cofactors and for substrate modeling. After alignment, the 2-pentenoyl substrates were manually translated and rotated into a position within the TcsD active site that corresponds to approximately the equivalent position of substrates in the aligned homolog structures. The feasibility of hydrogen bonding interactions was confirmed using the Measure tool in Coot considering a maximum distance of 3.5Å between heteroatoms. Substrate lengths were also determined using the Measure tool to calculate the distance from the carbonyl carbon of a given molecule to the distal carbon of the fatty acyl tail. The length of the 2-hexenoyl substrate was calculated using a substrate molecule that was generated using Ligand Builder in Coot. The approximate length of the 2-heptenoyl substrate was calculated by adding half the distance between the fourth and sixth (saturated) carbons of the 2-hexenoyl substrate to the total length of the 2-hexenoyl substrate.

**Bioinformatic identification of TcsD homologs**

First, a Hidden Markov Model (HMM) that describes γ,δ-ACADs was generated using the sequences of several TcsD enzymes from various FK506 producers and HliR.[21] The HMM was then used to search for homologs among all proteins in the Uniprot database, after which the genomes of organisms that contain potential γ,δ-ACADs were analyzed using antiSMASH.[22] An additional search was performed by querying the results of the HMM search against GenBank using protein BLAST.[23] Finally, CORASON-BGC was used for the identification of the syntenic genomic contexts of the identified genes.[24] The results from each of these searches were then manually curated based on the presence of key motifs identified through the structural and biochemical characterization of TcsD.

**Sequence alignments of TcsD and homologs**

A sequence alignment of TcsD, HliR, and several α,β-ACADs was generated using Clustal Omega.[25] The sequence alignment was visualized with respect to the TcsD structure using ESPript 3.0.[26] To calculate the positional occupancy of amino acids within the γ,δ-ACADs, a sequence alignment of the identified proteins was generated using MAFFT.[27] Positional occupancies were calculated using the bioinformatics tool UGene.[28]

**Position weight matrix**

The Shannon entropy of the aligned γ,δ-ACADs was calculated at each position within a sequence alignment. All of the identified γ,δ-ACAD sequences were aligned using MAFFT.[27] The MSA was then refined so that all positions maintained at least 10% occupancy using the ProDy python library. The Shannon entropy was then calculated using ProDy.[29,30] The sequence logo was generated using Weblogo (https://weblogo.berkeley.edu/logo.cgi).

### Supplementary Discussion

#### LC-MS/MS Phosphopantetheine ejection assay method development

In order to assay TcsD activity on TcsA-bound substrates, we first developed a targeted proteomics assay for the identification of substrates tethered to the phosphopantetheinyl arm of the acyl carrier proteins (ACP) domain of TcsA. In a typical "phosphopantetheine ejection" experiment,[17] a protein of interest is digested with trypsin and the digested peptides are analyzed via LC-MS/MS (main text Figure 2A). However, after digesting TcsA with trypsin we were unable to detect an ion representing the ACP active site peptide, most likely because the large size (~40 residues) of the tryptic peptide placed it outside of the detectable range of our mass spectrometer (Figure S17). Similarly, the active site peptide could not be detected after digestion with another commonly-used protease, chymotrypsin. In order to obtain an appropriately-sized peptide, we performed a sequential digestion of TcsA with two proteases, Asp-N then trypsin, and identified the resulting "holo" and "acyl" active site peptides using high resolution untargeted LC-MS (Figure S17 and S18). Surprisingly, Asp-N did not consistently cleave at the predicted residue D373 of the conserved "DSL" motif of the acyl carrier protein (ACP) of TcsA but instead at the preceding D369, possibly due to steric effects arising from the presence of the phosphopantetheine arm appended to S374.

#### Unidentified density in active site of TcsD crystal structure

An unknown metabolite consistently co-crystallized in the active site, occupying the space where substrates are predicted to bind (Figure S14). We attempted to identify this unknown density by performing untargeted high resolution LC-MS of denatured protein samples (Figures S15 and S16), but were unable to identify any unique masses in the supernatants of denatured TcsD samples. We also compared the denatured protein supernatant LC-MS chromatograms to several standards, including flavin adenine dinucleotide (FAD), Coenzyme A, pantetheine, pantothenate, and butyryl-CoA, but only the m/z corresponding to FAD could be detected in extracted ion chromatograms of either sample. We additionally used the Ligand Identification function of the Phenix software suite,[31,32,33] attempting to model 180 of the most common ligands from the Protein Data Bank into the density, but none of the ligands fit appropriately.

### Supplementary Data Tables

**Table S1**: Plasmids used in this study

| ICE Entry Number | Plasmid name | Description | Source |
|---|---|---|---|
| JPUB_013738 | pET28a-TcsA | TcsA in pET28a | This study |
| JPUB_013740 | pET28a-TcsAS98A | TcsA S98A in pET28a | This study |
| JPUB_013728 | pET28a-TcsD | TcsD in pET28a | This study |
| JPUB_013730 | pET28a-TcsDL83A | TcsD L83A in pET28a | This study |
| JPUB_013732 | pET28a-TcsDF79A | TcsD F79A in pET28a | This study |
| JPUB_013734 | pET28a-TcsDI363A | TcsD I363A in pET28a | This study |
| JPUB_013736 | pET28a-TcsDF79AL83A | TcsD F79AL83A in pET28a | This study |
| JPUB_013744 | pET28a-TcsDF79AI363A | TcsD F79AI363A in pET28a | This study |
| JPUB_013742 | pBbE7a-PP2216-6xHis | PP2216 in pBbE7a with a C-terminal 6xHis tag | This study |
| JPUB_013842 | pBbE7a-NmexACP-6xHis | Acyl carrier protein from *Nocardia mexicana* NBRC 108244 (*E. coli* codon-optimized) in pBbE7a with a C-terminal 6xHis tag | This study |
| JPUB_013844 | pBbE7a-NmexACAD-6xHis | Acyl-CoA dehydrogenase from *Nocardia mexicana* NBRC 108244 (*E. coli* codon-optimized) in pBbE7a with a C-terminal 6xHis tag | This study |
| N/A | pET-Sfp | Sfp in a pET vector | Reference 34 |
| N/A | pLK54 | MatB mutant | Reference 14 |
| N/A | pBbE7a-RFP | RFP behind a T7 promoter in a BglBrick plasmid | Reference 35 |

**Table S2**: Strains used in this study

| ICE Entry Number | Strain name | Description/Genotype | Source |
|---|---|---|---|
| N/A | *E. coli* DH5α | F⁻ *endA1 glnV44 thi-1 recA1 relA1 gyrA96 deoR nupG purB20* φ80d*lacZ*ΔM15 Δ(*lacZYA-argF*)U169, hsdR17($r_K^- m_K^+$), λ⁻ | QB3 Macrolab (http://qb3.berkeley.edu/macrolab/) |
| N/A | *E. coli* NEB Turbo | F' *proA*$^+$*B*$^+$ *lacI*$^q$ Δ*lacZM15 / fhuA2* Δ*(lac-proAB) glnV galK16 galE15 R(zgb-210::Tn10)*Tet$^S$ *endA1 thi-1* Δ*(hsdS-mcrB)5* | New England Biolabs |
| N/A | *E. coli* BL21 (DE3) | F⁻ *ompT gal dcm lon hsdS*$_B$($r_B^- m_B^-$) λ(DE3 [*lacI lacUV5-T7p07 ind1 sam7 nin5*]) [*malB*$^+$]$_{K-12}$(λ$^S$) | New England Biolabs |
| N/A | *Streptomyces tsukubensis* NRRL 18488 | Wild type strain | US Department of Agriculture, Agricultural Research Service Culture Collection (NRRL), Patent Collection |
| N/A | *Pseudomonas putida* KT2440 (ATCC 47054) | Wild type strain | American Type Culture Collection (ATCC) |
| JPUB_013737 | JBEI-107002 | *E. coli* DH5α containing pet28a-TcsA | This study |
| JPUB_013739 | JBEI-107003 | *E. coli* DH5α containing pet28a-TcsA S98A | This study |
| JPUB_013727 | JBEI-106997 | *E. coli* NEB Turbo containing pet28a-TcsD | This study |
| JPUB_013729 | JBEI-106998 | *E. coli* DH5α containing pet28a-TcsD L83A | This study |
| JPUB_013731 | JBEI-106999 | *E. coli* DH5α containing pet28a-TcsD F79A | This study |
| JPUB_013733 | JBEI-107000 | *E. coli* DH5α containing pet28a-TcsD I363A | This study |
| JPUB_013735 | JBEI-107001 | *E. coli* DH5α containing pet28a-TcsD F79AL83A | This study |
| JPUB_013743 | JBEI-107005 | *E. coli* DH5α containing pet28a-TcsD F79AI363A | This study |
| JPUB_013741 | JBEI-107004 | *E. coli* DH5α containing pBbE7a-PP2216-6xHis | This study |
| JPUB_013841 | JBEI-133594 | *E. coli* DH5α containing pBbE7a-NmexACP-6xHis | This study |
| JPUB_013843 | JBEI-133595 | *E. coli* DH5α containing pBbE7a-NmexACAD-6xHis | This study |

**Table S3**: Primers used in this work

| **Primer name** | **Sequence (5'-->3')** |
|---|---|
| TcsA-NdeI-F | TATACATATGGCGTTCCTCTTCCCCGGC |
| TcsA-XhoI-R | TATACTCGAGCGCCGCCCGGAAACGGAA |
| TcsD-NdeI-F | AAACATATGAGCGAATCCGAACGCCTCGGTA |
| TcsD-XhoI-R | TTTCTCGAGGGTACGTTTCGCGGTGGGGACG |
| TcsA-AT0-F-1 | TCGCCGGGCACGCTCTCGGGGA |
| TcsA-AT0-R-1 | CCACTGGTAACAGGATTAGCAGAGCGAGGTATGT |
| TcsA-AT0-F-2 | CACCGCCTACATACCTCGCTCTGCTAATCC |
| TcsA-AT0-R-2 | GCTCCGAACTCCCCGAGAGCGTGC |
| TcsD-mut-F | CTCTGTAGCACCGCCTACATACCTCGC |
| TcsD-mut-R | ACTGGTAACAGGATTAGCAGAGCGAGG |
| TcsD-F79A-F | GGTGGCGGCCACCCTGTTTCTGC |
| TcsD-F79A-R | AGGACGGGCAGAAACAGGGTGGCCGCC |
| TcsD-L83A-F | ACCCTGTTTGCGCCCGTCCTGAC |
| TcsD-L83A-R | GCTGGTCGTCAGGACGGGCGCAAACAGGGT |
| TcsD-I363-bb-F | GACCCAGAGCGCTGCCGGCACCT |
| TcsD-I363-bb-R | CTCGTAGGACAGGTGCCGGCAGCGC |
| TcsD-I363A-F | CACGCTTCGATCGCCGAGGGCGGCGACGAC |
| TcsD-I363A-R | GTCGTCGCCGCCCTCGGCGATCGAAGCGTG |
| PP2216-E7a-F | CAAAAGATCTTTTAAGAAGGAGATATACATATGCTGGTAAATGACGAGCAACAAC |
| PP2216-E7a-R | CTCGAGCCGCCGCCCGGAAACGGAACCGGTAAGATTGCGCGCAATGACCATGC |
| E7a-link-His-F | GGATCTGGCACTGGTAGT |
| E7a-link-His-R | CATATGTATATCTCCTTCTTAAAAGATCT |
| E7a-link-GG-F | CAGGTCTCAGGATCTGGCACTGGTAGT |
| BioB-GG-R | CAGGTCTCACATATGTATATCTCCTTCTTAAAAGATCTT |

**Table S4**: Synthetic DNA fragments used in this work

| DNA fragment name | Sequence (5’-->3’) |
|---|---|
| *Nocardia mexicana* ACP | caggtctcatATGAACGCGATAACTACAACCGAGATCGCCGAGGGCCTGCGGAGCATCGCCGAT<br>CGTCTGGACCTGGAACTCGAGAATGTCGATATTTCCGCGACGTCGTCGCTGGAAGACGACC<br>TCGAGATGGACTCGCTGAACCTGATGGACTTCCTGGTGTATCTCGAGAAGAGATATCACGT<br>GCAGGTCACCGACGAGCGCCTGCACGAGGTCGACACCATCGGCGATGTCGTGAACCTGCT<br>CAACGACCTGCTGGCATCGTCGCGCGACACCCCGCAGCCGTCCCGAGTCGGAAGTCGTgg<br>atatgagacctg |
| *Nocardia mexicana* ACAD | caggtctcatATGGGCGACATTGTCGAGCAGGCCCGCCATTTTGGACGTACAGTTCTAGCTGCT<br>GCCCCAGCACCTGATGTAGAAACCGTCTACGACGACCGGCACCCACTGTGGGAGCAGTTC<br>CGATCGGCCGACCTCGCCGACTGGTGGGTGCCCGCCGAATACGGCGGACGGGGTGTCG<br>GACTGTGCGAGTCGGTGAACGTCGTCTCCGAACTCTCCTACCACGACGCCGGATTCGCGT<br>TCGCCGCGTTTCTGCCGATCCTCGCGTCACGAATGCTCGAGCTCTACGGTCCCGAGGAGT<br>TGGCGCGCCGCTACTTGGCGGAGATGGCCACCCATGGATCCTTTGCAGCAGCGTTGGGA<br>AGTGAAGCTGAAGCTGGAAGTGAATTAGCTAGAACGCAGACCACGTTCCGCCGCGACGGC<br>GATGTGCTGCACATCAACGGCGATAAGCAGTTCTCGACCAACTTGGCGTTCGCGCGGTTC<br>TGCCTCGTGCTGGCCCGCGACGTCGACAACCCCCGAGATTTCGCCCTGATCCTGGTCCCG<br>TCGGACAGTCCGGGATTTGTTGTCGGCCAACGGTGGCCGATGTCGGGCCTGCACGGTAC<br>TGCCACGTACCCGGCAACTTTCACCGATTGCGTCGTCCCGGCAGCAAATCGGTTGTCGGG<br>CAACGGTATTCGGATCCTGGAGGTCGGGCTCGACGCGAGCCGGATCCTGATGGCCTCGA<br>TCGCGATCGGTCTGACCCGGCGGATTCGGGATCTGAGCATGGACTACGCGGCGAGCAAG<br>CGGCTCGGTGGGCAACCGCTGAACCGCAACGCCGTCTTCGGCGCCCGCATGGGTCAGCT<br>CGAGATGGAACTCGAAACGATGAAGGCCCAGTGCCGGTGCGCCGCGGCCGAATACGACG<br>ACATCTACCAGCGATCCGATCGGGCGGCGGTGTTCTATGCCGACGGCGTGCTGAAGTCG<br>GCGATTGTGGCCAAGATGCATTGTGGTCAGGTCGGCTGGCGGGTCGCGAGTCGGGCGTC<br>GGAGGCGTTCGGCGGTCTGGGCTACACCGCGGGCCACGACATCCAGCGGTGCCTGCGG<br>GACATGCGGCACATCGCGATCGTCGAGGGCGGCGACGACGTGCTGCGTGAACTGCTCTA<br>CGGACGCTACGTCAAGCGCGCATCGCGACGAGGAggatatgagacctg |

**Table S5:** - Summary of crystal parameters, data collection, and refinement statistics. Values in parentheses are for the highest resolution shell.

| | |
|---|---|
| **Crystal parameters** | |
| Space group | P 1 21 1 |
| Unit cell | 69.52 100.06 148.51 90 100.31 90 |
| **Data collection statistics** | |
| Wavelength (Å) | 1.00000 |
| Resolution range (Å) | 68.4 - 1.75 (1.813 - 1.75) |
| Total reflections | 835542 (85389) |
| Unique reflections | 196484 (19281) |
| Multiplicity | 4.3 (4.4) |
| Completeness (%) | 97.65 (96.41) |
| Mean I/sigma(I) | 8.99 (1.39) |
| Wilson B-factor | 19.7 |
| R-merge | 0.1119 (1.063) |
| R-meas | 0.1277 (1.207) |
| R-pim | 0.06086 (0.5664) |
| CC1/2 | 0.997 (0.524) |
| CC* | 0.999 (0.829) |
| **Refinement and model statistics** | |
| Reflections used in refinement | 196470 (19281) |
| Reflections used for R-free | 9864 (1003) |
| R-work | 0.1462 (0.2810) |
| R-free | 0.1787 (0.3138) |
| CC(work) | 0.974 (0.807) |
| CC(free) | 0.959 (0.765) |
| Number of non-hydrogen atoms | 13452 |
| macromolecules | 11781 |
| ligands | 212 |
| solvent | 1459 |
| Protein residues | 1546 |
| RMS(bonds) | 0.012 |
| RMS(angles) | 0.98 |
| Ramachandran favored (%) | 99.28 |
| Ramachandran allowed (%) | 0.72 |
| Ramachandran outliers (%) | 0 |
| Rotamer outliers (%) | 0.08 |
| Clashscore | 3.04 |
| Average B-factor | 27.08 |
| macromolecules | 26.03 |
| ligands | 17.5 |
| solvent | 36.93 |
| Number of TLS groups | 31 |

**Table S6:** ACAD structures used in alignment for substrate modeling

| PDB code | Host organism | Protein description | Amino acids used for LSQ alignment |
|---|---|---|---|
| XXXX | *Streptomyces tsukubensis* NRRL 18488 | TcsD | Ile 363 |
| | | | Glu 364 |
| | | | Gly 365 |
| 1jqi | *Rattus norvegicus* | Rat short chain ACAD | Tyr 367 |
| | | | Glu 368 |
| | | | Gly 369 |
| 2vig | *Homo sapiens* | Human short chain ACAD | Tyr 391 |
| | | | Glu 392 |
| | | | Gly 393 |
| 1udy | *Sus scrofa* | Pig medium chain ACAD | Tyr 375 |
| | | | Glu 376 |
| | | | Gly 377 |
| 1buc | *Megasphaera elsdenii* | Bacterial butyryl-CoA ACAD | Tyr 366 |
| | | | Glu 367 |
| | | | Gly 368 |
| 3mdd | *Sus scrofa* | Pig medium chain ACAD | Tyr 375 |
| | | | Glu 376 |
| | | | Gly 377 |

**Table S7**: Transitions used for the targeted LC-MS/MS-based detection of protein-bound intermediates

| Protein | Peptide | Precursor m/z | Precursor Charge | Product m/z | Product Charge | Collision Energy (eV) |
|---|---|---|---|---|---|---|
| tr\|E9KTG6\|E9KTG6_9ACTN_TcsAS98A | APPPTGGMLAVK | 569.818084 | 2 | 970.539014 | 1 | 18.7 |
| tr\|E9KTG6\|E9KTG6_9ACTN_TcsAS98A | APPPTGGMLAVK | 569.818084 | 2 | 873.48625 | 1 | 18.7 |
| tr\|E9KTG6\|E9KTG6_9ACTN_TcsAS98A | APPPTGGMLAVK | 569.818084 | 2 | 430.302396 | 1 | 18.7 |
| tr\|E9KTG6\|E9KTG6_9ACTN_TcsAS98A | AGELIAAAR | 436.253434 | 2 | 501.314357 | 1 | 14.5 |
| tr\|E9KTG6\|E9KTG6_9ACTN_TcsAS98A | AGELIAAAR | 436.253434 | 2 | 388.230293 | 1 | 14.5 |
| tr\|E9KTG6\|E9KTG6_9ACTN_TcsAS98A | AGELIAAAR | 436.253434 | 2 | 317.193179 | 1 | 14.5 |
| tr\|E9KTG6\|E9KTG6_9ACTN_TcsAS98A | DASALYATTMR | 600.289887 | 2 | 742.355236 | 1 | 19.6 |
| tr\|E9KTG6\|E9KTG6_9ACTN_TcsAS98A | DASALYATTMR | 600.289887 | 2 | 579.291907 | 1 | 19.6 |
| tr\|E9KTG6\|E9KTG6_9ACTN_TcsAS98A | DASALYATTMR | 600.289887 | 2 | 508.254793 | 1 | 19.6 |
| sp\|P39135\|SFP_BACSU | TKPISLEIAK | 550.339706 | 2 | 870.529495 | 1 | 18.1 |
| sp\|P39135\|SFP_BACSU | TKPISLEIAK | 550.339706 | 2 | 573.360639 | 1 | 18.1 |
| sp\|P39135\|SFP_BACSU | TKPISLEIAK | 550.339706 | 2 | 460.276575 | 1 | 18.1 |
| sp\|P39135\|SFP_BACSU | TKPISLEIAK | 550.339706 | 2 | 331.233982 | 1 | 18.1 |
| Holo-ACP | DLGVDSLAMTELQAHALQR[+340.085794] | 1204.571117 | 2 | 261.126739 | 1 | 30 |
| Holo-ACP | DLGVDSLAMTELQAHALQR[+340.085794] | 803.38317 | 3 | 261.126739 | 1 | 30 |
| Holo-ACP | DSLAMTELQAHALQR[+340.085794] | 1012.470675 | 2 | 261.126739 | 1 | 30 |
| Holo-ACP | DSLAMTELQAHALQR[+340.085794] | 675.316209 | 3 | 261.126739 | 1 | 30 |
| Apo-ACP | DLGVDSLAMTELQAHALQR | 1034.52822 | 2 | 1297.668131 | 1 | 33.1 |
| Apo-ACP | DLGVDSLAMTELQAHALQR | 1034.52822 | 2 | 1166.627646 | 1 | 33.1 |
| Apo-ACP | DLGVDSLAMTELQAHALQR | 1034.52822 | 2 | 1065.579967 | 1 | 33.1 |
| Apo-ACP | DLGVDSLAMTELQAHALQR | 1034.52822 | 2 | 936.537374 | 1 | 33.1 |
| Apo-ACP | DLGVDSLAMTELQAHALQR | 1034.52822 | 2 | 823.45331 | 1 | 33.1 |
| Apo-ACP | DLGVDSLAMTELQAHALQR | 690.021239 | 3 | 936.537374 | 1 | 20 |
| Apo-ACP | DLGVDSLAMTELQAHALQR | 690.021239 | 3 | 823.45331 | 1 | 20 |
| Apo-ACP | DLGVDSLAMTELQAHALQR | 690.021239 | 3 | 695.394733 | 1 | 20 |
| Apo-ACP | DLGVDSLAMTELQAHALQR | 690.021239 | 3 | 624.357619 | 1 | 20 |
| Apo-ACP | DLGVDSLAMTELQAHALQR | 690.021239 | 3 | 487.298707 | 1 | 20 |
| Apo-ACP | DSLAMTELQAHALQR | 842.427778 | 2 | 1166.627646 | 1 | 27.1 |
| Apo-ACP | DSLAMTELQAHALQR | 842.427778 | 2 | 1065.579967 | 1 | 27.1 |
| Apo-ACP | DSLAMTELQAHALQR | 842.427778 | 2 | 936.537374 | 1 | 27.1 |
| Apo-ACP | DSLAMTELQAHALQR | 842.427778 | 2 | 823.45331 | 1 | 27.1 |

| | | | | | | |
|---|---|---|---|---|---|---|
| Apo-ACP | DSLAMTELQAHALQR | 842.427778 | 2 | 695.394733 | 1 | 27.1 |
| Apo-ACP | DSLAMTELQAHALQR | 561.954277 | 3 | 823.45331 | 1 | 15.4 |
| Apo-ACP | DSLAMTELQAHALQR | 561.954277 | 3 | 695.394733 | 1 | 15.4 |
| Apo-ACP | DSLAMTELQAHALQR | 561.954277 | 3 | 624.357619 | 1 | 15.4 |
| Apo-ACP | DSLAMTELQAHALQR | 561.954277 | 3 | 487.298707 | 1 | 15.4 |
| Apo-ACP | DSLAMTELQAHALQR | 561.954277 | 3 | 416.261593 | 1 | 15.4 |
| 2-pentenoyl-ACP | DLGVDSLAMTELQAHALQR[+422.127659] | 1245.59205 | 2 | 343.168604 | 1 | 30 |
| 2-pentenoyl-ACP | DLGVDSLAMTELQAHALQR[+422.127659] | 830.730459 | 3 | 343.168604 | 1 | 30 |
| 2-pentenoyl-ACP | DSLAMTELQAHALQR[+422.127659] | 1053.491607 | 2 | 343.168604 | 1 | 30 |
| 2-pentenoyl-ACP | DSLAMTELQAHALQR[+422.127659] | 702.663497 | 3 | 343.168604 | 1 | 30 |
| 2_4-pentadienoyl-ACP | DLGVDSLAMTELQAHALQR[+420.112009] | 1244.584225 | 2 | 341.152954 | 1 | 30 |
| 2_4-pentadienoyl-ACP | DLGVDSLAMTELQAHALQR[+420.112009] | 830.058575 | 3 | 341.152954 | 1 | 30 |
| 2_4-pentadienoyl-ACP | DSLAMTELQAHALQR[+420.112009] | 1052.483782 | 2 | 341.152954 | 1 | 30 |
| 2_4-pentadienoyl-ACP | DSLAMTELQAHALQR[+420.112009] | 701.991614 | 3 | 341.152954 | 1 | 30 |
| Hydroxypentanoyl-ACP | DLGVDSLAMTELQAHALQR[+440.138224] | 1254.597332 | 2 | 361.179169 | 1 | 30 |
| Hydroxypentanoyl-ACP | DLGVDSLAMTELQAHALQR[+440.138224] | 836.73398 | 3 | 361.179169 | 1 | 30 |
| Hydroxypentanoyl-ACP | DSLAMTELQAHALQR[+440.138224] | 1062.49689 | 2 | 361.179169 | 1 | 30 |
| Hydroxypentanoyl-ACP | DSLAMTELQAHALQR[+440.138224] | 708.667019 | 3 | 361.179169 | 1 | 30 |
| Dihydroxypentanoyl-ACP | DLGVDSLAMTELQAHALQR[+456.133138] | 1262.594789 | 2 | 377.174083 | 1 | 30 |
| Dihydroxypentanoyl-ACP | DLGVDSLAMTELQAHALQR[+456.133138] | 842.065618 | 3 | 377.174083 | 1 | 30 |
| Dihydroxypentanoyl-ACP | DSLAMTELQAHALQR[+456.133138] | 1070.494347 | 2 | 377.174083 | 1 | 30 |
| Dihydroxypentanoyl-ACP | DSLAMTELQAHALQR[+456.133138] | 713.998657 | 3 | 377.174083 | 1 | 30 |
| 3-hydroxy-45-pentenoyl-ACP | DLGVDSLAMTELQAHALQR[+438.122574] | 1253.589507 | 2 | 359.163519 | 1 | 30 |
| 3-hydroxy-45-pentenoyl-ACP | DLGVDSLAMTELQAHALQR[+438.122574] | 836.062097 | 3 | 359.163519 | 1 | 30 |
| 3-hydroxy-45-pentenoyl-ACP | DSLAMTELQAHALQR[+438.122574] | 1061.489065 | 2 | 359.163519 | 1 | 30 |
| 3-hydroxy-45-pentenoyl-ACP | DSLAMTELQAHALQR[+438.122574] | 707.995135 | 3 | 359.163519 | 1 | 30 |
| 2-hexenoyl-ACP | DLGVDSLAMTELQAHALQR[+436.143309] | 1252.599875 | 2 | 357.184254 | 1 | 30 |
| 2-hexenoyl-ACP | DLGVDSLAMTELQAHALQR[+436.143309] | 835.402342 | 3 | 357.184254 | 1 | 30 |
| 2-hexenoyl-ACP | DSLAMTELQAHALQR[+436.143309] | 1060.499432 | 2 | 357.184254 | 1 | 30 |
| 2-hexenoyl-ACP | DSLAMTELQAHALQR[+436.143309] | 707.33538 | 3 | 357.184254 | 1 | 30 |
| Hydroxyhexanoyl-ACP | DLGVDSLAMTELQAHALQR[+454.153874] | 1261.605157 | 2 | 375.194819 | 1 | 30 |
| Hydroxyhexanoyl-ACP | DLGVDSLAMTELQAHALQR[+454.153874] | 841.405864 | 3 | 375.194819 | 1 | 30 |
| Hydroxyhexanoyl-ACP | DSLAMTELQAHALQR[+454.153874] | 1069.504715 | 2 | 375.194819 | 1 | 30 |

| | | | | | | |
|---|---|---|---|---|---|---|
| Hydroxyhexanoyl-ACP | DSLAMTELQAHALQR[+454.153874] | 713.338902 | 3 | 375.194819 | 1 | 30 |
| 3-hydroxy-45-hexenoyl-ACP | DLGVDSLAMTELQAHALQR[+452.138224] | 1260.597332 | 2 | 373.179169 | 1 | 30 |
| 3-hydroxy-45-hexenoyl-ACP | DLGVDSLAMTELQAHALQR[+452.138224] | 840.73398 | 3 | 373.179169 | 1 | 30 |
| 3-hydroxy-45-hexenoyl-ACP | DSLAMTELQAHALQR[+452.138224] | 1068.49689 | 2 | 373.179169 | 1 | 30 |
| 3-hydroxy-45-hexenoyl-ACP | DSLAMTELQAHALQR[+452.138224] | 712.667019 | 3 | 373.179169 | 1 | 30 |
| Dihydroxyhexanoyl-ACP | DLGVDSLAMTELQAHALQR[+470.148788] | 1269.602614 | 2 | 391.189733 | 1 | 30 |
| Dihydroxyhexanoyl-ACP | DLGVDSLAMTELQAHALQR[+470.148788] | 846.737502 | 3 | 391.189733 | 1 | 30 |
| Dihydroxyhexanoyl-ACP | DSLAMTELQAHALQR[+470.148788] | 1077.502172 | 2 | 391.189733 | 1 | 30 |
| Dihydroxyhexanoyl-ACP | DSLAMTELQAHALQR[+470.148788] | 718.67054 | 3 | 391.189733 | 1 | 30 |
| 2_4-hexadienoyl-ACP | DLGVDSLAMTELQAHALQR[+434.127659] | 1251.59205 | 2 | 355.168604 | 1 | 30 |
| 2_4-hexadienoyl-ACP | DLGVDSLAMTELQAHALQR[+434.127659] | 834.730459 | 3 | 355.168604 | 1 | 30 |
| 2_4-hexadienoyl-ACP | DSLAMTELQAHALQR[+434.127659] | 1059.491607 | 2 | 355.168604 | 1 | 30 |
| 2_4-hexadienoyl-ACP | DSLAMTELQAHALQR[+434.127659] | 706.663497 | 3 | 355.168604 | 1 | 30 |
| tr\|E9KTG9\|E9KTG9_9ACTN_TcsD | DAPLALYER | 524.277106 | 2 | 861.482879 | 1 | 17.3 |
| tr\|E9KTG9\|E9KTG9_9ACTN_TcsD | DAPLALYER | 524.277106 | 2 | 580.308937 | 1 | 17.3 |
| tr\|E9KTG9\|E9KTG9_9ACTN_TcsD | DAPLALYER | 524.277106 | 2 | 467.224873 | 1 | 17.3 |
| tr\|E9KTG9\|E9KTG9_9ACTN_TcsD | DAPLALYER | 524.277106 | 2 | 304.161545 | 1 | 17.3 |
| tr\|E9KTG9\|E9KTG9_9ACTN_TcsD | YTAVTVPR | 453.755809 | 2 | 472.287808 | 1 | 15.1 |
| tr\|E9KTG9\|E9KTG9_9ACTN_TcsD | YTAVTVPR | 453.755809 | 2 | 371.24013 | 1 | 15.1 |
| tr\|E9KTG9\|E9KTG9_9ACTN_TcsD | YTAVTVPR | 453.755809 | 2 | 272.171716 | 1 | 15.1 |
| 2-heptenoyl-ACP | DLGVDSLAMTELQAHALQR[+450.158959] | 1259.6077 | 2 | 371.199904 | 1 | 30 |
| 2-heptenoyl-ACP | DLGVDSLAMTELQAHALQR[+450.158959] | 840.074225 | 3 | 371.199904 | 1 | 30 |
| 2-heptenoyl-ACP | DSLAMTELQAHALQR[+450.158959] | 1067.507257 | 2 | 371.199904 | 1 | 30 |
| 2-heptenoyl-ACP | DSLAMTELQAHALQR[+450.158959] | 712.007264 | 3 | 371.199904 | 1 | 30 |
| 2_4-heptadienoyl-ACP | DLGVDSLAMTELQAHALQR[+448.143309] | 1258.599875 | 2 | 369.184254 | 1 | 30 |
| 2_4-heptadienoyl-ACP | DLGVDSLAMTELQAHALQR[+448.143309] | 839.402342 | 3 | 369.184254 | 1 | 30 |
| 2_4-heptadienoyl-ACP | DSLAMTELQAHALQR[+448.143309] | 1066.499432 | 2 | 369.184254 | 1 | 30 |
| 2_4-heptadienoyl-ACP | DSLAMTELQAHALQR[+448.143309] | 711.33538 | 3 | 369.184254 | 1 | 30 |
| Hydroxyheptanoyl-ACP | DLGVDSLAMTELQAHALQR[+468.169524] | 1268.612982 | 2 | 389.210469 | 1 | 30 |
| Hydroxyheptanoyl-ACP | DLGVDSLAMTELQAHALQR[+468.169524] | 846.077747 | 3 | 389.210469 | 1 | 30 |
| Hydroxyheptanoyl-ACP | DSLAMTELQAHALQR[+468.169524] | 1076.51254 | 2 | 389.210469 | 1 | 30 |
| Hydroxyheptanoyl-ACP | DSLAMTELQAHALQR[+468.169524] | 718.010785 | 3 | 389.210469 | 1 | 30 |
| Dihydroxyheptanoyl-ACP | DLGVDSLAMTELQAHALQR[+484.164439] | 1276.61044 | 2 | 405.205384 | 1 | 30 |

| | | | | | | |
|---|---|---|---|---|---|---|
| Dihydroxyheptanoyl-ACP | DLGVDSLAMTELQAHALQR[+484.164439] | 851.409385 | 3 | 405.205384 | 1 | 30 |
| Dihydroxyheptanoyl-ACP | DSLAMTELQAHALQR[+484.164439] | 1084.509997 | 2 | 405.205384 | 1 | 30 |
| Dihydroxyheptanoyl-ACP | DSLAMTELQAHALQR[+484.164439] | 723.342424 | 3 | 405.205384 | 1 | 30 |
| 3-hydroxy-45-heptenoyl-ACP | DLGVDSLAMTELQAHALQR[+466.153874] | 1267.605157 | 2 | 387.194819 | 1 | 30 |
| 3-hydroxy-45-heptenoyl-ACP | DLGVDSLAMTELQAHALQR[+466.153874] | 845.405864 | 3 | 387.194819 | 1 | 30 |
| 3-hydroxy-45-heptenoyl-ACP | DSLAMTELQAHALQR[+466.153874] | 1075.504715 | 2 | 387.194819 | 1 | 30 |
| 3-hydroxy-45-heptenoyl-ACP | DSLAMTELQAHALQR[+466.153874] | 717.338902 | 3 | 387.194819 | 1 | 30 |
| Pentanoyl-ACP | DLGVDSLAMTELQAHALQR[+424.1] | 1246.599875 | 2 | 345.184254 | 1 | 30 |
| Pentanoyl-ACP | DLGVDSLAMTELQAHALQR[+424.1] | 831.402342 | 3 | 345.184254 | 1 | 30 |
| Pentanoyl-ACP | DSLAMTELQAHALQR[+424.1] | 1054.499432 | 2 | 345.184254 | 1 | 30 |
| Pentanoyl-ACP | DSLAMTELQAHALQR[+424.1] | 703.33538 | 3 | 345.184254 | 1 | 30 |
| Butyryl-ACP | DLGVDSLAMTELQAHALQR[+410.1] | 1239.59205 | 2 | 331.168604 | 1 | 30 |
| Butyryl-ACP | DLGVDSLAMTELQAHALQR[+410.1] | 826.730459 | 3 | 331.168604 | 1 | 30 |
| Butyryl-ACP | DSLAMTELQAHALQR[+410.1] | 1047.491607 | 2 | 331.168604 | 1 | 30 |
| Butyryl-ACP | DSLAMTELQAHALQR[+410.1] | 698.663497 | 3 | 331.168604 | 1 | 30 |
| Crotonyl-ACP | DLGVDSLAMTELQAHALQR[+408.1] | 1238.584225 | 2 | 329.152954 | 1 | 30 |
| Crotonyl-ACP | DLGVDSLAMTELQAHALQR[+408.1] | 826.058575 | 3 | 329.152954 | 1 | 30 |
| Crotonyl-ACP | DSLAMTELQAHALQR[+408.1] | 1046.483782 | 2 | 329.152954 | 1 | 30 |
| Crotonyl-ACP | DSLAMTELQAHALQR[+408.1] | 697.991614 | 3 | 329.152954 | 1 | 30 |
| Hydroxybutyryl-ACP | DLGVDSLAMTELQAHALQR[+426.1] | 1247.589507 | 2 | 347.163519 | 1 | 30 |
| Hydroxybutyryl-ACP | DLGVDSLAMTELQAHALQR[+426.1] | 832.062097 | 3 | 347.163519 | 1 | 30 |
| Hydroxybutyryl-ACP | DSLAMTELQAHALQR[+426.1] | 1055.489065 | 2 | 347.163519 | 1 | 30 |
| Hydroxybutyryl-ACP | DSLAMTELQAHALQR[+426.1] | 703.995135 | 3 | 347.163519 | 1 | 30 |
| Propylmalonyl-ACP | DLGVDSLAMTELQAHALQR[+468.1] | 1268.594789 | 2 | 389.174083 | 1 | 30 |
| Propylmalonyl-ACP | DLGVDSLAMTELQAHALQR[+468.1] | 846.065618 | 3 | 389.174083 | 1 | 30 |
| Propylmalonyl-ACP | DSLAMTELQAHALQR[+468.1] | 1076.494347 | 2 | 389.174083 | 1 | 30 |
| Propylmalonyl-ACP | DSLAMTELQAHALQR[+468.1] | 717.998657 | 3 | 389.174083 | 1 | 30 |
| Allymalonyl-ACP | DLGVDSLAMTELQAHALQR[+466.1] | 1267.586964 | 2 | 387.158433 | 1 | 30 |
| Allymalonyl-ACP | DLGVDSLAMTELQAHALQR[+466.1] | 845.393735 | 3 | 387.158433 | 1 | 30 |
| Allymalonyl-ACP | DSLAMTELQAHALQR[+466.1] | 1075.486522 | 2 | 387.158433 | 1 | 30 |
| Allymalonyl-ACP | DSLAMTELQAHALQR[+466.1] | 717.326773 | 3 | 387.158433 | 1 | 30 |

Note: peptides with added masses in brackets (e.g. "DSLAMTELQAHALQR[+466.1]") depict modified peptides with phosphopantetheine arms/substrates.

**Table S8**: Transitions used for the targeted LC-MS/MS-based detection of protein-bound intermediates

| **Protein** | **Peptide** | **Precursor Ion** | **Precursor charge** | **Product Ion** | **Product charge** | **Collision Energy (eV)** |
|---|---|---|---|---|---|---|
| Holo-Nmex-ACP | MDSLNLMDFLVYLE[+340.085794].light | 1021.94986 | 2 | 261.12674 | 1 | 30 |
| Holo-Nmex-ACP | MDSLNLMDFLVYLE[+340.085794].light | 681.635668 | 3 | 261.12674 | 1 | 30 |
| Nmex-ACP | GLRSIADRLDLE.light | 679.37534 | 2 | 760.38356 | 1 | 22.1 |
| Nmex-ACP | GLRSIADRLDLE.light | 679.37534 | 2 | 645.35662 | 1 | 22.1 |
| Nmex-ACP | GLRSIADRLDLE.light | 679.37534 | 2 | 489.25551 | 1 | 22.1 |
| Nmex-ACP | GLRSIADRLDLE.light | 453.252652 | 3 | 489.25551 | 1 | 11.5 |
| Nmex-ACP | GLRSIADRLDLE.light | 453.252652 | 3 | 376.17144 | 1 | 11.5 |
| Nmex-ACP | GLRSIADRLDLE.light | 453.252652 | 3 | 261.1445 | 1 | 11.5 |
| Nmex-ACP | NVDISATSSLEDDLE.light | 804.367772 | 2 | 820.35707 | 1 | 25.9 |
| Nmex-ACP | NVDISATSSLEDDLE.light | 804.367772 | 2 | 733.32504 | 1 | 25.9 |
| Nmex-ACP | NVDISATSSLEDDLE.light | 804.367772 | 2 | 620.24098 | 1 | 25.9 |
| Pentanoyl-Nmex-ACP | MDSLNLMDFLVYLE[+424.143309].light | 1063.97862 | 2 | 345.18425 | 1 | 30 |
| Pentanoyl-Nmex-ACP | MDSLNLMDFLVYLE[+424.143309].light | 709.65484 | 3 | 345.18425 | 1 | 30 |
| Pentenoyl-Nmex-ACP | MDSLNLMDFLVYLE[+422.127659].light | 1062.9708 | 2 | 343.1686 | 1 | 30 |
| Pentenoyl-Nmex-ACP | MDSLNLMDFLVYLE[+422.127659].light | 708.982956 | 3 | 343.1686 | 1 | 30 |
| Pentadienoyl-Nmex-ACP | MDSLNLMDFLVYLE[+420.112009].light | 1061.96297 | 2 | 341.15295 | 1 | 30 |
| Pentadienoyl-Nmex-ACP | MDSLNLMDFLVYLE[+420.112009].light | 708.311073 | 3 | 341.15295 | 1 | 30 |
| Hydroxypentanoyl-Nmex-ACP | MDSLNLMDFLVYLE[+440.138224].light | 1071.97608 | 2 | 361.17917 | 1 | 30 |
| Hydroxypentanoyl-Nmex-ACP | MDSLNLMDFLVYLE[+440.138224].light | 714.986478 | 3 | 361.17917 | 1 | 30 |
| Butyryl-Nmex-ACP | MDSLNLMDFLVYLE[+410.127659].light | 1056.9708 | 2 | 331.1686 | 1 | 30 |
| Butyryl-Nmex-ACP | MDSLNLMDFLVYLE[+410.127659].light | 704.982956 | 3 | 331.1686 | 1 | 30 |
| Crotonyl-Nmex-ACP | MDSLNLMDFLVYLE[+408.112009].light | 1055.96297 | 2 | 329.15295 | 1 | 30 |
| Crotonyl-Nmex-ACP | MDSLNLMDFLVYLE[+408.112009].light | 704.311073 | 3 | 329.15295 | 1 | 30 |
| Hydroxybutyryl-Nmex-ACP | MDSLNLMDFLVYLE[+426.122574].light | 1064.96825 | 2 | 347.16352 | 1 | 30 |
| Hydroxybutyryl-Nmex-ACP | MDSLNLMDFLVYLE[+426.122574].light | 710.314595 | 3 | 347.16352 | 1 | 30 |
| Hexanoyl-Nmex-ACP | MDSLNLMDFLVYLE[+438.158959].light | 1070.98645 | 2 | 359.1999 | 1 | 30 |
| Hexanoyl-Nmex-ACP | MDSLNLMDFLVYLE[+438.158959].light | 714.326723 | 3 | 359.1999 | 1 | 30 |
| Hexenoyl-Nmex-ACP | MDSLNLMDFLVYLE[+436.143309].light | 1069.97862 | 2 | 357.18425 | 1 | 30 |
| Hexenoyl-Nmex-ACP | MDSLNLMDFLVYLE[+436.143309].light | 713.65484 | 3 | 357.18425 | 1 | 30 |
| Hexadienoyl-Nmex-ACP | MDSLNLMDFLVYLE[+434.127659].light | 1068.9708 | 2 | 355.1686 | 1 | 30 |

| | | | | | | |
|---|---|---|---|---|---|---|
| Hexadienoyl-Nmex-ACP | MDSLNLMDFLVYLE[+434.127659].light | 712.982956 | 3 | 355.1686 | 1 | 30 |
| Hydroxyhexanoyl-Nmex-ACP | MDSLNLMDFLVYLE[+454.153874].light | 1078.9839 | 2 | 375.19482 | 1 | 30 |
| Hydroxyhexanoyl-Nmex-ACP | MDSLNLMDFLVYLE[+454.153874].light | 719.658361 | 3 | 375.19482 | 1 | 30 |
| Heptanoyl-ACP | MDSLNLMDFLVYLE[+452.174609].light | 1077.99427 | 2 | 373.21555 | 1 | 30 |
| Heptanoyl-ACP | MDSLNLMDFLVYLE[+452.174609].light | 718.998606 | 3 | 373.21555 | 1 | 30 |
| Hydroxyheptanoyl-Nmex-ACP | MDSLNLMDFLVYLE[+468.169524].light | 1085.99173 | 2 | 389.21047 | 1 | 30 |
| Hydroxyheptanoyl-Nmex-ACP | MDSLNLMDFLVYLE[+468.169524].light | 724.330245 | 3 | 389.21047 | 1 | 30 |
| Heptenoyl-Nmex-ACP | MDSLNLMDFLVYLE[+450.158959].light | 1076.98645 | 2 | 371.1999 | 1 | 30 |
| Heptenoyl-Nmex-ACP | MDSLNLMDFLVYLE[+450.158959].light | 718.326723 | 3 | 371.1999 | 1 | 30 |
| Heptadienoyl-Nmex-ACP | MDSLNLMDFLVYLE[+448.143309].light | 1075.97862 | 2 | 369.18425 | 1 | 30 |
| Heptadienoyl-Nmex-ACP | MDSLNLMDFLVYLE[+448.143309].light | 717.65484 | 3 | 369.18425 | 1 | 30 |
| sp\|P39135\|SFP_BACSU | YSDLLAKDKDE.light | 648.819532 | 2 | 705.34136 | 1 | 21.1 |
| sp\|P39135\|SFP_BACSU | YSDLLAKDKDE.light | 648.819532 | 2 | 634.30425 | 1 | 21.1 |
| sp\|P39135\|SFP_BACSU | YSDLLAKDKDE.light | 648.819532 | 2 | 506.20928 | 1 | 21.1 |

**Table S9: Protein sequences used to build γ,δ-ACAD HMM used in genome mining**

| GenBank Accession Number | Host organism | Sequence |
|---|---|---|
| ADU56309.1 | *Streptomyces sp.* KCTC 11604BP | MSESERLGIVRDFVAREILGREGILDSLADAPLALYERFAETGLMNWWV PKEHGGLGLGLEESVRIVSELAYGDAGVAFTLFLPVLTTSMIGWYGSEEL KERFLGPLVARRGFCATLGSEHEAGSELARISTTVRRDGDTLVLDGTKA FSTSTDFARFLVVIARSADDPARYTAVTVPRDAPGLRVDKRWDVIGMRA SATYQVSFSDCRVPGDNALNGNGLRLLEIGLNASRILIAASALGVARRIR DVCMEYGKTKSLKGAPLVKDGVFAGRLGQFEMQIDVMANQCLAAARAY DATAARPDAARVLLRQGAQKSALTAKMFCGQTAWQIASTASEMFGGIG YTHDMVIGKLLRDVRHASIIEGGDDVLRDLVYQRFVVPTAKRT |
| WP_055551085.1 | *Streptomyces kanamyceticus* | MSEPEHLDTVRKFVAQEVLGRETHLDSLADAPLALYERFAETGLMNWW VPEEHGGLGLGLEDSVRIVSELAYGDAGVAFTLFLPILTTSMVSWYGSA ELKEKLLDPLVAHRGFCATLGSEHEAGSELAKISTVVRRDGEGLVLDGT KAFSTSTDFAQFLVVIARSAENPTRYLAVAVERDAPGLRIDKRWDVIGLR ASATYQVSFSDCHVPAGNALDGHGLRLLEIGLNASRILIAATALGVARRIR DLCMEYAKTKSLKGAPLVNDAVFAGRLGQFEMQIEVMANQCLAAARTY DATAARPDAARTLLRQGAQKSALTAKMFCGQTAWQIASTASEMFGGIG YTHDVPIGKLLRDVRHASIIEGGDDVLRDLVFHRFVVPTAKRT |
| WP_006350839.1 | *Streptomyces tsukubensis* | MSESERLGIVRDFVAREILGREGILDSLADAPLALYERFAETGLMNWWV PKEHGGLGLGLEESVRIVSELAYGDAGVAFTLFLPVLTTSMIGWYGSEEL KERFLGPLVARRGFCATLGSEHEAGSELARISTTVRRDGDTLVLDGTKA FSTSTDFARFLVVIARSADDPARYTAVTVPRDAPGLRVDKRWDVIGMRA SATYQVSFSDCRVPGDNALNGNGLRLLEIGLNASRILIAASALGVARRIR DVCMEYGKTKSLKGAPLVKDGVFAGRLGQFEMQIDVMANQCLAAARAY DATAARPDAARVLLRQGAQKSALTAKMFCGQTAWQIASTASEMFGGIG YTHDMVIGKLLRDVRHASIIEGGDDVLRDLVYQRFVVPTAKRT |
| AMM72019.1 | *Haliangium ochraceum* DSM 14365 | MSADTTKKNPLIEPIRGFVREHVLGREQQLDAGGELPLDIYEAFRKAGLA NWWLPESYGGHGLSLEQSVDIVAELAYGDAGLAFAFFLPILSTSVIEQFG SEEQKQRYLPALAKSGGSCATMASEEKAGSELIRTEALARGSAEEGFKL SGDKYFSTNADTAELLIVYARIAGPTPAYGAFLVPRSADGIRIVRRWEMN GLRASGTYELELRDCPAESQLAGNGLRILEVGLNSSRTLMAACAVGIAR RVRDVCLGYARNKEIKNTKLFNNHVFGAKLGQMEAELDGMMAVCRTAA REMDEIASREDAAKVFFREGTLKSVIVAKMLCGQLGWKIASVGSESLGG LGYTHDSIIGKLLRDVRYVSLVEAGDDVLRDLVFSRHVLPRFMSEIE |

**Table S10: Uncharacterized homologs of TcsD that contain characteristic γ,δ-ACAD motifs**

| Species | Protein GenBank Accession | DNA GenBank Accession | Gene cluster type (Antismash annotation) | Closest homologous cluster (Antismash) |
|---|---|---|---|---|
| Streptomyces kanamyceticus | ADU56239.1 | HM116536.1 | N/A | Allylmalonyl-ACP/FK506 (T1PKS) |
| Streptomyces tacrolimicus | ADU56353.1 | HM116538.1 | N/A | Allylmalonyl-ACP/FK506 (T1PKS) |
| Streptomyces tsukubensis VKM Ac-2618D | WP_006350839.1 | NZ_SGFG01000008.1 | N/A | Allylmalonyl-ACP/FK506 (T1PKS) |
| Streptomyces sp. KCTC 11604BP | ADU56309.1 | HM116537.1 | N/A | Allylmalonyl-ACP/FK506 (T1PKS) |
| Streptomyces tsukubensis NRRL 18488 | WP_006350839.1 | NZ_AJSZ01000908.1 | N/A | Allylmalonyl-ACP/FK506 (T1PKS) |
| Streptomyces tsukubensis F601 | WP_077974278.1 | NZ_MVFC01000056.1 | N/A | Allylmalonyl-ACP/FK506 (T1PKS) |
| Streptomyces sp. MJM7001 | WP_006350839.1 | HQ696504.1 | N/A | Allylmalonyl-ACP/FK506 (T1PKS) |
| Haliangium ochraceum DSM 14365 | AMM72019.1 | KU523555.1 | N/A | Haliangicin (T1PKS) |
| Nocardia brasiliensis ATCC 700358 | WP_014984668.1 | NC_018681.1 | T1PKS | Akaeolide (polyketide) (16%) |
| Nocardia brasiliensis HUJEG-1 isolate P-200 | WP_014984668.1 | NZ_LRRM01000006.1 | T1PKS | Akaeolide (polyketide) (16%) |
| Saccharomonospora saliphila YIM 90502 | WP_019815635.1 | NZ_KB912660.1 | PKS-like,T1PKS | Arsenopolyketides (45%) |
| Streptomyces sp. ADI96-15 | WP_023416081.1 | NZ_ML123109.1 | PKS-like,T1PKS | Arsenopolyketides (54%) |
| Streptomyces sp. PVA 94-07 | WP_023416081.1/ESQ07377.1 | NZ_CM002273.1 | PKS-like,T1PKS | Arsenopolyketides (58%) |
| Streptomyces sp. Root63 | WP_023416081.1 | NZ_LMGX01000018.1 | PKS-like,T1PKS | Arsenopolyketides (58%) |
| Streptomyces sp. Root1295 | WP_023416081.1 | NZ_LMEL01000021.1 | PKS-like,T1PKS | Arsenopolyketides (58%) |

| | | | | |
|---|---|---|---|---|
| Streptomyces sp. GBA 94-10 | WP_023416081.1 | NZ_CM002271.1 | PKS-like,T1PKS | Arsenopolyketides (58%) |
| Streptomyces sp. CB00072 | WP_073868671.1 | NZ_LIPB01000003.1 | PKS-like,T1PKS | Arsenopolyketides (58%) |
| Millisia brevis | WP_066907456.1 | NZ_BCRN01000007.1 | NRPS,T1PKS | Aurantimycin (18%) |
| Pseudonocardia endophytica | WP_132431031.1 | NZ_SMFZ01000002.1 | NRPS,PKS-like,T1PKS | Butyrolactol A (46%) |
| Herbidospora sp. NEAU-GS14 | WP_137246715.1 | NZ_SZQA01000007.1 | T1PKS | Butyrolactol A (33%) |
| Herbidospora yilanensis strain NBRC 106371 | WP_062352189.1 | NZ_BBXE01000030.1 | NRPS,T1PKS | Butyrolactol A (40%) |
| Streptomyces sp. AmelKG-E11A | WP_099283133.1 | NZ_AQRJ01000070.1 | T1PKS/NRPS/lassopeptide | Butyrolactol A (40%) |
| Streptomyces uncialis | WP_073788609.1 | NZ_LFBV01000001.1 | T1PKS/NRPS/lassopeptide | Butyrolactol A (40%) |
| Herbidospora cretacea strain NRRL B-16917 | WP_030450088.1 | NZ_JODQ01000001.1 | NRPS,T1PKS,arylpolyene | Butyrolactol A (46%) |
| Herbidospora daliensis strain NBRC 106372 | WP_062432695.1 | NZ_BBXF01000003.1 | NRPS,T1PKS,arylpolyene | Butyrolactol A (46%) |
| Herbidospora sakaeratensis strain NBRC 102641 | WP_062330499.1 | NZ_BBXC01000007.1 | NRPS,T1PKS,ladderane | Butyrolactol A (46%) |
| Herbidospora mongoliensis strain NBRC 105882 | WP_066370148.1 | NZ_BBXD01000017.1 | NRPS,T1PKS,ladderane | Butyrolactol A (46%) |
| Alloactinosynnema album | WP_091377637.1 | | T1PKS/NRPS | Butyrolactol A (46%) |
| Nonomuraea sp. PA1-10 | WP_139634331 | NZ_VDLX01000013.1 | NRPS,T1PKS | Butyrolactol A (53%) |
| Nonomuraea coxensis DSM 45129 | WP_020541001.1 | NZ_KB903944.1 | T1PKS | Butyrolactol A (66%) |
| Labedaea rhizosphaerae DSM 45361 | WP_133849167.1 | NZ_SNXZ01000002.1 | NRPS,T1PKS,betalactone,transAT-PKS-like | Butyrolactol A (40%) |
| Nocardia vulneris W9851 | WP_052281359.1 | NZ_JNFP00000000.1 | T1PKS | Chalcomycin (9%) |
| Nocardia vulneris NBRC 108936 | WP_052281359.1 | NZ_BDCI01000001.1 | T1PKS | Chalcomycin (9%) |
| Amycolatopsis coloradensis | WP_076160419.1 | NZ_MQUQ01000006.1 | butyrolactone | Chlorothricin (4%) |

| | | | | |
|---|---|---|---|---|
| Amycolatopsis coloradensis | WP_076160419.1 | NZ_MQUQ00000000 (NZ_MQUQ01000006.1) | butyrolactone | Chlorothricin (4%) |
| Actinocrispum wychmicini | WP_132116054.1 | NZ_SLWS01000003.1 | PKS-like,T1PKS | Chlorothricin (48%) |
| Nocardia suismassiliense S-137 | WP_107655957.1 | NZ_LT985361.1 | NRPS,T1PKS,ladderane | Coelimycin (29%) |
| Nocardia mexicana | WP_068019852.1 | NZ_QQAZ01000001 | butyrolactone/ladderane | Colabomycin (4%) |
| Nocardia sp. BMG51109 | WP_024802972.1 | NZ_JAFQ01000004.1 | T1PKS | Stambomycin (40%) |
| Streptomyces peucetius subsp. caesius ATCC 27952 | WP_017584471.1 | NZ_CP022438.1 | T1PKS,T2PKS | Oligomycin (61%) |
| Actinosynnema mirum ATCC 29888 | WP_015803413.1 | NC_013093.1 | T1PKS | Cyclizidine (41%) |
| Streptomyces puniciscabiei | WP_069777509.1 | NZ_CP017248.1 | T1PKS | E-837 (100%) |
| Lechevalieria aerocolonigenes | WP_051784425.1 | NZ_BBOJ01000031.1 | arylpolyene,butyrolactone | Enduracidin (4%) |
| Saccharothrix sp. NRRL B-16348 | WP_053716783.1 | NZ_LGED01000125.1 | NRPS | Erythrochelin (85%) |
| Saccharothrix carnea | WP_106615758.1 | NZ_PYAX01000004.1 | NRPS | Erythrochelin (85%) |
| Saccharothrix texasensis | WP_123742396.1 | NZ_RJKM01000001.1 | NRPS | Erythrochelin (85%) |
| Streptomyces sp. P3 | WP_107448820.1 | NZ_CP028369.1 | LAP,PKS-like,T1PKS,T2PKS | Hedamycin (43%) |
| Kitasatospora griseola | WP_043910458.1 | NZ_JXZB01000002.1 | NRPS,PKS-like,T1PKS,T2PKS | Hedamycin (31%) |
| Kitasatospora sp. CB02891 | WP_100586111.1 | NZ_NNBO01000004.1 | NRPS,PKS-like,T1PKS,T2PKS | Hedamycin (34%) |
| Streptomyces sp. FBKL.4005 | WP_059247715.1 | NZ_NPKF01000001.1 | LAP,PKS-like,T1PKS,T2PKS | Hedamycin (43%) |
| Streptomyces reticuli | WP_059247715.1 | NZ_LN997842.1 | LAP,PKS-like,T2PKS | Hedamycin (46%) |
| Actinocorallia populi strain A251 | WP_106396532.1 | NZ_PVZV01000001.1 | PKS-like,T1PKS,T2PKS | Hedamycin (56%) |
| Streptomyces sp. t99 | WP_030720926.1 | NZ_NTGQ01000069.1 | T1PKS,T2PKS | Hedamycin (59%) |
| Streptomyces sp. st140 | WP_030720926.1 | NZ_NTGS01000037.1 | T2PKS | Hedamycin (59%) |

| | | | | |
|---|---|---|---|---|
| Streptomyces sp. f51 | WP_030720926.1 | NZ_NTHH01000040.1 | T1PKS,T2PKS | Hedamycin (62%) |
| Thermomonospora curvata ATCC 19995 | WP_012853108.1 | NC_013510.1 | PKS-like,T1PKS,T2PKS | Hedamycin (81%) |
| Streptomyces sp. ms115 | WP_097947834.1 | NZ_NTHD01000048.1 | PKS-like,T1PKS,T2PKS | Hedamycin (87%) |
| Streptomyces sp. b62 | WP_030720926.1 | NZ_NTHK01000012.1 | PKS-like,T1PKS,T2PKS | Hedamycin (87%) |
| Streptomyces sp. f150 | WP_030720926.1 | NZ_NTHG01000036.1 | PKS-like,T1PKS,T2PKS | Hedamycin (87%) |
| Streptomyces griseus subsp. griseus NRRL F-5144 | WP_030720926.1 | NZ_JOGA01000026.1 | PKS-like,T1PKS,T2PKS | Hedamycin (87%) |
| Streptomyces sp. gb14 | WP_030720926.1 | NZ_NTHF01000021.1 | PKS-like,T1PKS,T2PKS | Hedamycin (87%) |
| Streptomyces sp. SS07 | WP_030720926.1 | NZ_KZ268499.1 | PKS-like,T1PKS,T2PKS,terpene | Hedamycin (87%) |
| Streptomyces rubellomurinus subsp. indigoferus | KJS54098.1 | JZKG00000000.1 | T1PKS | Hedamycin (9%) |
| Streptomyces sp. NRRL S-146 | WP_031110854.1 | NZ_JOAW01000493.1 | T2PKS | Hedamycin (9%) |
| Streptomyces sp. Ag82_G6-1 | WP_097222862.1 | NZ_OCNA01000001.1 | PKS-like, T1PKS, T2PKS | Hedamycin (90%) |
| Nocardia alba DSM 44684 | TCJ89880.1 | SMFR01000008.1 | PKS-like,T1PKS,T2PKS | Hedamycin (25%) |
| Nocardia alba NBRC 108234 | WP_067458355.1 | NZ_BDAX01000033.1 | PKS-like,T1PKS,T2PKS | Hedamycin (25%) |
| Streptomyces sp. NP10 | WP_126932507.1 | NZ_PDIQ01000024.1 | PKS-like,T1PKS,T2PKS | Hedamycin (50%) |
| Streptomyces sp. JS01 | WP_030720926.1 | NZ_JPWW01000020.1 | PKS-like,T1PKS,T2PKS | Hedamycin (87%) |
| Streptomyces sp. Root264 | KRD19112.1 | NZ_LMIZ01000001 | LAP,PKS-like,T1PKS,T2PKS | Hedamycin (43%) |
| Actinophytocola oryzae DSM 45499 | WP_133905254.1 | NZ_SOCP01000009.1 | T1PKS | Incednine (17%) |
| Streptomyces sp. WAC 01529 | WP_125514820.1 | CP029617.1 | T1PKS/NRPS | Lorneic acid A (23%) |
| uncultured bacterium | ASV46999.1 | KY560362.1 | T1PKS | Lorneic acid A (23%) |
| Streptomyces oceani | WP_070197325.1 | NZ_LJGU00000000.1 | T1PKS | Lorneic acid A (23%) |

| | | | | |
|---|---|---|---|---|
| Streptomyces puniciscabiei | WP_069776466.1 | NZ_CP017248.1 | T1PKS,terpene | Lorneic acid A (28%) |
| Streptomyces sp. DS1-2 | WP_120696069.1 | NZ_RBDY01000003.1 | T1PKS | Methymycin / pikromycin (57%) |
| Sorangium cellulosum | KYF78568.1 | JEMB01002769.1 | T1PKS | Micromonolactam (100%) |
| Actinosynnema pretiosum strain X47 | WP_096495686.1 | NZ_CP023445.1 | T1PKS | Microtermolide A (33%) |
| Streptomyces sp. CNQ-509 | WP_047018908.1 | NZ_CP011492.1 | NRPS,PKS-like,T1PKS | Microtermolide A (53%) |
| Streptomyces sp. AZ1-7 | WP_120696069.1 | NZ_RBDX01000002.1 | T1PKS | Nocardiopsin (21%) |
| Catenulispora acidiphila DSM 44928 | WP_015795553.1 | NC_013131.1 | T1PKS/NRPS | Octacosamicin (29%) |
| Streptomyces sp. NRRL S-118 | WP_031080613.1 | NZ_KL591043.1 | T1PKS | Piericidin A1 (58%) |
| Streptomyces griseus subsp. griseus NRRL F-5144 | WP_030723159.1 | NZ_JOGA01000040.1 | NRPS | Polyoxypeptin (40%) |
| Streptomyces phaeoluteigriseus strain DSM 41896 | OJT46852.1 | MPOH00000000.2 | NRPS,T1PKS,other | Polyoxypeptin (75%) |
| Streptomyces sp. XY006 | WP_094055126.1 | NZ_NOKT01000017.1 | NRPS,other | Polyoxypeptin (40%) |
| Streptomyces sp. E5N91 s-91 | WP_121712942.1 | NZ_RAIE01000715.1 | other | Polyoxypeptin (40%) |
| Pseudonocardiaceae bacterium YIM PH 21723 | WP_120088448.1 | NZ_QZFT00000000.1 | T1PKS | Pyrronazol B (9%) |
| Nocardia altamirensis NBRC 108246 | WP_069164184.1 | NZ_BDAY01000046.1 | T1PKS | Stambomycin (40%) |
| Streptomyces sp. NRRL WC-3742 | WP_031071135.1 | NZ_JOCF01000031.1 | NRPS,T1PKS | Lydicamycin (40%) |
| Kitasatospora azatica KCTC 9699 | WP_083976688.1 | NZ_JQMO01000003.1 | PKS-like,T1PKS | Zincophorin (61%) |
| Streptomyces rubellomurinus ATCC 31215 | WP_017584471.1 | NZ_JZKH01000090.1 | N/A | N/A |
| Streptomyces albidoflavus | WP_128462586.1 | NZ_QHCQ00000000.1 | N/A | N/A |

| | | | | |
|---|---|---|---|---|
| | | (NZ_SCDQ01000064 .1) | | |
| Streptomyces sp. PRh5 | WP_051573751.1 | NZ_JABQ01000071. 1 | N/A | N/A |
| Streptomyces sp. FXJ1.172 | WP_067044802.1 | NZ_LWRP01000068. 1 | N/A | N/A |
| Streptomyces sp. AVP053U2 | WP_062189972.1 | NZ_LMTQ02000008. 1 | N/A | N/A |
| Enhygromyxa salina | KIG11693.1 | JMCC00000000.2 | N/A | N/A |
| Streptomyces tricolor NRRL B-16925 | WP_086702637.1 | NZ_MUMF00000000. 1 | N/A | N/A |
| Streptomyces sp. 8K308 | WP_132929808.1 | NZ_SMKC01000043. 1 | N/A | N/A |
| Mycobacteroides abscessus | WP_079869619.1/SHS51726.1 | NZ_FSAT01000004.1 | N/A | N/A |
| Streptomyces atriruber | WP_055564739.1 | NZ_LIPN01000084.1 (NZ_SMKI00000000. 1) | N/A | N/A |
| Streptomyces hainanensis DSM 41900 | WP_132818863.1 | NZ_SMKI01000162.1 | N/A | N/A |
| Nocardiopsis valliformis DSM 45023 | WP_017584471.1 | NZ_ANAZ01000058. 1 | N/A | N/A |
| Streptomyces sp. E2N166 | WP_121750360.1 | NZ_RAIF01000032.1 | N/A | N/A |
| Streptomyces sp. NBRC 110035 | WP_042171193.1 | NZ_BBNN01000027. 1 | N/A | N/A |
| Plesiocystis pacifica SIR-1 | WP_006969759.1/EDM81215.1 | NZ_ABCS01000005. 1 | transAT-PKS | N/A |
| Nonomuraea sp. SBT364 | WP_049575489.1 | NZ_LAVL01000205.1 | T1PKS | N/A |

Abbreviations: Polyketide synthase (PKS), type I polyketide synthase (T1PKS), type II polyketide synthase (T2PKS), nonribosomal peptide synthetase (NRPS), lassopeptide (LAP

### Supplementary Figures

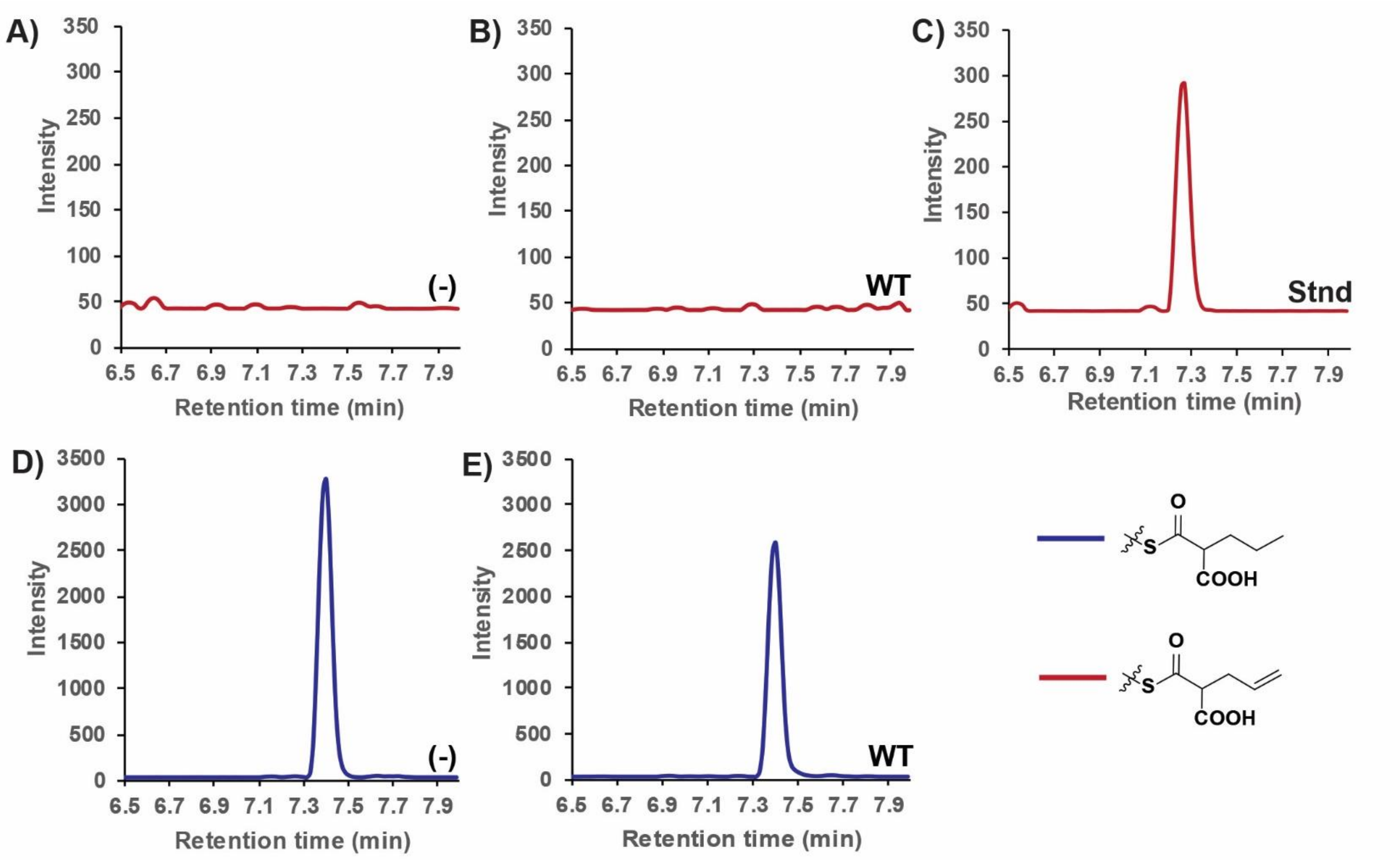

**Figure S1**. TcsD activity on propylmalonyl-ACP (pathway *A2* from Figure 1) analyzed via targeted LC-MS/MS. WT = wild type TcsD, (-) = negative control, Stnd = standard. The chromatograms shown depict: **A)** the allylmalonyl-ACP transition for the negative control reaction **B)** the allylmalonyl-ACP transition for the wild type TcsD reaction **C) a**n allylmalonyl-ACP standard **D)** the propylmalonyl-ACP transition for the negative control reaction **E)** the propylmalonyl-ACP transition for the TcsD wild type reaction

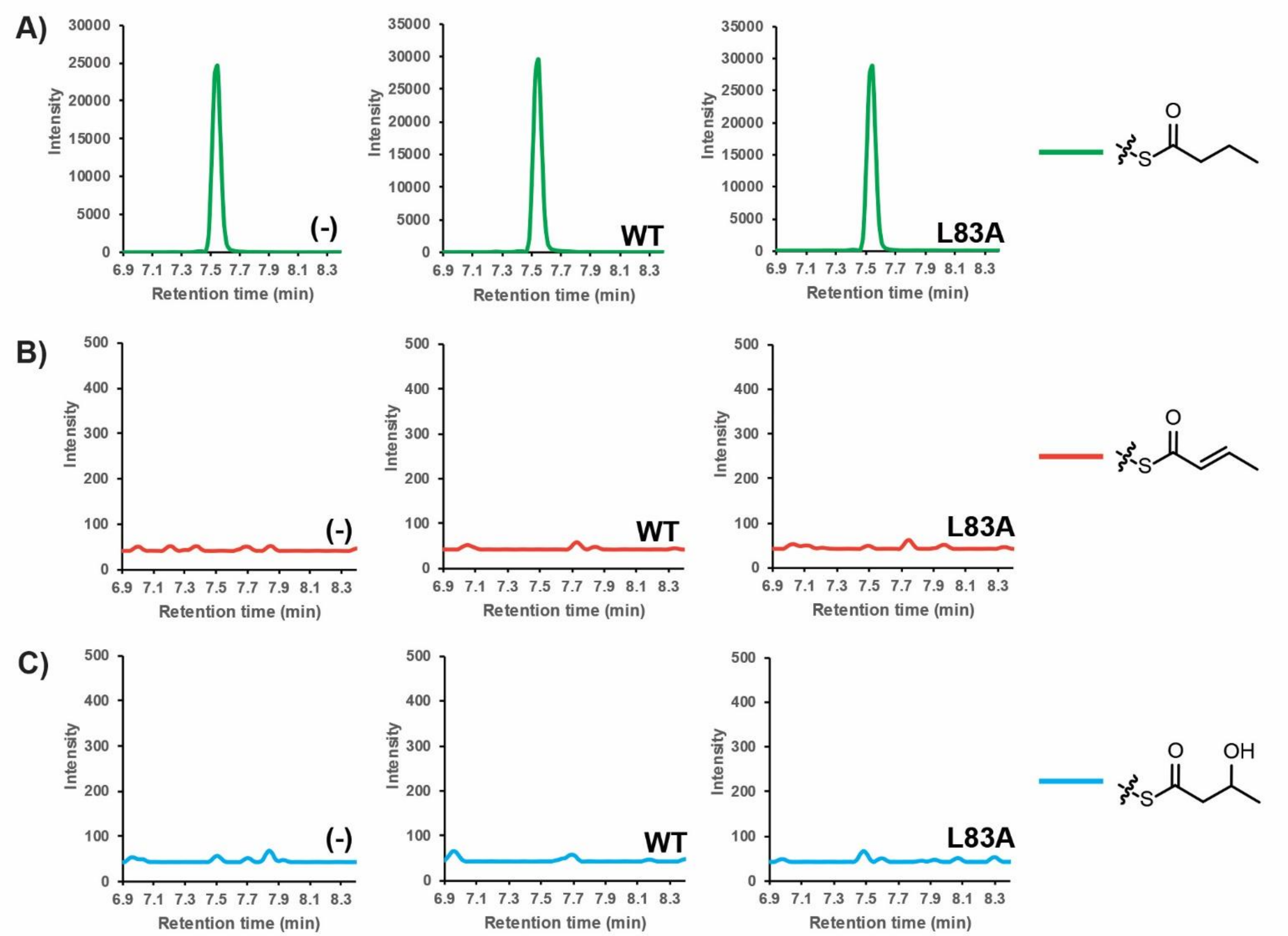

**Figure S2**. Activity of TcsD wild type and L83A mutant on butyryl-ACP analyzed by targeted LC-MS/MS. WT = wild type TcsD, (-) = negative control, L83A = TcsD L83A mutant. Chromatograms representing different transitions are color coded (key is shown on right side of figure). The chromatograms shown depict: **A)** butyryl-ACP, **B)** crotonyl-ACP, or **C)** 3-hydroxybutyryl-ACP for each assay (negative control, TcsD wild type, or TcsD L83A).

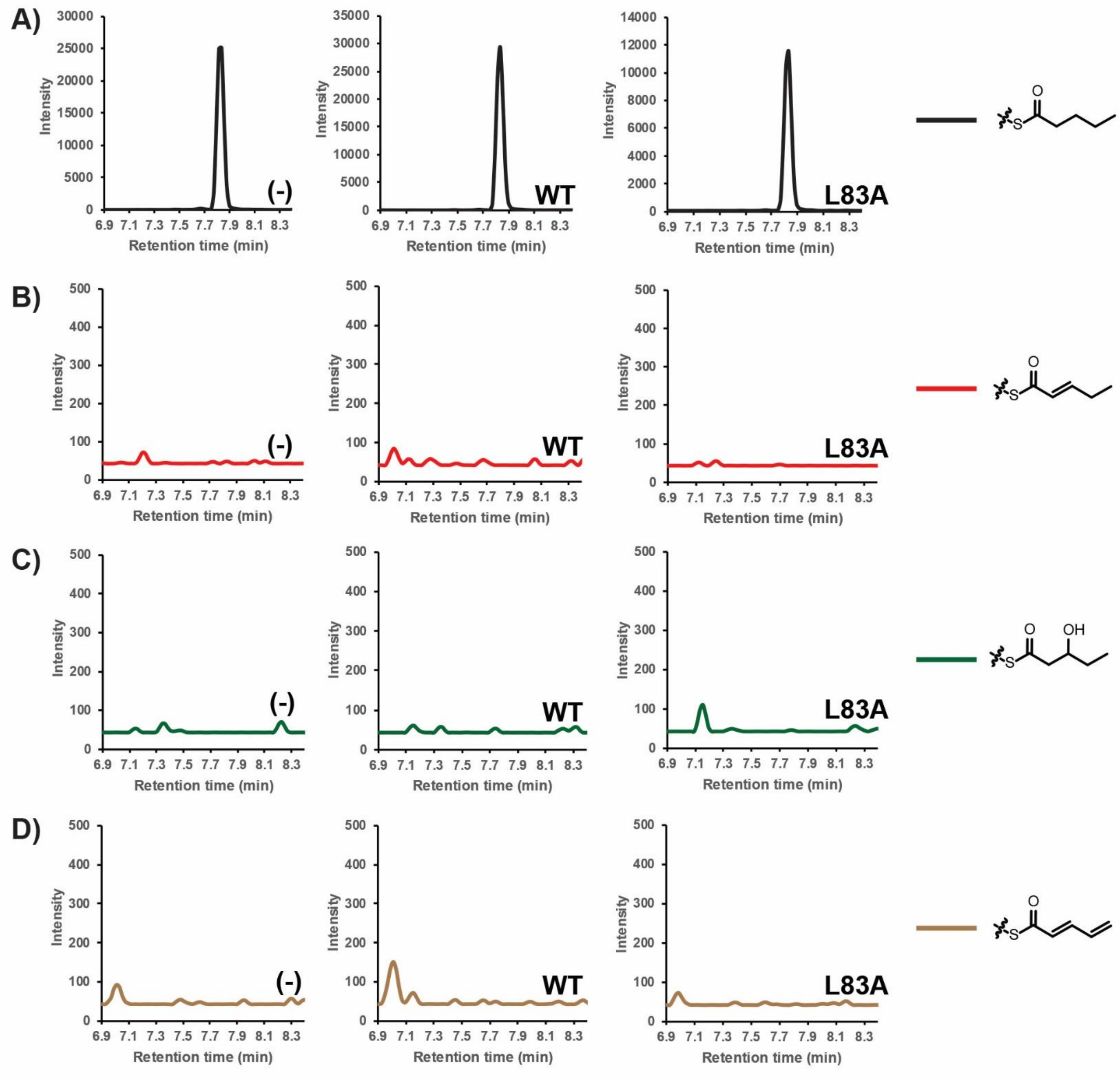

**Figure S3**. Activity of TcsD wild type and L83A mutant on pentanoyl-ACP analyzed by targeted LC-MS/MS. WT = wild type TcsD, (-) = negative control, L83A = TcsD L83A mutant. Chromatograms representing different transitions are color coded (key is shown on right side of figure). The chromatograms shown depict: **A)** pentanoyl-ACP, **B)** 2-pentenoyl-ACP, **C)** the 3-hydroxypentanoyl-ACP, or **D)** 2,4-pentadienoyl-ACP for each assay (negative control, TcsD wild type, or TcsD L83A).

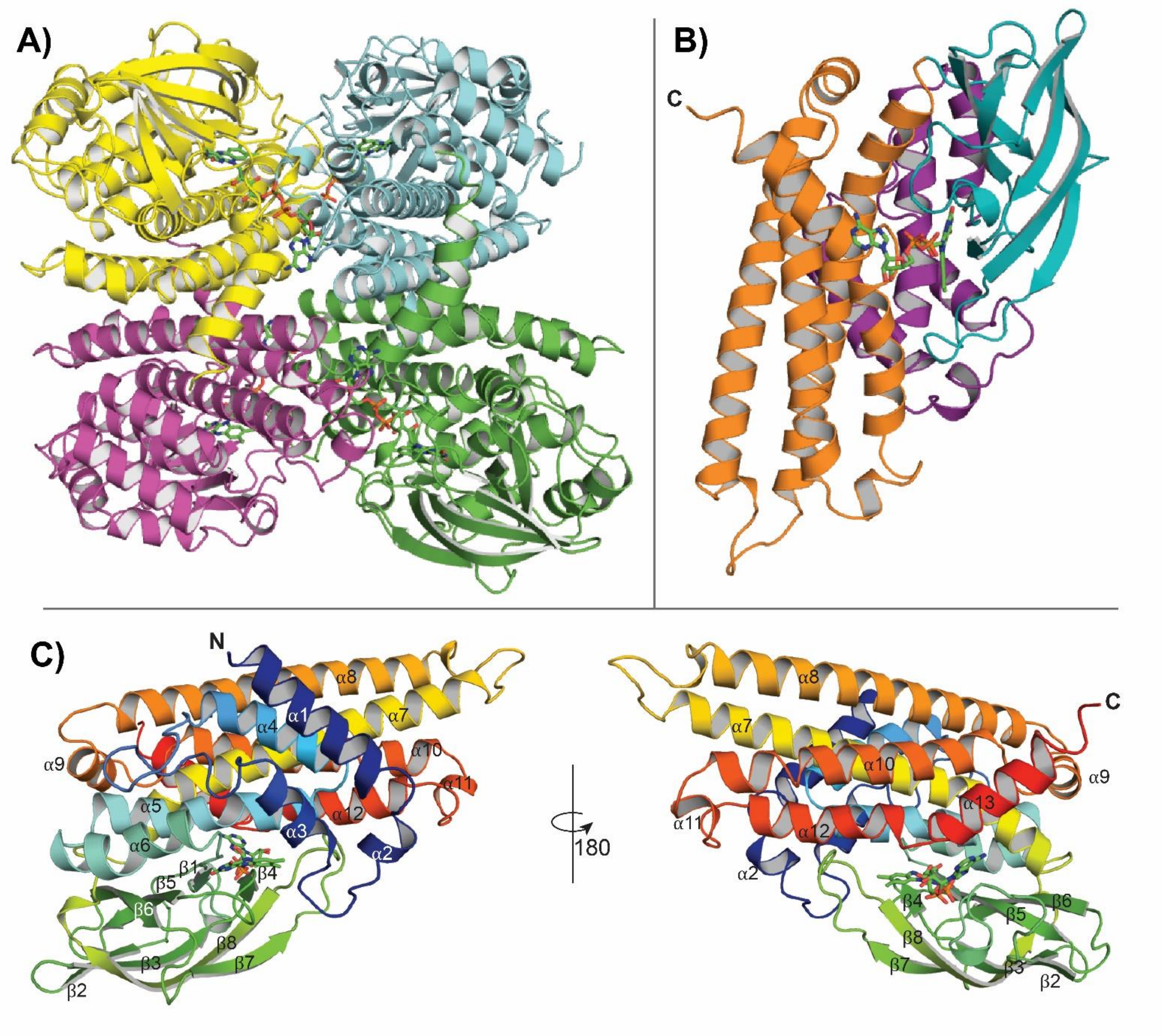

**Figure S4**. Overall structure of TcsD. **A)** Tetrameric structure of TcsD. Each subunit is shown in a different color. FAD cofactors are shown as sticks. **B)** A single TcsD subunit. The N-terminal α-helix domain (purple), middle β-sheet domain (teal), and C-terminal α-helix domain (orange) are highlighted. C) A single TcsD monomer colored in a progressive rainbow from its N- (blue) to C-terminus (red). Helices and sheets are numbered.

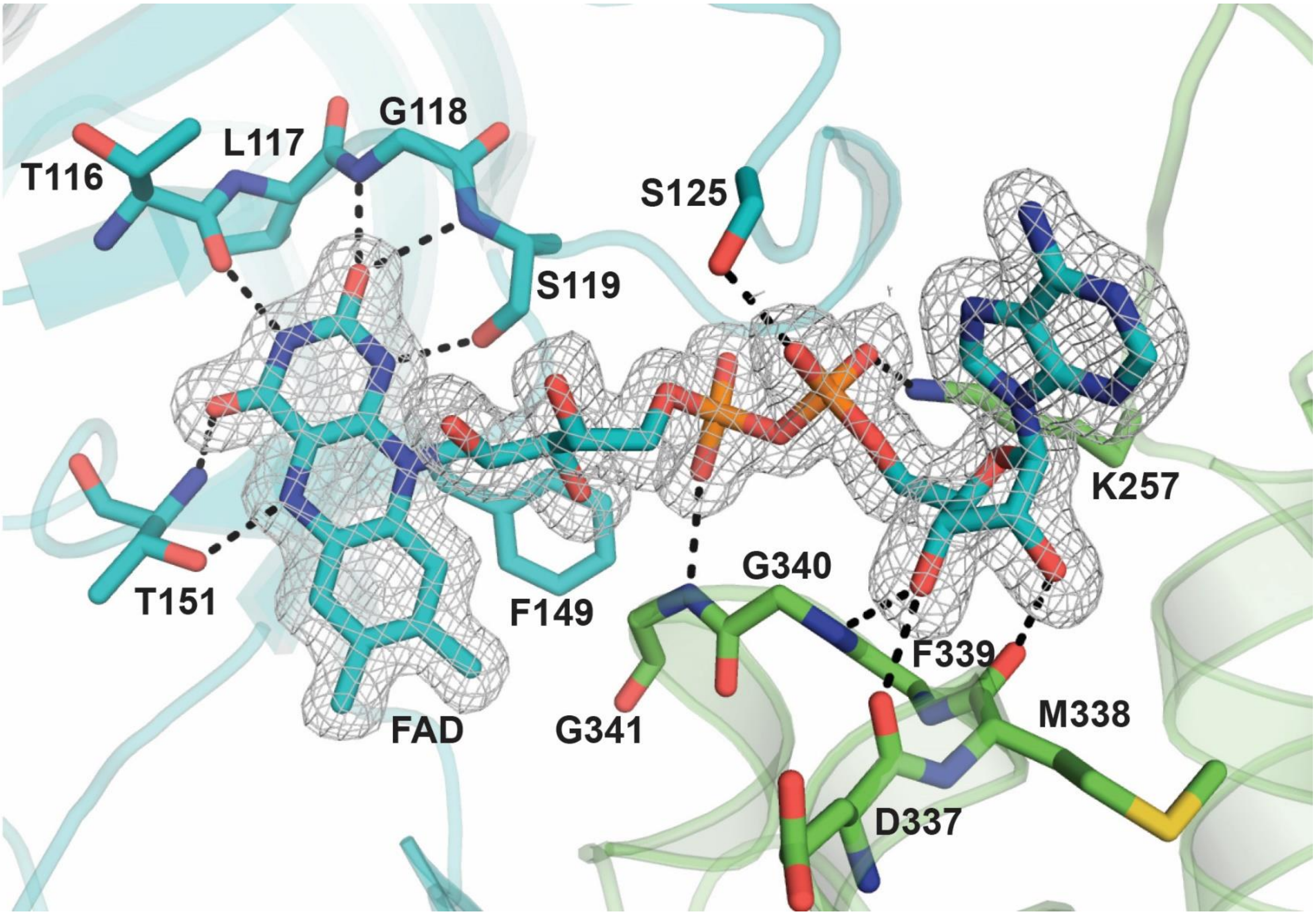

**Figure S5**.FAD-interacting residues of TcsD. The $2F_o - F_c$ electron density map of FAD is shown with gray mesh and is contoured at 1σ. Residues that interact with FAD via hydrogen bonds are shown as sticks, with hydrogen bonds depicted as black dashed lines. Residues pertaining to the same subunit as FAD (T116, L117, G118, S119, S125, F149, T151) are shown in blue, while residues pertaining to the adjacent subunit (K257, D337, M338, F339, G340, G341) are shown in green.

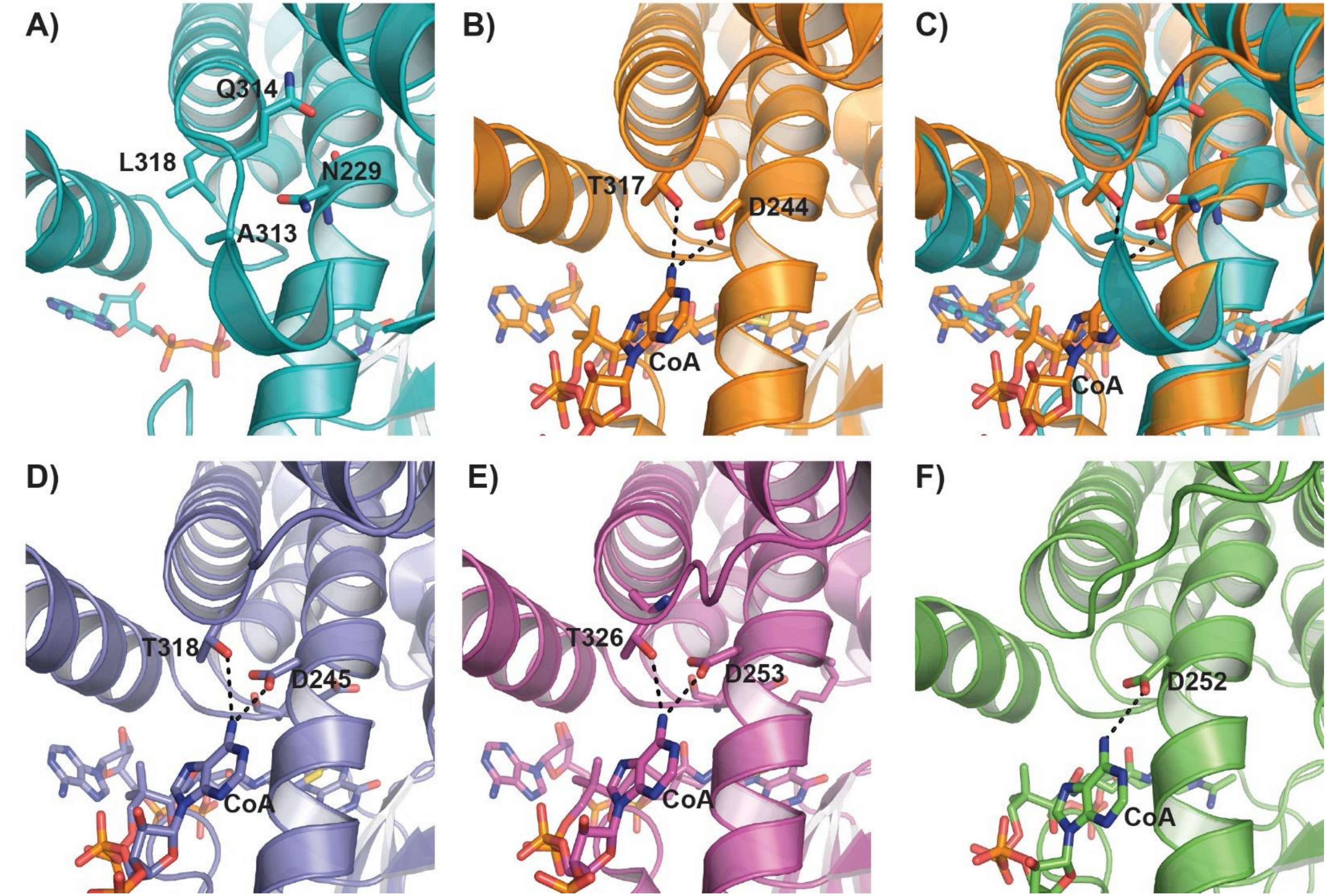

**Figure S6**. Residues of ACADs that hydrogen bond with the nucleotide portion of CoA in **A)** TcsD, **B)** *M. esldenii* butyryl-CoA dehydrogenase (1buc), **C)** *M. esldenii* butyryl-CoA dehydrogenase overlaid with TcsD, **D)** *Sus scrofa* medium chain ACAD (1udy), **E)** rat short chain ACAD (1jqi), **F)** human isovaleroyl-CoA dehydrogenase (1ivh). TcsD has an Asn residue in the place of the conserved Asp of α,β-ACADs. Helix 9 occupies some of the space where the nucleotide portion of CoA would normally bind. The conserved Thr residue is replaced with a Leu residue.

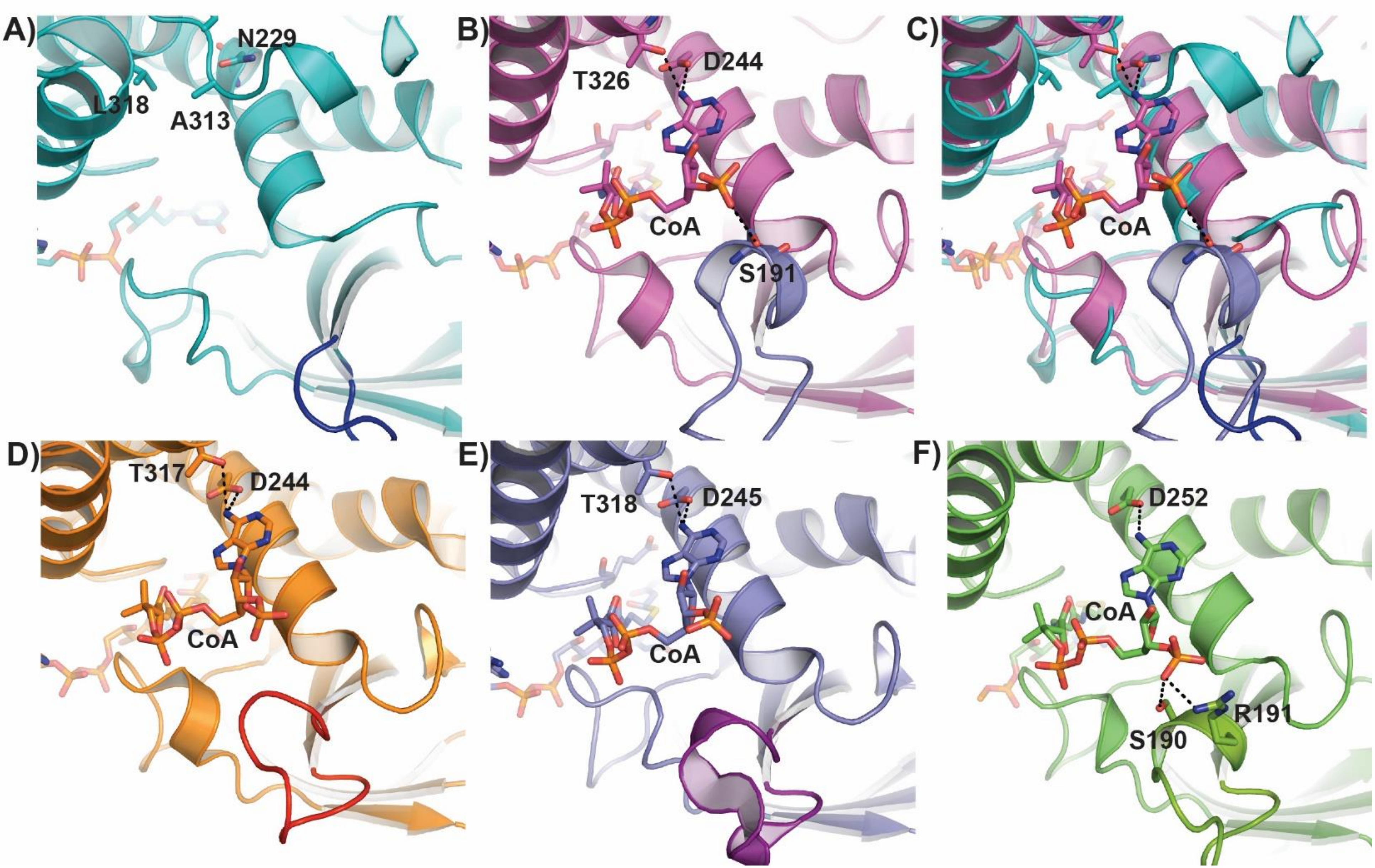

**Figure S7**.CoA phosphate-binding region of **A)** TcsD, **B)** rat short chain ACAD (1jqi), **C)** rat short-chain ACAD overlaid with TcsD, **D)** *M. esldenii* butyryl-CoA dehydrogenase (1buc), **E)** *Sus scrofa* medium chain ACAD (1udy), **F)** human isovaleroyl-CoA dehydrogenase (1ivh). The loop that approaches the phosphate groups is colored in a different shade in each panel.

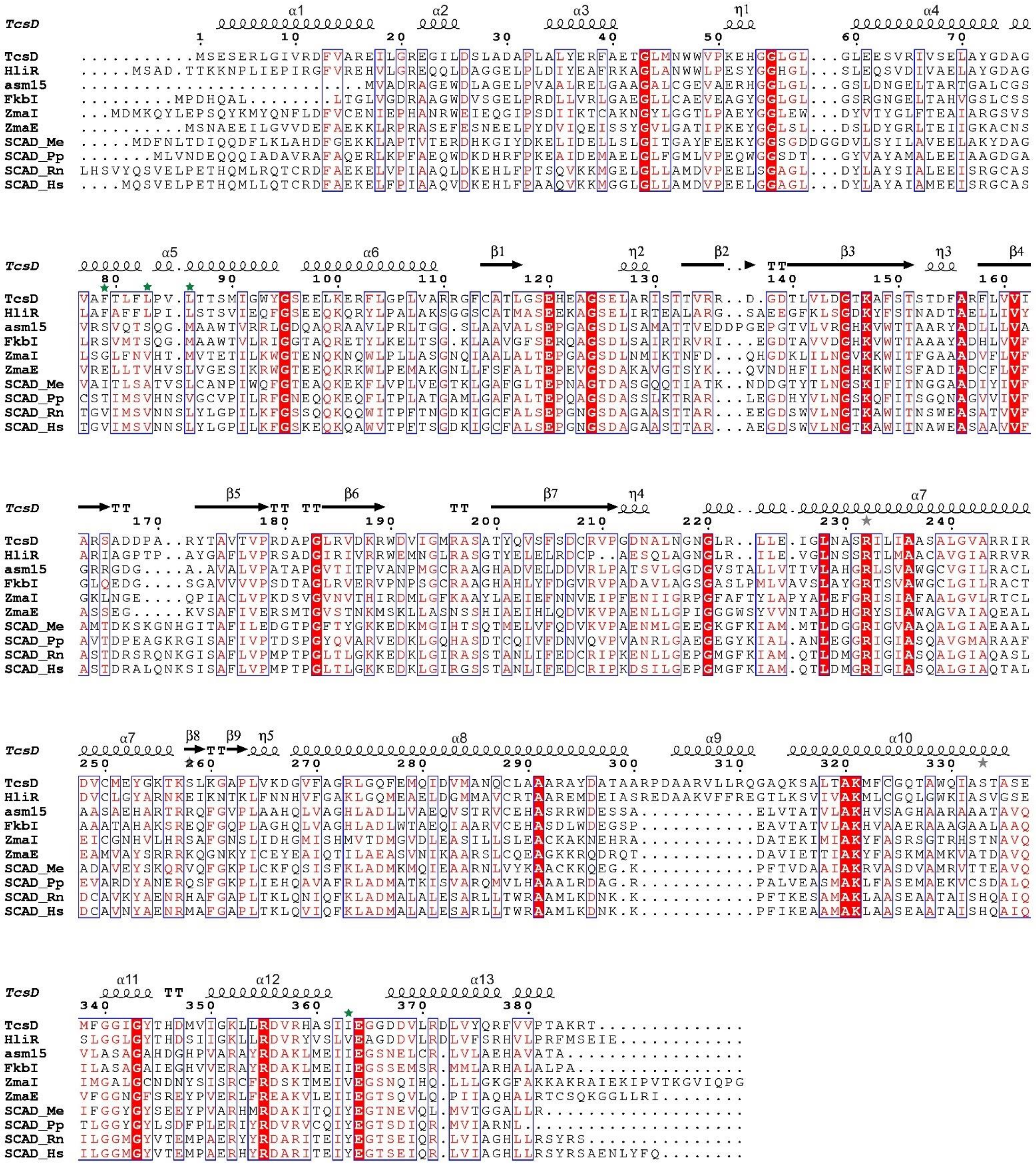

**Figure S8**. Sequence alignment of TcsD with homologs. Structural elements noted above the alignment are based on the TcsD structure. Proteins included in the alignment and their Genbank accession numbers are: γ,δ-ACADs TcsD (ADU56309.1) and HliR (AMM72019.1), hydroxymalonate semialdehyde dehydrogenases asm15 (AAM54093.1), FkbI (TAI41666.1), and ZmaE (AAD40109.1), aminomalonate semialdehyde dehydrogenase ZmaI (AAR87758.1), and α,β-ACADs SCAD_Me (pdb 1buc), SCAD_Pp (NP_744365.1), SCAD_Rn (pdb 1jqi), and SCAD_Hs (pdb 2vig). Green stars denote residues that form the active site pocket.

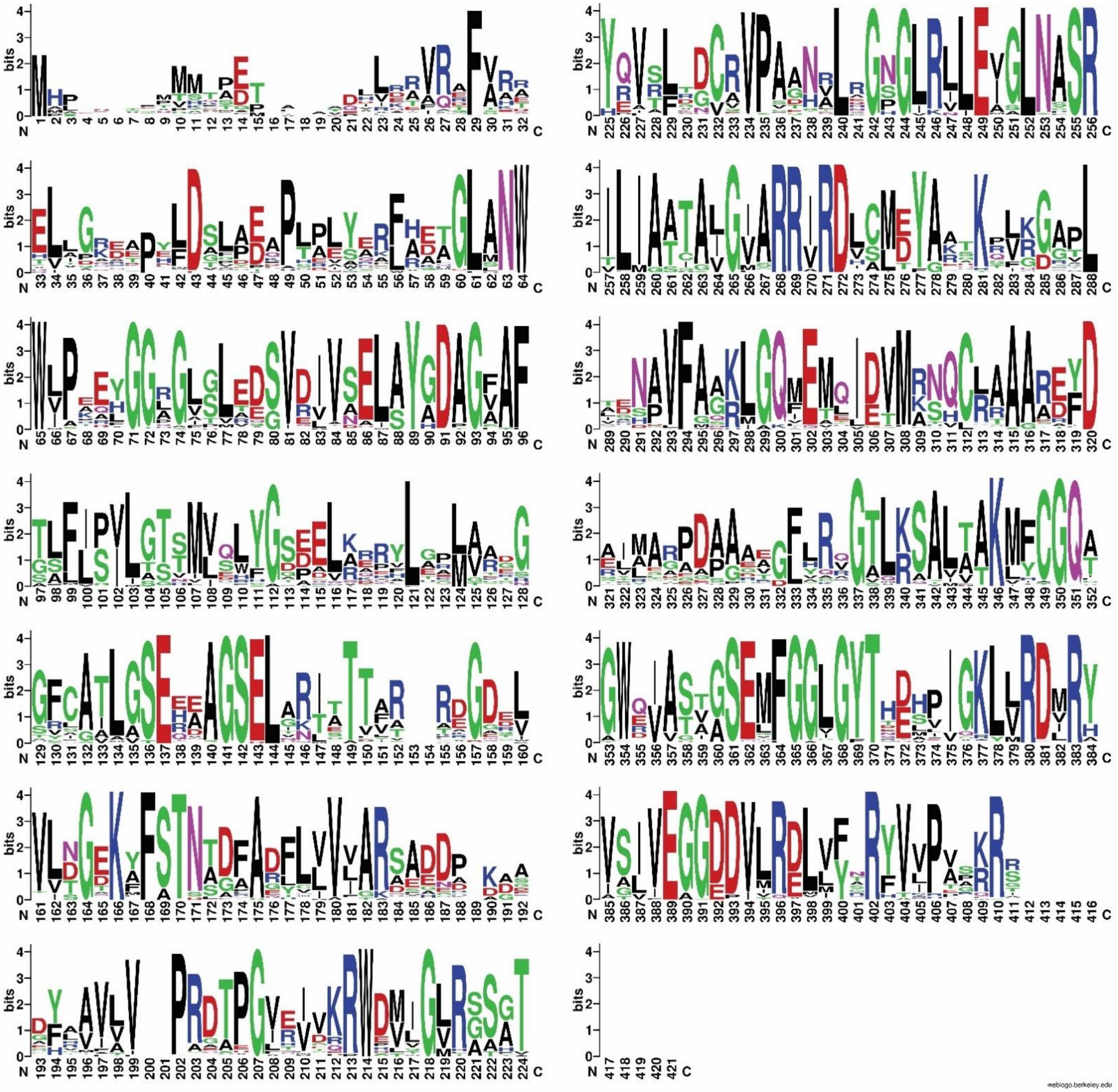

**Figure S9**. Web logo depicting the conservation of amino acids across a multiple sequence alignment of all γ,δ-ACADs identified in this work. Larger-sized letters represent higher conservation, while smaller letters represent less conserved amino acids.

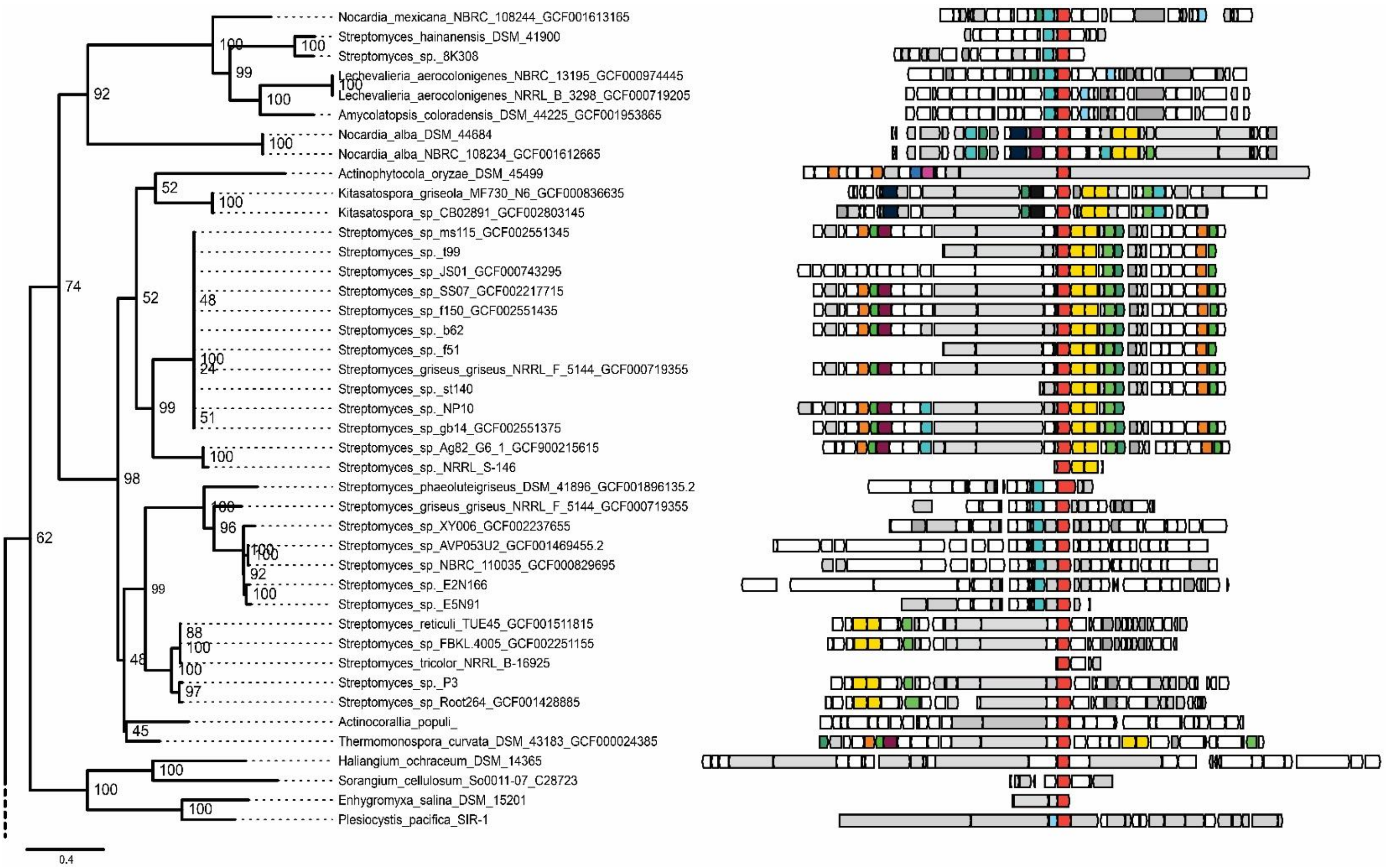

**Figure S10A**. Phylogenetic tree and genomic contexts of γ,δ-ACADs identified in this work. This figure represents the top portion of the tree and connects to the bottom portion (Figure S10B).

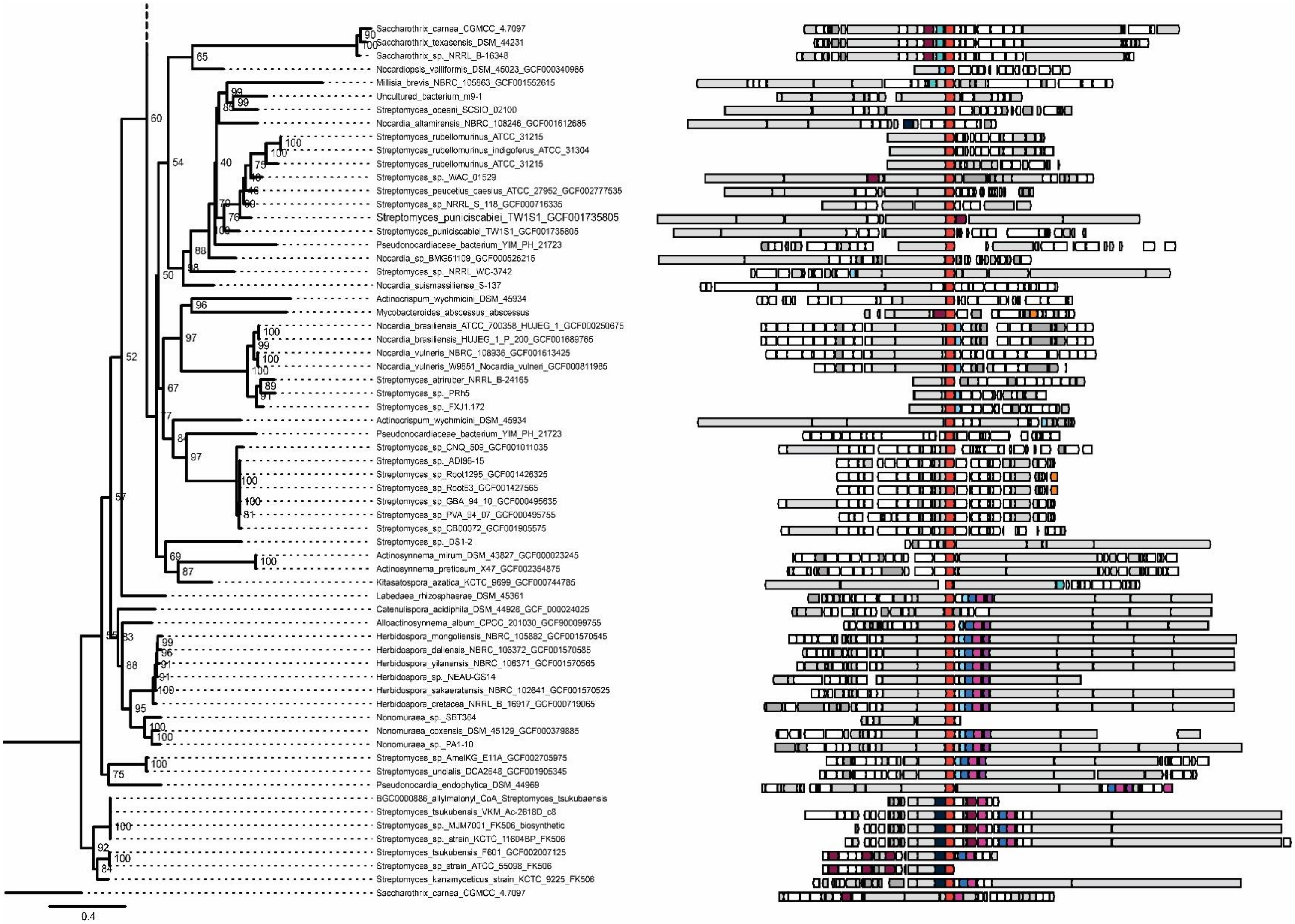

**Figure S10B**. Phylogenetic tree and genomic contexts of γ,δ-ACADs identified in this work. This figure represents the bottom portion of the tree and connects to the top portion (Figure S10A).

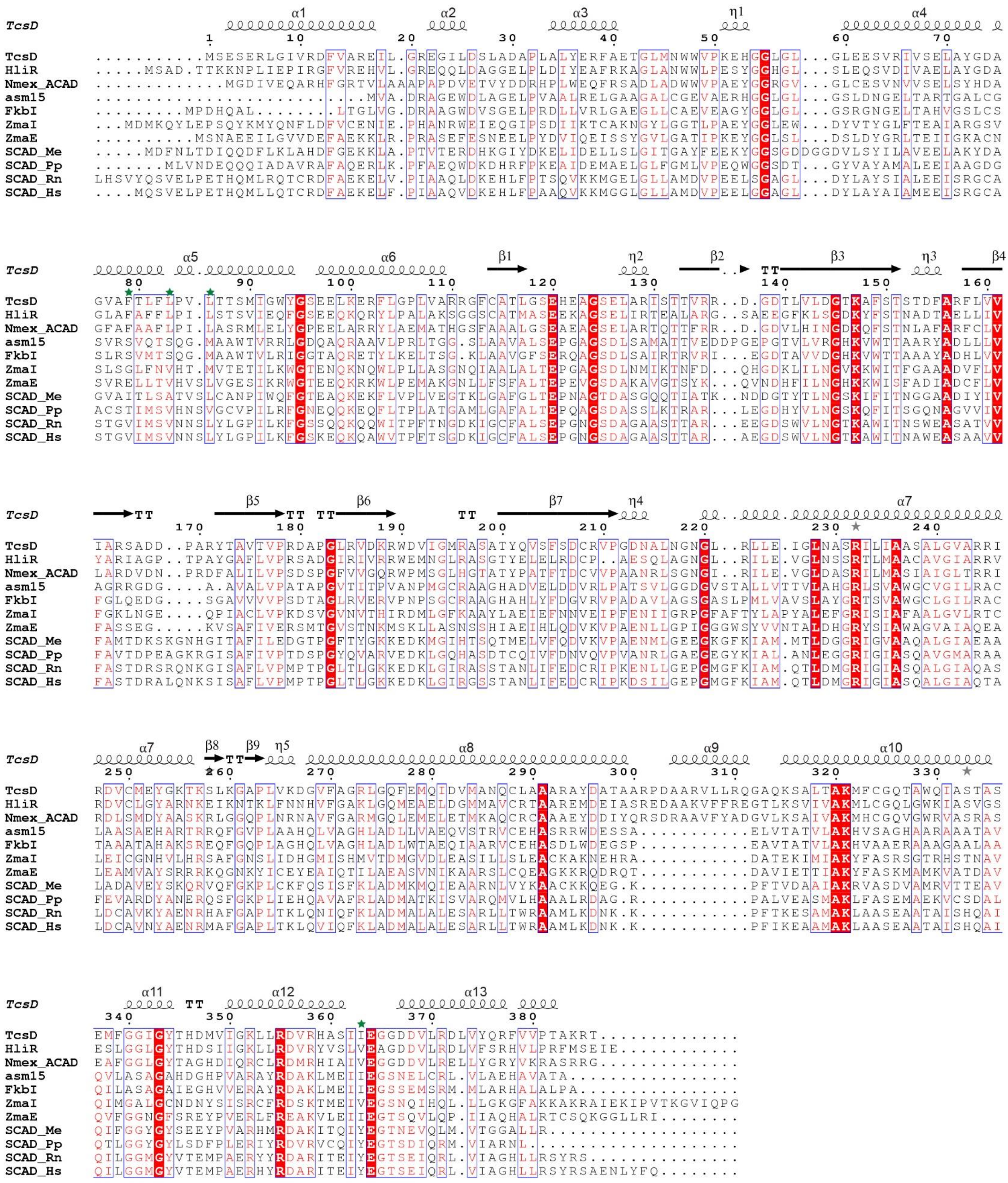

**Figure S11**. Sequence alignment of TcsD with homologs, including Nmex-ACAD. Structural elements are based on the TcsD structure, and green stars denote residues that line the active site pocket. Proteins included in the alignment and their Genbank accession numbers are listed in the description for Figure S8.

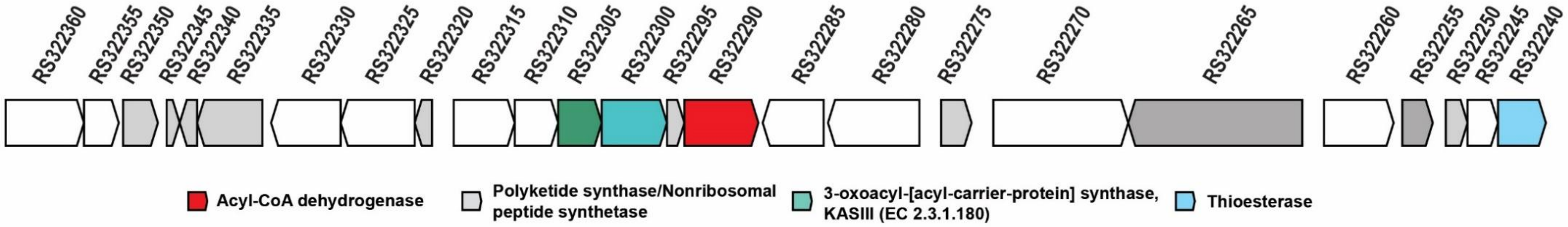

| Locus tag | Gene product | GenBank Protein ID |
|---|---|---|
| DFR68_RS32240 | thioesterase | WP_068019833.1 |
| DFR68_RS32245 | hypothetical protein | WP_068019835.1 |
| DFR68_RS32250 | hypothetical protein | WP_068019837.1 |
| DFR68_RS32255 | response regulator transcription factor | WP_068019839.1 |
| DFR68_RS32260 | histidine kinase | WP_068019841.1 |
| DFR68_RS32265 | AAA family ATPase | WP_068019843.1 |
| DFR68_RS32270 | MMPL family transporter | WP_068019846.1 |
| DFR68_RS32275 | ester cyclase | WP_084519643.1 |
| DFR68_RS32280 | MFS transporter | WP_068019850.1 |
| DFR68_RS32285 | hypothetical protein | WP_114699758.1 |
| DFR68_RS32290 | acyl-CoA dehydrogenase | WP_068019852.1 |
| DFR68_RS32295 | acyl carrier protein | WP_068019854.1 |
| DFR68_RS32300 | ketoacyl-ACP synthase III | WP_084519644.1 |
| DFR68_RS32305 | SDR family oxidoreductase | WP_068019857.1 |
| DFR68_RS32310 | HAD family phosphatase | WP_068019859.1 |
| DFR68_RS32315 | hypothetical protein | WP_084519645.1 |
| DFR68_RS32320 | acyl carrier protein | WP_068019863.1 |
| DFR68_RS32325 | beta-ketoacyl-[acyl-carrier-protein] synthase family protein | WP_068019865.1 |
| DFR68_RS32330 | hypothetical protein | WP_068019867.1 |
| DFR68_RS32335 | DUF2236 domain-containing protein | WP_068019869.1 |
| DFR68_RS32340 | hypothetical protein | WP_068019872.1 |
| DFR68_RS32345 | hypothetical protein | WP_114699759.1 |
| DFR68_RS32350 | SRPBCC family protein | WP_068019876.1 |
| DFR68_RS32355 | hypothetical protein | WP_068019878.1 |
| DFR68_RS32360 | MCE family protein | WP_084519646.1 |

**Figure S12**. Genomic context of the *Nocardia mexicana* γ,δ-acyl-CoA dehydrogenase (Nmex-ACAD) and predicted gene products for each coding sequence.

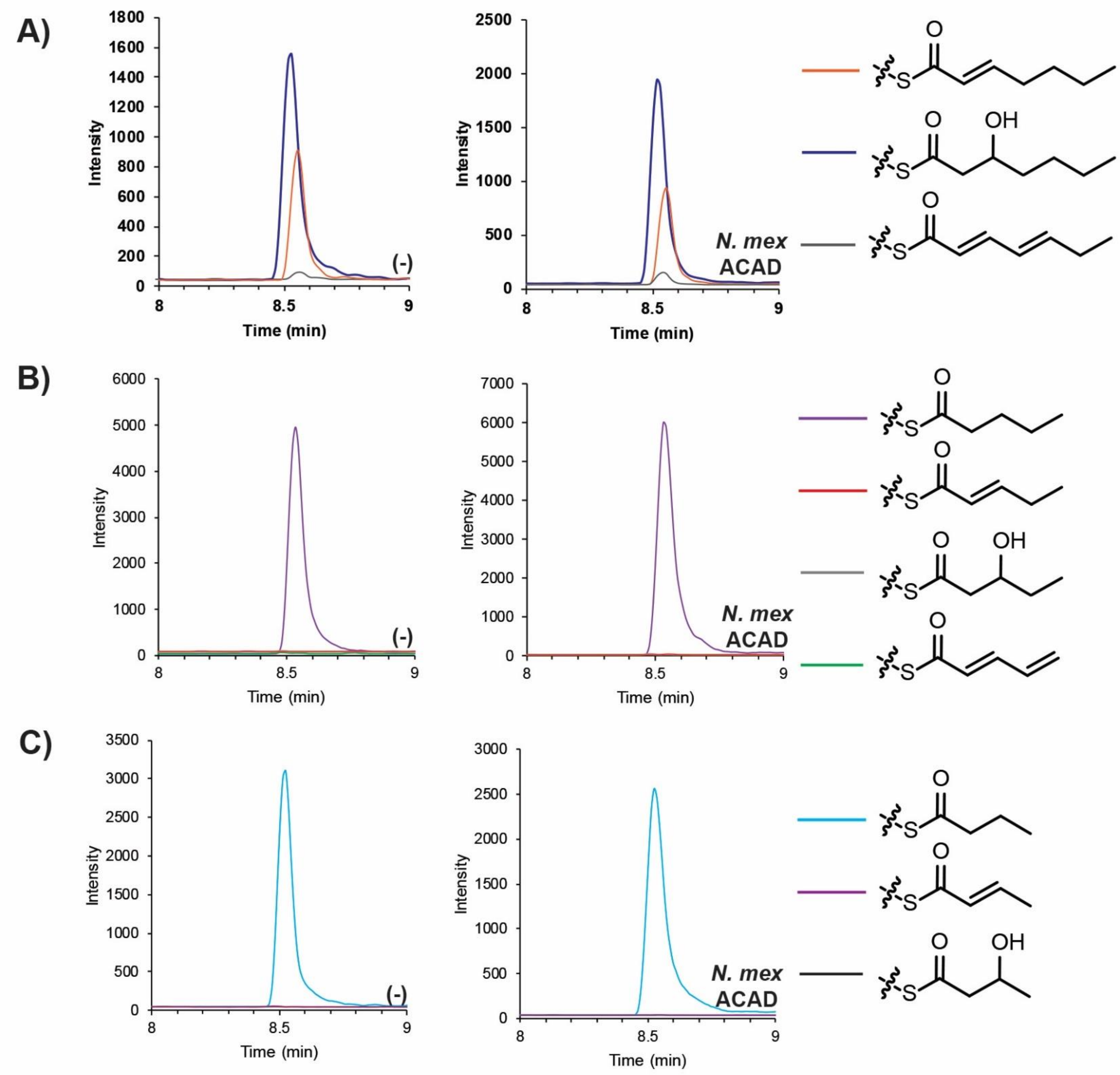

**Figure S13**. Activity of Nmex-ACAD analyzed by targeted LC-MS/MS. *N.mex* ACAD = wild type Nmex-ACAD, (-) = negative control. Chromatograms representing different transitions are color coded (key is shown on right side of figure). The chromatograms shown depict assays on the following substrates: **A)** 2-heptenoyl-ACP, **B)** pentanoyl-ACP, or **C)** butyryl-ACP. For all samples, only the substrate (no product) was observed.

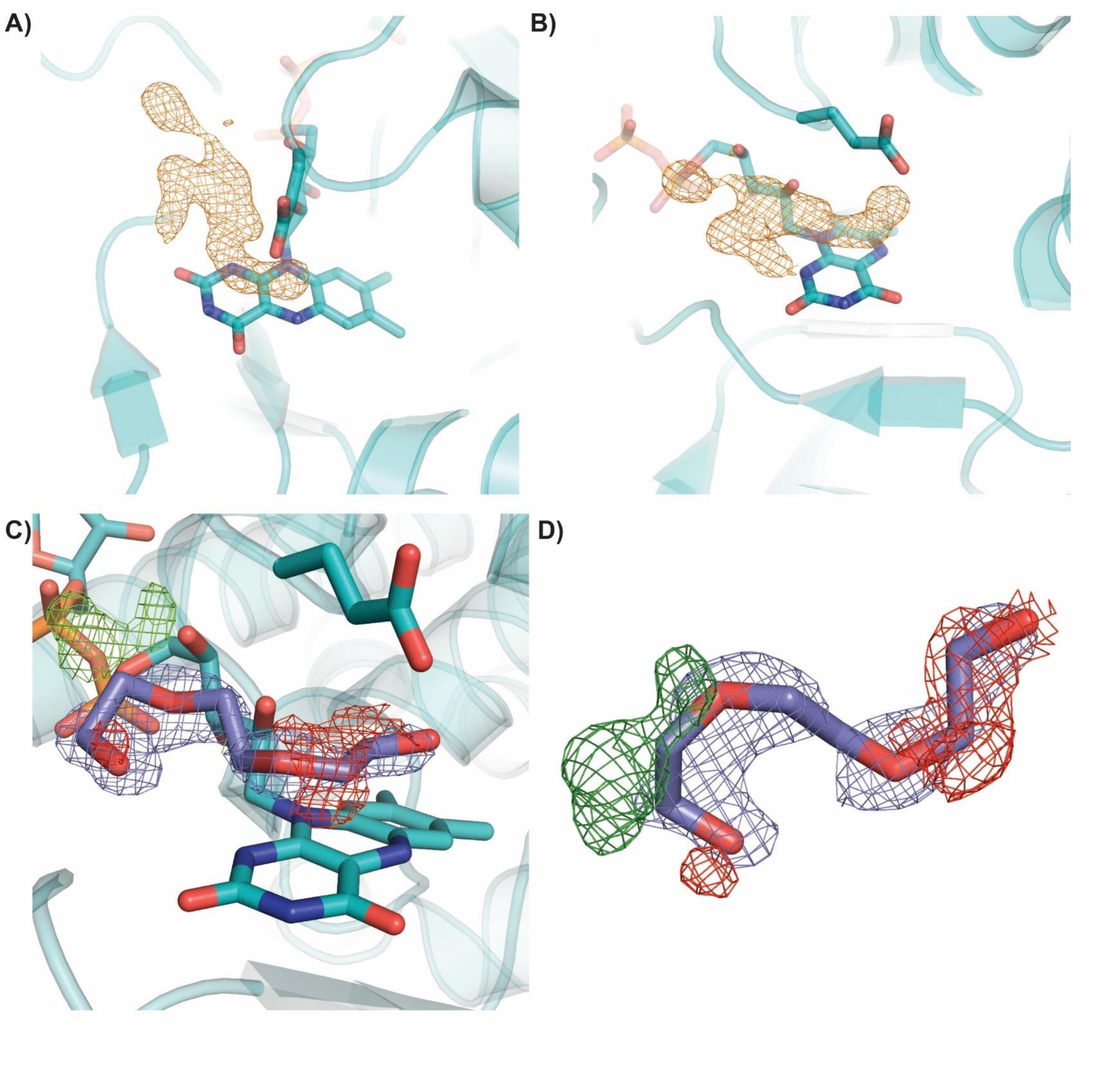

**Figure S14**. Unidentified density in active site of TcsD and modeling of PEG trimer into density. **A) and B)** Shape of the unidentified density (in orange mesh) in the TcsD active site ($F_o - F_c$ map at 3σ) **C) and D)** Model of TcsD after refinement with a PEG trimer (purple sticks) occupying the unknown density, showing that this PEG fragment does not fit properly within the density. The $2F_o - F_c$ map is shown in blue mesh at 1σ. The $F_o - F_c$ map is shown in green at 3σ and in red at -3σ.

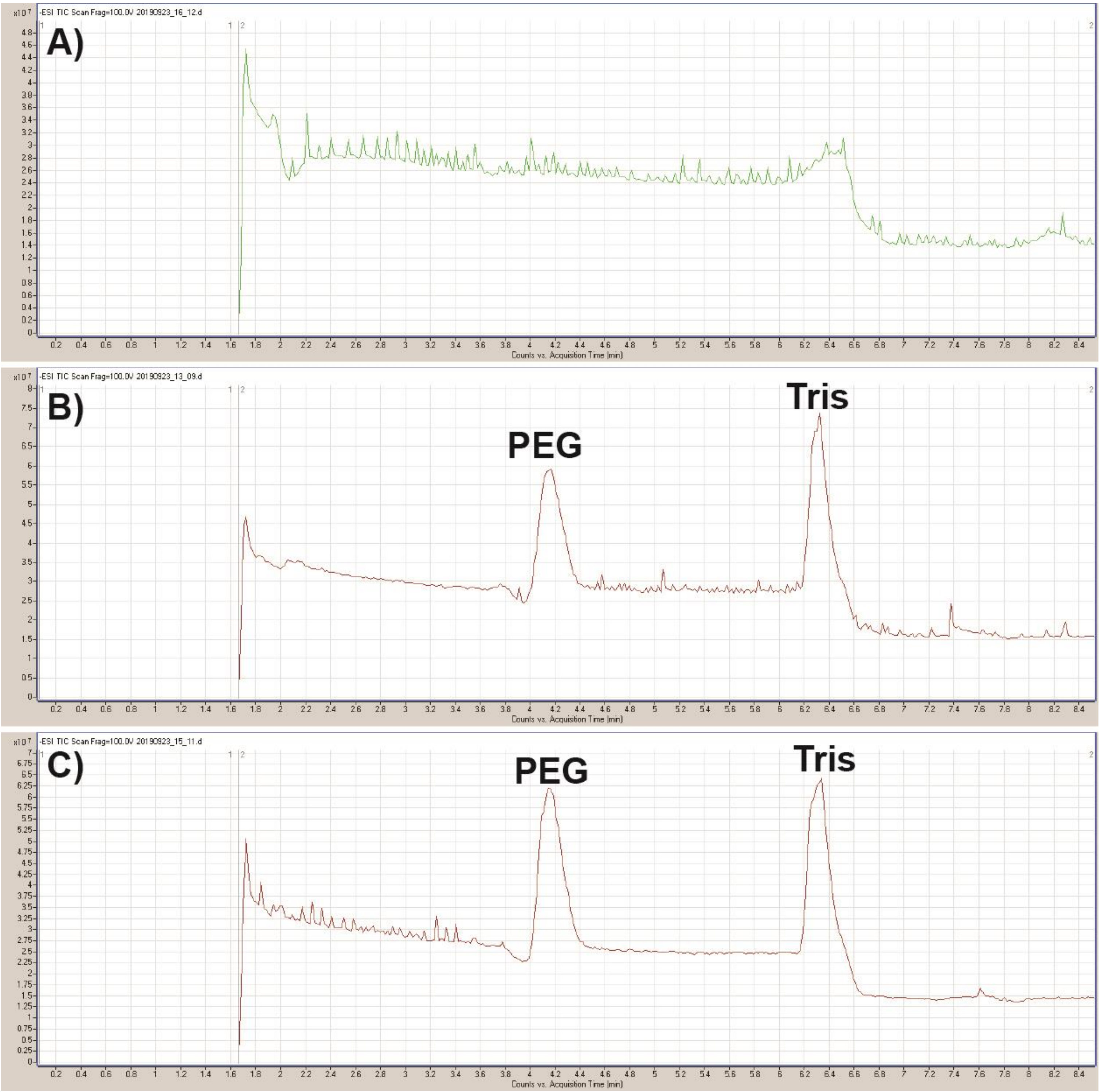

**Figure S15**. High resolution untargeted LC-TOF analysis of denatured purified TcsD samples in negative ion mode. Samples are as follows: **A)** Crystallization buffer **B)** Supernatant from boiled TcsD **C)** Supernatant from acetonitrile-denatured TcsD. Peaks pertaining to PEG (a common LCMS contaminant) and Tris are labeled.

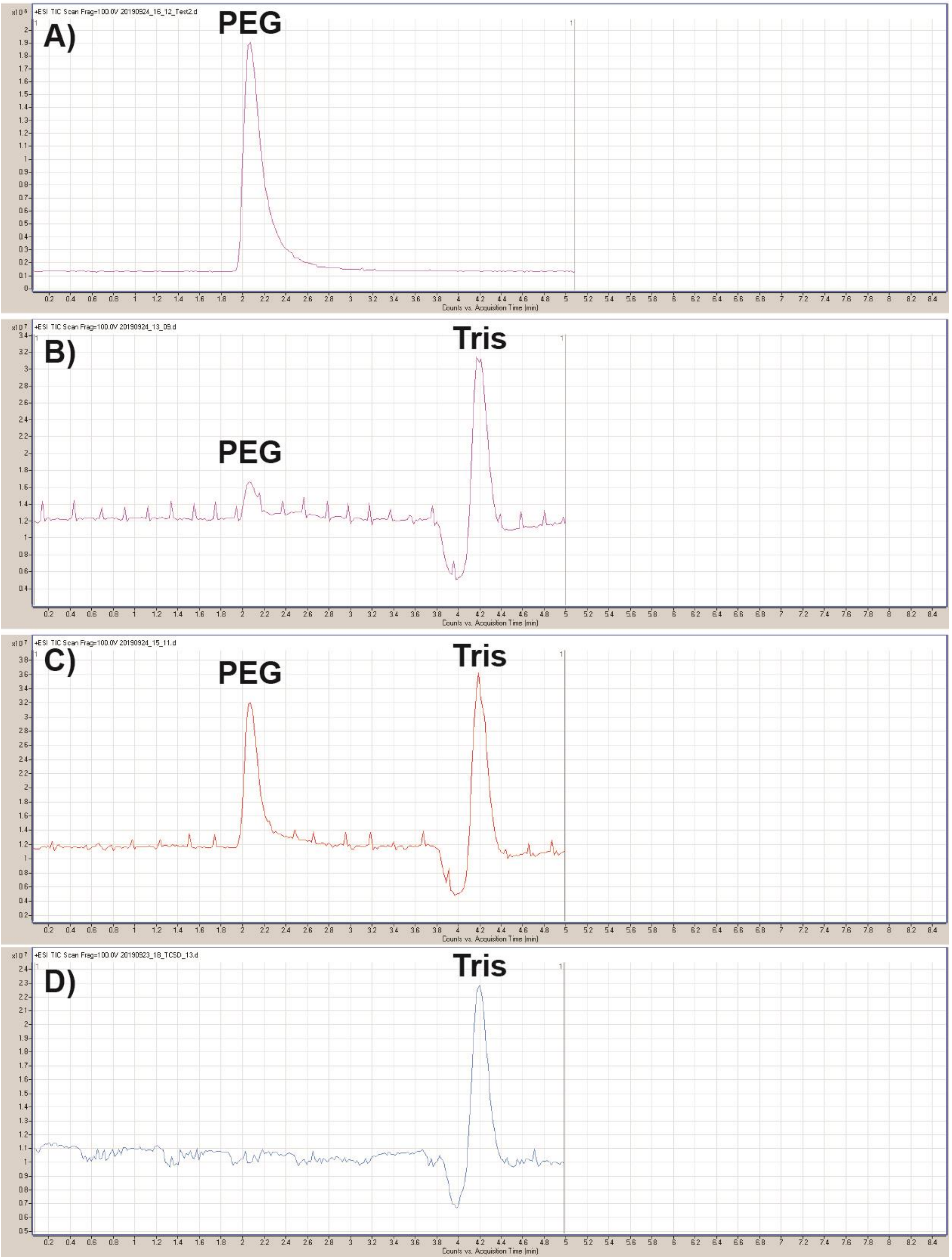

**Figure S16**. High resolution untargeted LC-TOF analysis of denatured purified TcsD samples in positive ion mode. Samples are as follows: **A)** Crystallization buffer **B)** Supernatant from boiled TcsD **C)** Supernatant from acetonitrile-denatured TcsD, **D)** lysis buffer. Peaks pertaining to PEG (a common LCMS contaminant) and Tris are labeled.

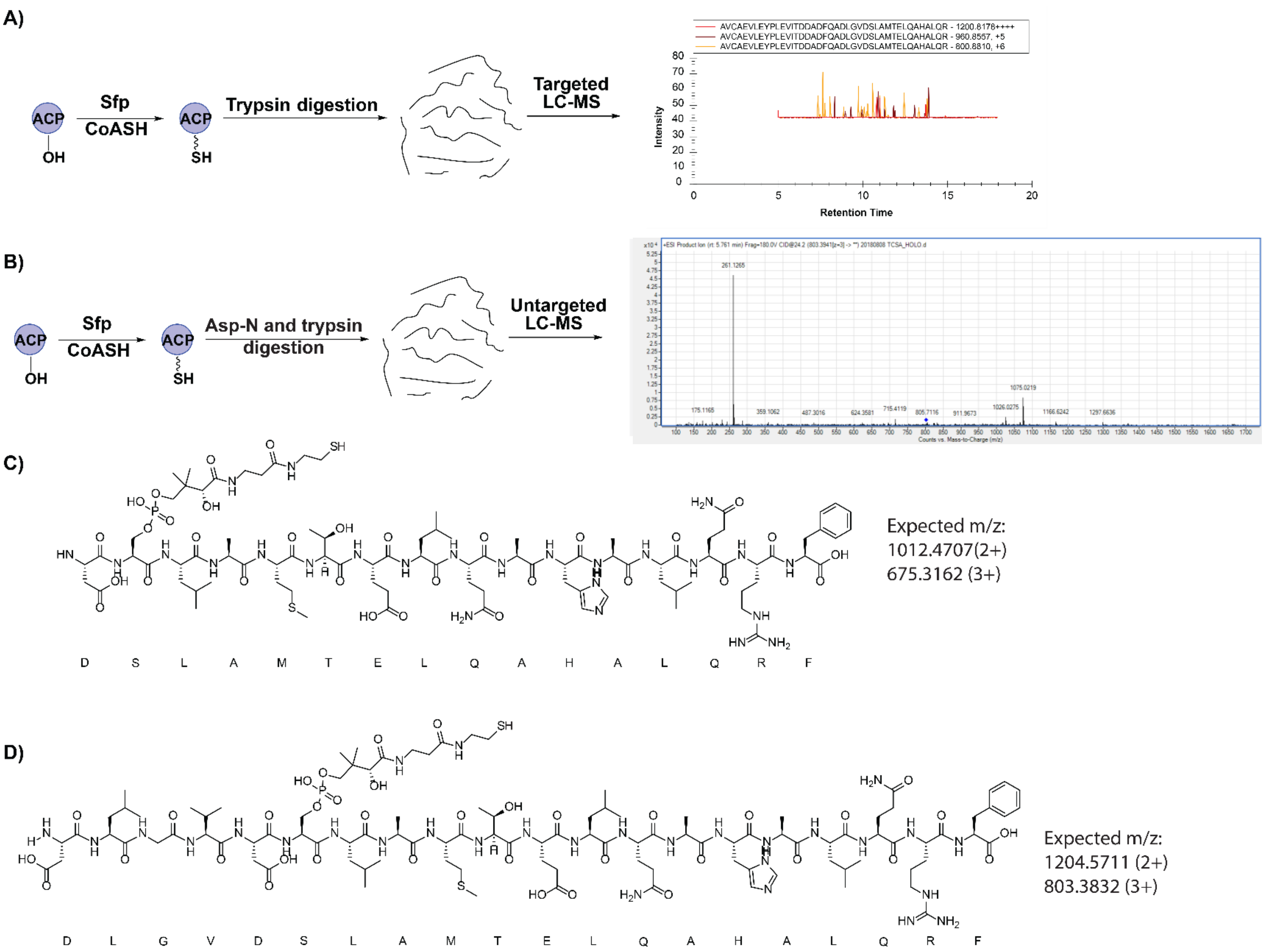

**Figure S17.** Development of a phosphopantetheine ejection method for studying intermediates bound to TcsA-ACP. **A)** Digestion of TcsA with trypsin results in an active site peptide 40 amino acids in length which is not detectable using targeted LC-MS/MS. **B)** Dual digestion of TcsA with Asp-N and trypsin results in a detectable peptide from which phosphopantetheine ejection can be observed. **C)** Predicted TcsA active site peptide resulting from TcsA digestion with Asp-N and trypsin. **D)** Observed TcsA active site peptide resulting from TcsA digestion with Asp-N and trypsin (verified with data in Figure S18).

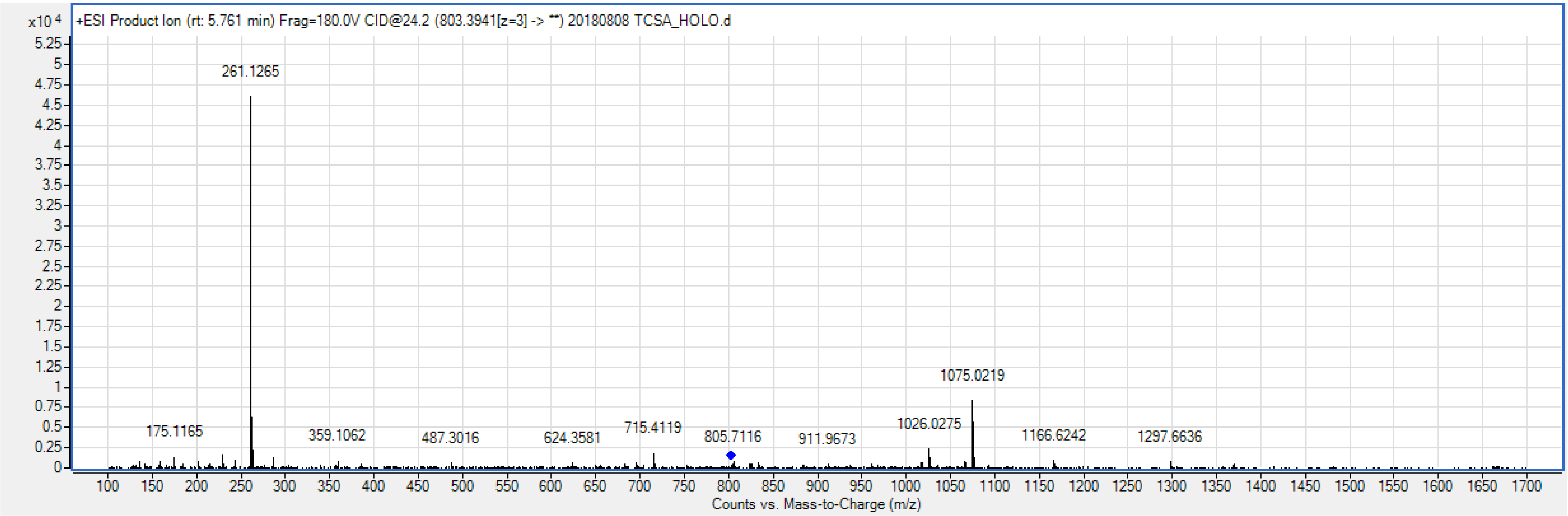

**Figure S18.** Product ion mass spectrum showing phosphopantetheine ejection ion resulting from holo-TcsA digested with Asp-N and trypsin. The product ion is derived from a parent ion with m/z = 803.394 [z=3], representing the peptide “DLGVDSLAMTELQAHALQR” in which Asp-N misses one cleavage, not the expected “DSLAMTELQAHALQR” active site peptide (see Figure S17).

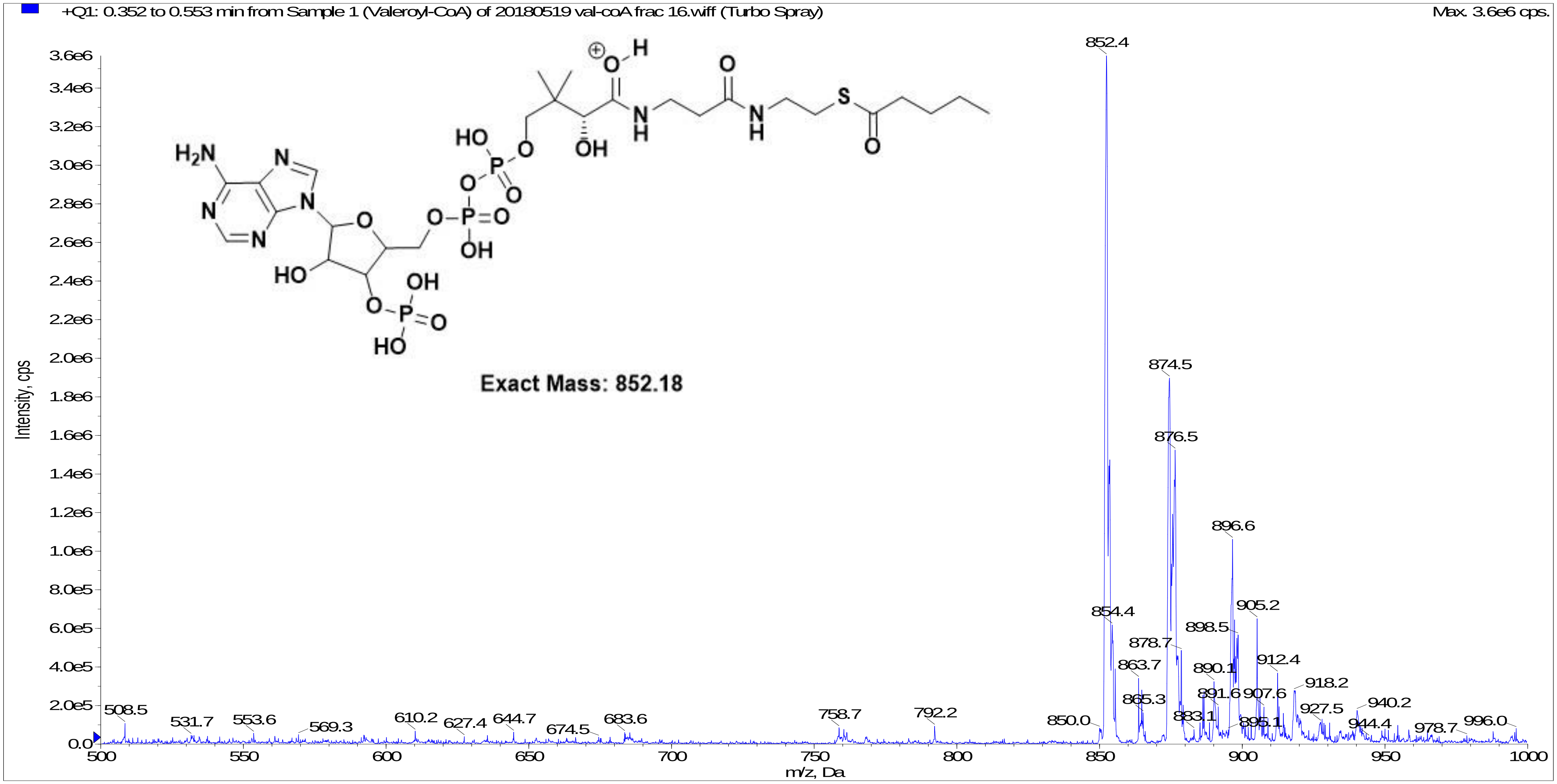

**Figure S19.** Mass spectrum of synthesized pentanoyl-CoA. An m/z of 852.18 [z=1] represents the [M+H] ion, while the m/z of 874.5 [z=1] represents the [M+Na] ion.

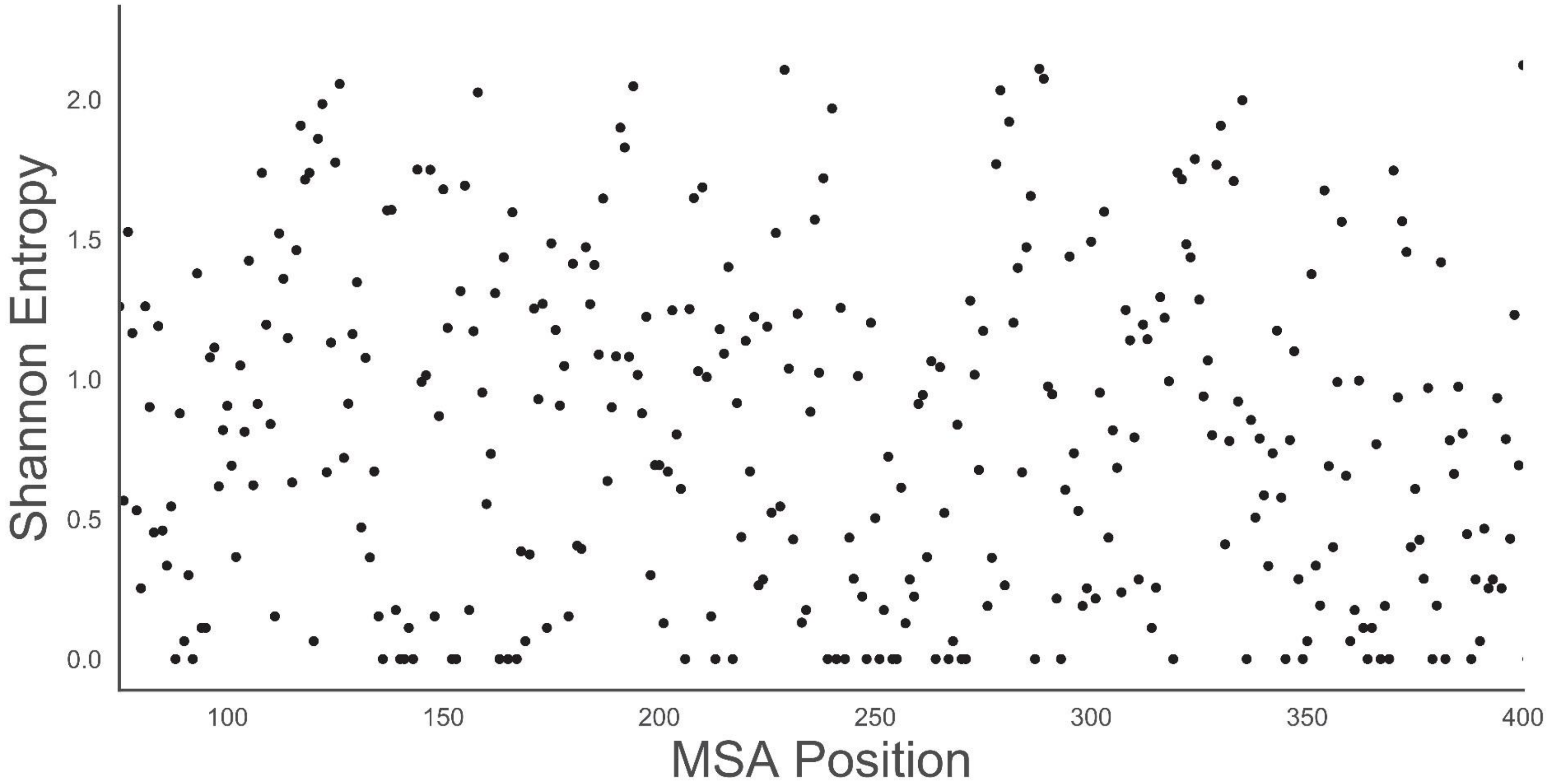

**Figure S20**. Shannon entropy plot depicting amino acid conservation at each position of a multiple sequence alignment of all γ,δ-ACADs identified in this work. An entropy value of 0 represents 100% conservation whereas larger entropy values represent less conservation. This plot was used to generate a positional conservation web logo (Figure S6).

### Data Availability

***Structural data deposition***: The atomic coordinates and structural factors of TcsD have been deposited in the Protein Data Bank, PDB ID code XXXX.

***LC-MS/MS data deposition***: All targeted LC-MS/MS data from TcsD assays has been uploaded to Panorama Public[16] and is publicly available at the following link: XXXX.

***Strain availability***: All plasmids and strains generated in this work are available to the public through the Joint Bioenergy Institute's Inventory of Composable Elements: https://public-registry.jbei.org/

### Supplementary Information References
